# Antiviral Drug Discovery with an Optimized Biochemical Dengue Protease Assay: Improved Predictive Power for Antiviral Efficacy

**DOI:** 10.1101/2024.06.05.597555

**Authors:** Johannes Lang, Sudip Kumar Dutta, Lisa Reichert, Nikos Kühl, Byron Martina, Christian D. Klein

## Abstract

The viral NS2B-NS3 protease is a promising drug target to combat dengue virus (DENV) and other emerging flaviviruses. The discovery of novel DENV protease inhibitors with antiviral efficacy is hampered by the low predictive power of biochemical assays. We herein present a comparative evaluation of biochemical DENV protease assay conditions and their benchmarking against antiviral efficacy and a protease-specific reporter gene assay. Variations were performed with respect to pH, type of detergent, buffer, and substrate. The revised assay conditions were applied in a medicinal chemistry effort aimed at phenylglycine protease inhibitors. This validation study demonstrated a considerably improved predictive power for antiviral efficacy in comparison to previous approaches. An extensive evaluation of phenylglycine-based DENV protease inhibitors with highly diverse *N*-terminal caps indicates further development potential in this structural region. Furthermore, the phenylglycine moiety may be less essential than previously assumed, providing a development option towards reduced lipophilicity and thereby an improved pharmacokinetic and toxicity profile.

**Graphical Abstract:** 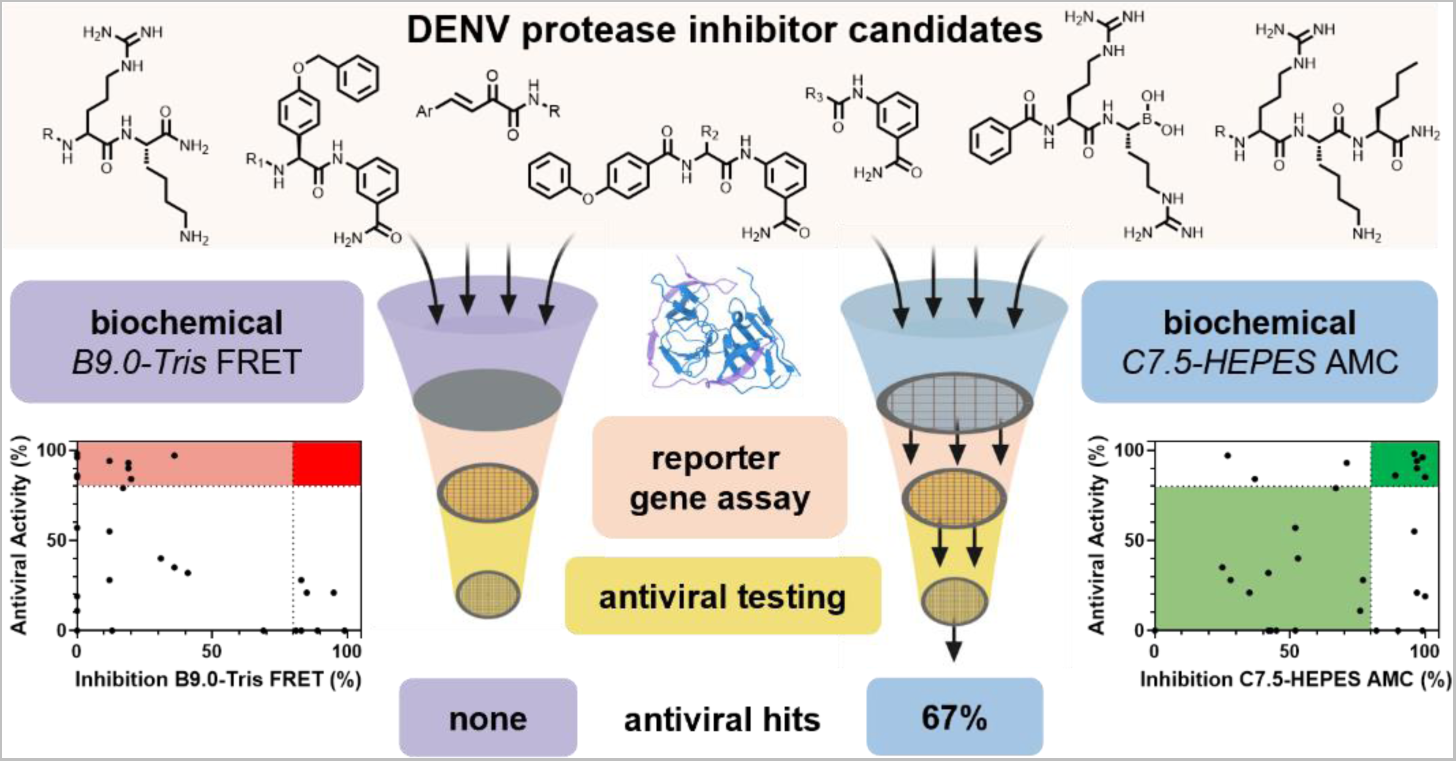

## 1. Introduction

Dengue virus (DENV) is one of the most challenging threats to global health. The world-wide incidence of this mosquito-borne viral infection has grown rapidly and is supposed to further gain momentum by factors such as spreading of transmission vectors and urbanization.^1^ Tetravalent dengue vaccines have shown efficacy in clinical trials^2, 3^ and were approved by multiple countries. Nevertheless, antiviral drugs are urgently needed against dengue and other medically relevant flaviviruses, such as West Nile and Zika virus, for which no FDA-approved vaccines exist. Viral enzymes and in particular the flaviviral NS2B-NS3 protease are promising targets for the development of specific antiviral compounds.^4^ This is emphasised by the success of approved drugs targeting the proteases of other viruses such as human immunodeficiency virus (HIV) and severe acute respiratory syndrome coronavirus 2 (SARS-CoV-2). Recently published inhibitors for flaviviral NS2B-NS3 proteases include macrocyclic substrate analogues^5^, piperazine derivatives^6^ as well as carbazole derivatives^7^ whose amidines can be masked as amidoxime prodrugs.^6, 8^ The latter carbazoles have been further developed into indolazepinone derivatives that showed promising activity against all DENV serotypes and Zika virus.^9^ Besides, activated esters as covalent inhibitors^10^ and benzothiazoles as allosteric inhibitors^11^ were published.

We previously reported phenylglycine derivatives as inhibitors of the DENV NS2B-NS3 protease.^12, 13^ They had considerably higher activity in a cell-based dengue protease reporter gene assay (DENV2proHeLa) than earlier, covalent and non-covalent, peptidic inhibitors.^14, 15^ This DENV2proHeLa reporter gene assay implements full-length NS3 with its cofactor NS2B without an artificial linker in a native biological, intracellular environment and thereby avoids potential artifacts that may arise in biochemical protease assays.^8, 16, 17^ Furthermore, the DENV2proHeLa integrates pharmacokinetic parameters like membrane permeation and – to some degree – metabolic stability into early compound characterisation.

Early-stage testing of potential antiviral protease inhibitors mainly relies on evaluation of inhibitory activity against the isolated NS2B-NS3 protease. Several DENV protease constructs have been published.^18^ One of the most commonly used constructs is constituted of the protease domain of NS3 covalently linked *via* a G_4_SG_4_ linker to the hydrophilic 40-residue core domain of NS2B (NS2B(H)-NS3pro).^19^ This construct was found to have highest catalytic activity at around pH 9. Therefore, many drug discovery campaigns relying on this construct have used pH values between 8.5 and 9.0.^5, 7, 20, 21^ While the G_4_SG_4_ linker enabled reproducible assay results it was also found to cause a different conformation of the protease^22^, restrict access to the active site for longer (hexapeptidic) ligands^23^ and limit catalytic efficiency of the protease^24^. Besides, it does not cover *cis*-cleavage of NS3 which was proposed to leave a gap in the development of anti-DENV compounds.^8^ Recent work from our group was therefore aimed at fundamental aspects of the substrate processing and ligand binding properties of DENV protease, with a particular emphasis on induced-fit vs. conformational selection mechanisms (Behnam & Klein, in revision, preprint available^25^). One result of this work was the identification of a bithiophene sensor tag to study ligand binding, irrespective of catalytic processing, to DENV protease under *in vitro* conditions.^26^

Discrepancies between biochemical and cellular/antiviral activities can be expected and explained for highly polar and reactive DENV protease inhibitors, such as the peptide-boronic acids which we published previously.^14^ In the course of our work, however, the matter became increasingly concerning when we progressed towards the preclinical development of drug-like, non-basic phenylglycine derivatives.^13^ In effect, the biochemical assay became useless to predict the antiviral efficacy of this compound class. In contrast, the DENV2proHeLa reporter gene assay remained valuable and provided clear indication that the compounds under study inhibited the DENV protease under intracellular conditions. For example, **1** (see *Figure 2*) had an EC_50_ of 0.7 µM in a DENV titer reduction assay. This value is consistent with the DENV2proHeLa (EC_50_ 0.7 µM). In stark contrast, **1** only gave 19% inhibition at 50 µM in the standard biochemical DENV protease assay.^13^ The inhibitory activity of **1** in the isolated protease assay could be partially restored by changing the detergent from Brij 58 to CHAPS, but this effect was restricted to a subset of the compounds.^13^ Similar observations were made by other researchers, and some examples shall be outlined here:

Swarbrick *et al*. showed their compounds’ influence on autocleavage of the protease. This autoprocessing cannot be performed by the NS2B(H)-NS3pro construct in the standard isolated protease biochemical assays. This could explain the more than 10-fold lower half-maximal inhibitory concentration in an antiviral assay compared to the biochemical assay.^8^

Wu *et al.* likewise reported a low correlation between inhibition of the isolated protease and antiviral activity for their set of non-competitive protease inhibitors.^27^ In contrast, protease inhibition in cell culture was correlated to antiviral activity. The authors hypothesised that a low degree of protease inhibition could lead to strong antiviral activity due to interference of polypeptide processing by intermediate processing products. This had been reported before for HIV-1.^28^

Yang *et al.* performed a high-throughput screen in a biochemical assay with more than 40,000 compounds which led to only three active compounds.^29^ Revealingly, the antiviral EC_50_ of the lead compound was 0.17 µM, around 100-fold lower compared to its biochemical IC_50_ of 15 µM.

In summary, the capability of previously used biochemical assay procedures for flaviviral proteases in high-throughput screens, hit-to-lead and lead optimization efforts to reliably predict antiviral efficacy is doubtful. Nevertheless, the resource requirements and safety restrictions of cellular, and especially antiviral assays are barely applicable in early-stage drug discovery programs. It is also highly desirable to use a clearly defined and target-specific assay to obtain inhibitors with sufficient selectivity for the viral protease. Thus, we initiated a research project to amend the conditions used in the isolated DENV protease biochemical assay in order to re-establish it as a useful tool with predictive power for functional antiviral activity.

## 2. Results and Discussion

### 2.1 Editing the Conditions of the Biochemical DENV Protease Assay

Please notice the subsequent use of a shorthand notation for assay conditions, such as *B7.5-Tris* FRET. In this example, “B” indicates the Brij detergent, used at pH 7.5, Tris buffer, and a FRET substrate.

Our standard biochemical DENV protease assay uses the NS2B(H)-NS3pro construct at a pH of 9.0.^30^ To move the pH value closer to *in vivo* conditions, various buffer components were evaluated at pH 7.5 (see *Figure 1*A). This pH value had successfully been used e.g. by Lin *et al*.^10^ To evaluate the cleavage efficiency of the substrate at the given conditions, the increase of the fluorescence signal over 16 min after addition of 50 µM substrate was monitored and the respective slope calculated in the range of 3 min to 13 min. No cleavage of our standard FRET substrate (2-Abz-nKRRS-(3-NO_2_)-Y-NH_2_) could be observed at the tested conditions (*B7.5- Tris* FRET, see *Figure 1*B). In contrast to that, efficient cleavage of the 4-aminomethylcoumarin (AMC) substrate Bz-nKRR-AMC, which had been published before^31^, was observed (*B7.5-Tris* AMC conditions). It showed a slope of 4534 RFU/min (see *Figure 1*E).

**Figure 1.**
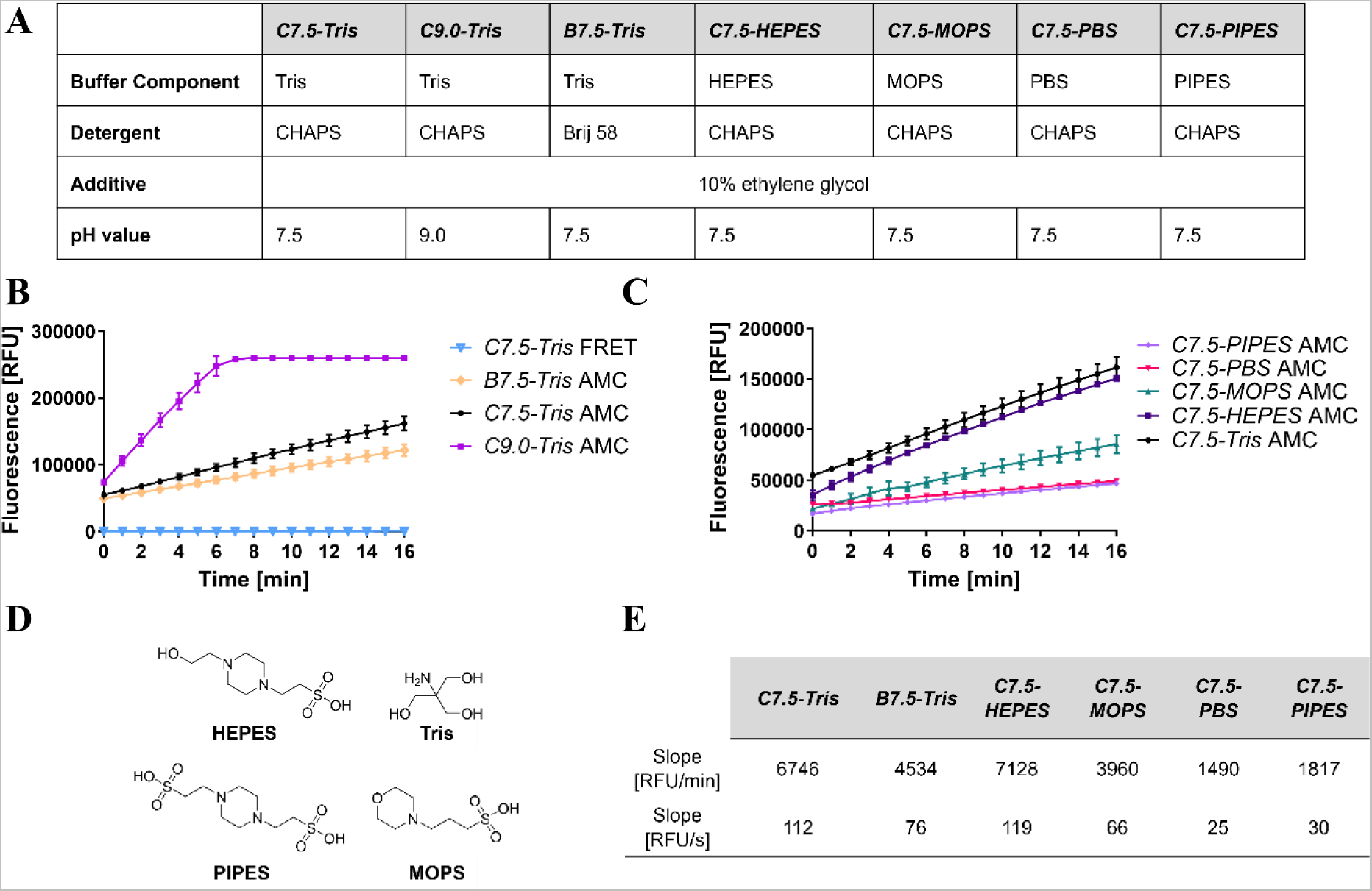
Revision of the buffer used in the biochemical DENV-2 protease assay. **(A)** Table of buffers used in the evaluation indicating their respective buffer substance, detergent, additive and pH value. **(B)** Comparison of fluorescence signal slopes using different substrates, detergents, and pH values. The diverging excitation and emission wavelengths for the respective substrates are provided in the *Supporting Information*. **(C)** Comparison of the *C7.5* AMC conditions using different buffer substances. **(D)** Structural formulae of buffer substances used in this evaluation. **(E)** Table showing respective calculated slopes of different biochemical assay conditions per minute and per second.

**Figure 2.**
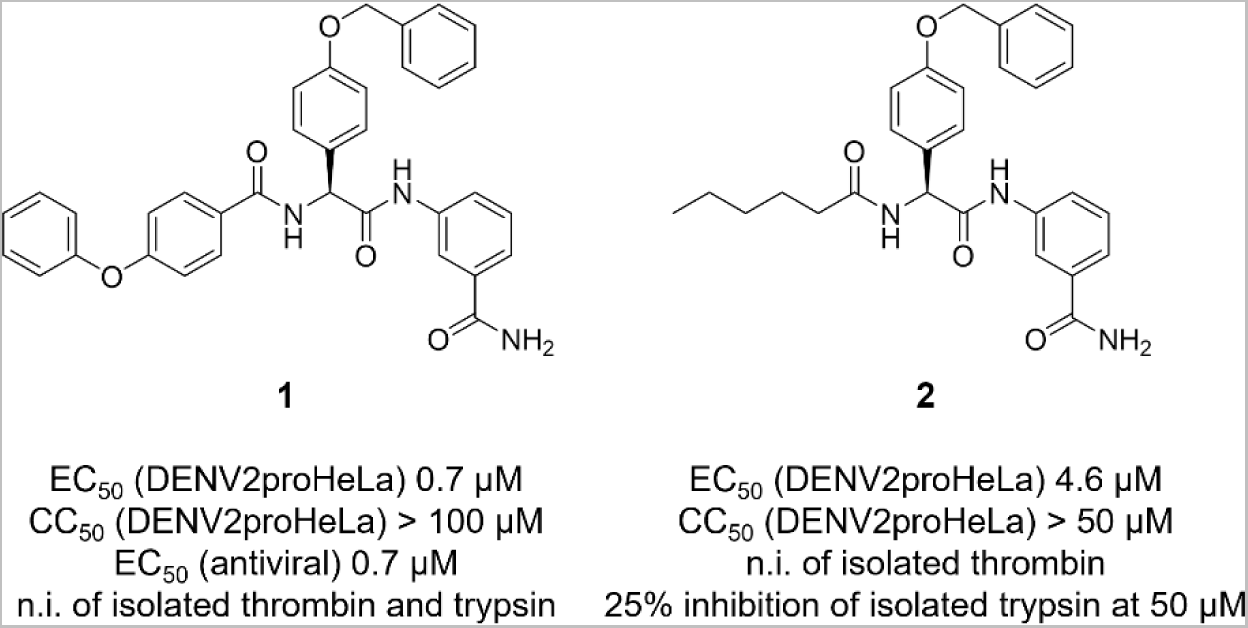
Reference compounds of this work.^13^

Early investigations on the effect of detergents on flaviviral protease activity have led to preferred use of Brij and CHAPS over Triton X-100 as detergent component in the biochemical assay.^32^ Influence on the fluorometric readout by detergents could also be shown for different proteases such as cathepsins.^33^ Therefore, the zwitterionic CHAPS and the non-ionic Brij 58 were compared in this setup. The substrate cleavage was more efficient using CHAPS (*C7.5- Tris* AMC, slope: 6746 RFU/min) as a detergent compared to Brij 58 (*B7.5-Tris* AMC, slope: 4534 RFU/min, see *Figure 1*B). This again emphasises the detergent’s influence on protease activity. Combined with previous findings that the use of CHAPS rather than Brij 58 led to improved antiviral predictability^13^, the former was chosen as detergent of choice in the following experiments.

Next, the pH value was changed back to 9.0 (*C9.0-Tris* AMC). The fluorescence signal then reached the detector maximum after 6 minutes (see *Figure 1*B). These findings confirmed the assumption that the substrate would be cleaved much more efficiently at higher pH values.^19^ Still, Bz-nKRR-AMC can provide satisfying results at a pH value of 7.5.

Further, substrate cleavage using different buffer substances was evaluated. The respective structural formulae can be found in *Figure 1*D. The raw data and the respective slopes can be seen in *Figures 1*C and *1*E. Phosphate-buffered saline (PBS) and piperazine-*N*,*N*′-bis(2- ethanesulfonic acid) (PIPES) resulted in unsatisfactory substrate processing. For PBS, this was not surprising, since inorganic salts were previously shown to decrease protease activity.^19^ The preferred buffering range of PIPES is below the pH value of 7.5 used in this experiment. 3-(*N*- Morpholino)propanesulfonic acid (MOPS) led to increased protease activity. The highest activity was observed with the established tris(hydroxymethyl)aminomethane (Tris) and 4-(2- hydroxyethyl)-1-piperazine-ethanesulfonic acid (HEPES). These are commonly used buffer substances at *in vivo* pH ranges. Since HEPES gave a higher slope with 7128 RFU/min, the *C7.5-HEPES* AMC conditions were chosen for further evaluation.

### 2.2 Evaluating the Predictive Power of *C7.5-HEPES* AMC conditions

To evaluate whether the *C7.5-HEPES* AMC conditions have a higher predictive power for the compound potencies in the cellular gene reporter assay DENV2proHeLa, phenylglycine derivatives were used as model compounds. In this class, we observed a particularly notable deviation between activities in biochemical vs. cellular assays. **1**^13^, with a 4-phenoxybenzoic acid substituent, and **2**^13^, with a hexanoic acid substituent, served as lead structures. The compounds have EC_50_ values of 0.7 µM and 4.6 µM in the DENV2proHeLa, respectively. **1** additionally showed antiviral activity with an EC_50_ of 0.7 µM in a DENV2 titer reduction assay.

To our delight, the reference compounds were active at the chosen *C7.5-HEPES* AMC conditions. **1** gave 100 ± 0.5% (IC_50_ 13 µM) and **2** 64 ± 5.1% inhibition at 50 µM inhibitor concentration (see *Figure 3* and *Supporting Information*). These inhibition values differ from the ones found in the DENV2proHeLa. This divergence had been found before.^12^ It can be explained by the different concentrations used in both assays and the further complexity of the cellular DENV2proHeLa. Compounds might not be able to cross the membrane or are metabolised within the cell. Besides, the DENV2proHeLa comprises a full-length, non-linked NS2B-NS3 construct unlike the biochemical assay. In agreement with the DENV2proHeLa data, the biochemical results indicate the inferiority of **2** vs. **1**. With respect to a screening use of the biochemical assay, we would now classify both **1** and **2** as “hits”, an observation that would have been impossible using the previously established *B9.0-Tris* FRET conditions.

**Figure 3.**
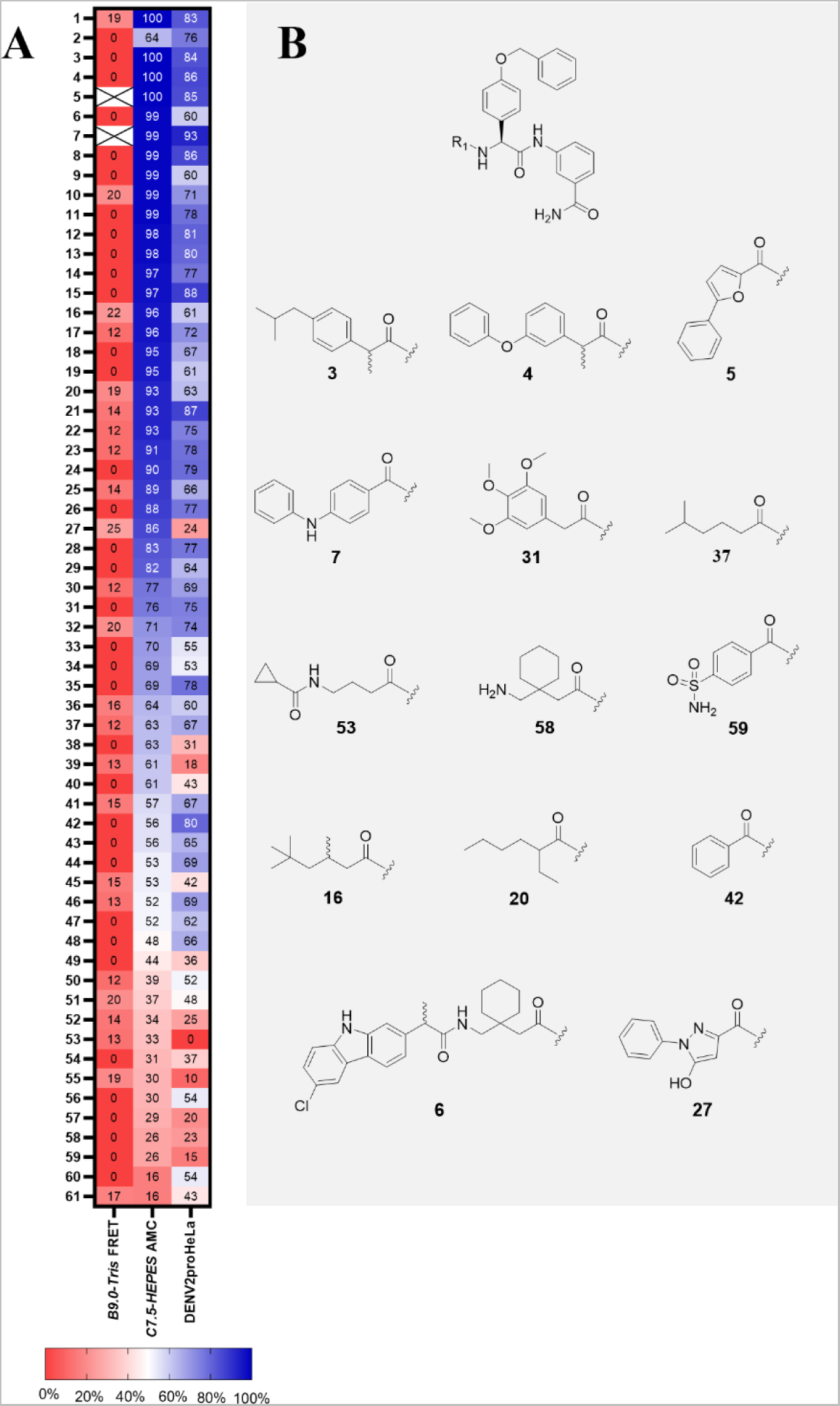
Comparison of assay results of phenylglycine derivatives with varied amino substituent. **(A)** Heatmap comparing the inhibitory activity in the biochemical assay using the *B9.0-Tris* FRET conditions (***B9.0-Tris* FRET**) and the *C7.5-HEPES* AMC conditions at 50 µM (***C7.5-HEPES* AMC**) and the DENV2proHeLa at 12.5 µM (**DENV2proHeLa**). Values in each box represent inhibitory activity in %. Apart from reference compounds **1** and **2**, values are arranged in decreasing order according to the *C7.5-HEPES* AMC assay. X = compound showed autofluorescence at the used conditions. See also *Table S1*. **(B)** Structural formulae of selected compounds. A complete set of compounds is shown in *Table S1*.

To underpin the improved predictive power of the biochemical assay, a broader set of phenylglycine derivatives was synthesised. Furthermore, it was desired to assess the impact on assay readout by certain parts of the molecules. First studies focused on the substituent on the phenylglycine’s nitrogen. The inhibition values using the *C7.5-HEPES* AMC conditions at 50 µM were compared to the ones in the DENV2proHeLa at 12.5 µM. These concentrations represent the standard screening values in our group and are routinely used to evaluate whether a compound is a promising inhibitor.^12, 13, 17^ The standard screening concentration in the DENV2proHeLa reporter gene assay is lower than in the biochemical assay mainly because of cytotoxicity considerations for “random” screening compounds. A comparison of both values as well as structures of selected compounds are given in *Figure 3*. The data presented in each box in *Figure 3*B shows the inhibition in percent in the respective assay system. Data with all structural formulae are provided in the *Supporting Information*.

The chosen set of compounds gave inhibition values in the revised biochemical assay ranging from 16% to 100%. The most active compounds at the given concentration were **3**, **4** and **5** that also represent three of the most active compounds in the DENV2proHeLa. *Vice versa*, **7**, the compound with the highest inhibition in the DENV2proHeLa, also exerted high inhibition of 99% in the biochemical assay. Moderately lower inhibition could also equally be seen in both systems, e.g. in **31** (76 and 75% inhibition, respectively) and **37** (63 and 67% inhibition, respectively). Moreover, the trend continued for compounds with low to no activity. E.g. **58** and **59** were weak inhibitors in both systems. **53**, being inactive in the DENV2proHeLa, likewise showed very low activity in the biochemical assay with only 30% inhibition.

At the same time, not all compounds with promising activity in the biochemical assay performed equally in the DENV2proHeLa. **9** and **16** showed more than 90% inhibition of the isolated protease but only around 60% inhibition in the DENV2proHeLa. **27** and **39** were three times less active in the cellular assay compared to the biochemical assay.

Nevertheless, considering that the biochemical assay is an early-stage testing system with a higher throughput, these results were very satisfactory. Assuming a “hit threshold” at 75% inhibition, with **35** and **42** only two out of 61 compounds would have been falsely eliminated before cellular testing. Other compounds that performed better in the DENV2proHeLa compared to the biochemical assay, such as **60** and **61**, would not have been considered a hit either way.

The *B9.0-Tris* FRET conditions were not suited to predict any of the hits with activity in the DENV2proHeLa cellular assay. None of the tested compounds exceeds 25% inhibition at 50 µM under these conditions. Besides, the conditions were unsuitable to determine inhibitory values for **7** and **5** as these compounds are fluorescent at the wavelengths used for the FRET substrate. This was not the case for the wavelengths used for the AMC substrate.

To further evaluate the predictability of the *C7.5-HEPES* AMC conditions, a second set of compounds was synthesised. It included derivatives of the central 4-benzyloxyphenylglycine moiety. In that way, the reliability of the novel conditions could be evaluated using chemically more diverse compounds. A comparison of inhibition values in both testing systems is shown in *Figure 4*A. **62**, **63** and **64** were the most potent compounds in the biochemical assay and were also three of the most active compounds in the DENV2proHeLa. Decreased inhibition (**75**, **76**) and inferiority of **78** were observed in both assay systems. In contrast, **68**’s activity *in vitro* was not represented in the DENV2proHeLa. Out of this set of compounds, no compound would have been falsely eliminated at a 75% inhibition threshold for a hit. Similar to the first set of compounds with divergent substituents on the phenylglycine’s nitrogen, the *C7.5-HEPES* AMC conditions have a good predictive power for the DENV2proHeLa in this set of compounds as well. The *B9.0-Tris* FRET conditions would not have identified any hits from this set.

**Figure 4.**
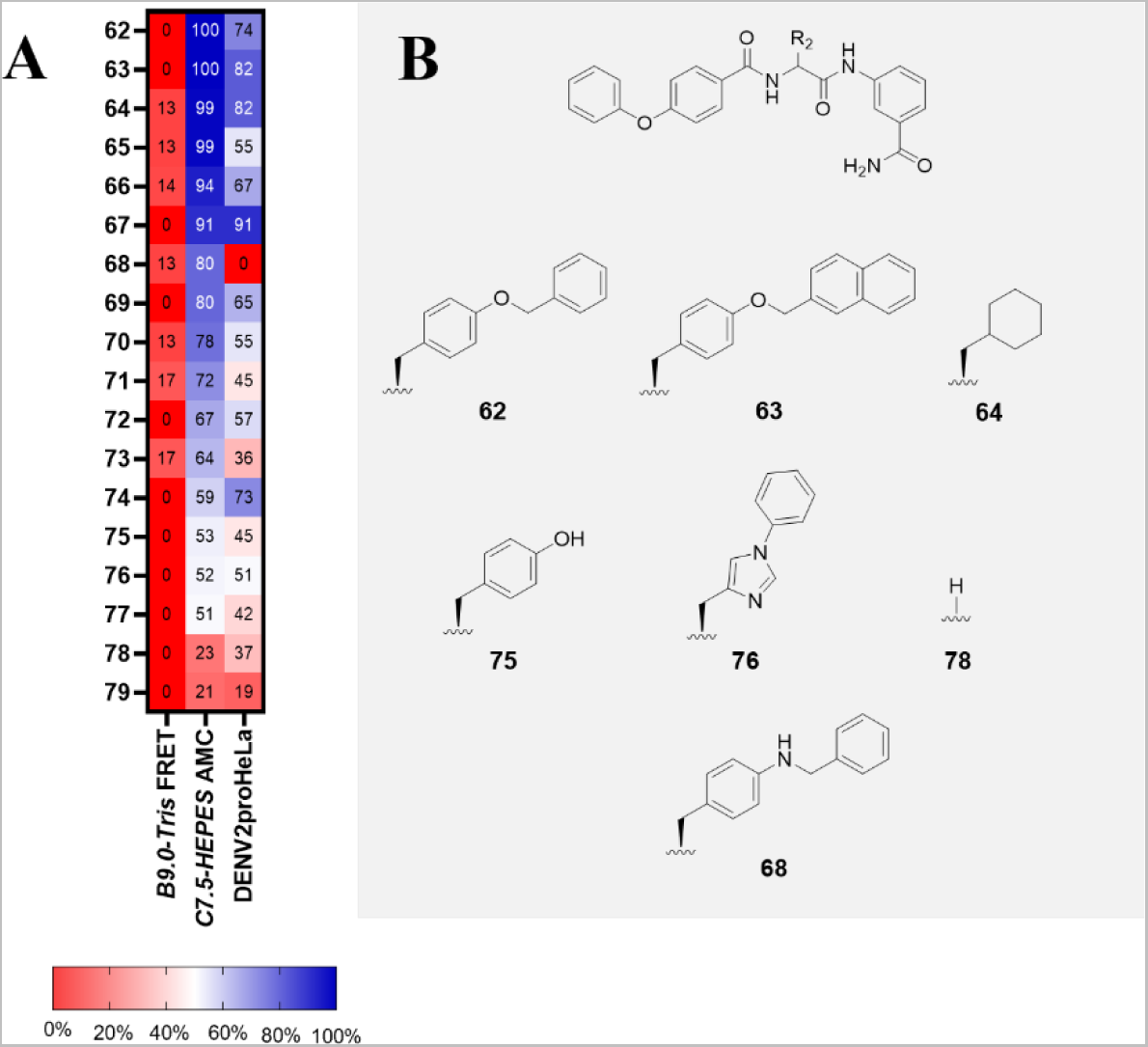
Comparison of assay results of inhibitors with a varied core amino acid. **(A)** Heatmap comparing the inhibitory activity in the biochemical assay using the *B9.0-Tris* FRET conditions (***B9.0-Tris* FRET**) and the *C7.5-HEPES* AMC conditions at 50 µM (***C7.5-HEPES* AMC**) as well as the DENV2proHeLa at 12.5 µM (**DENV2proHeLa**). Values in each box represent inhibitory activity in %. Values are arranged in a decreasing manner according to the *C7.5-HEPES* AMC assay. See also *Table S2*. **(B)** Structural formulae of selected compounds. The structural formulae of the complete set is provided in *Table S2*.

A graphical comparison of inhibitory values using the *C7.5-HEPES* AMC conditions at 50 µM and the DENV2proHeLa at 12.5 µM of all compounds is shown in *Figure 5*A. The red area contains compounds with activity above the 75% “hit threshold” under the *C7.5-HEPES* AMC conditions, the blue area indicates activity above 75% in the DENV2proHeLa. The overlapping area (grey) represents all inhibitors that were considered a hit in both testing systems, the non-coloured area represents compounds considered a hit in neither system. By extension, the latter two areas illustrate the percentage of compounds for which the *C7.5-HEPES* AMC conditions could predict cellular activity correctly: an accuracy of 82%. Only three compounds out of the full set, **2**, **35**, and **42**, were false negatives, a ratio of 4%. An analogous graphical comparison for the *B9.0-Tris* FRET conditions is shown in *Figure 5*B, where the biochemical conditions failed to predict any cellular hits. Besides, there is no obvious trend visible that potent compounds in the DENV2proHeLa tend to show relatively higher activity under the *B9.0-Tris* FRET conditions.

**Figure 5.**
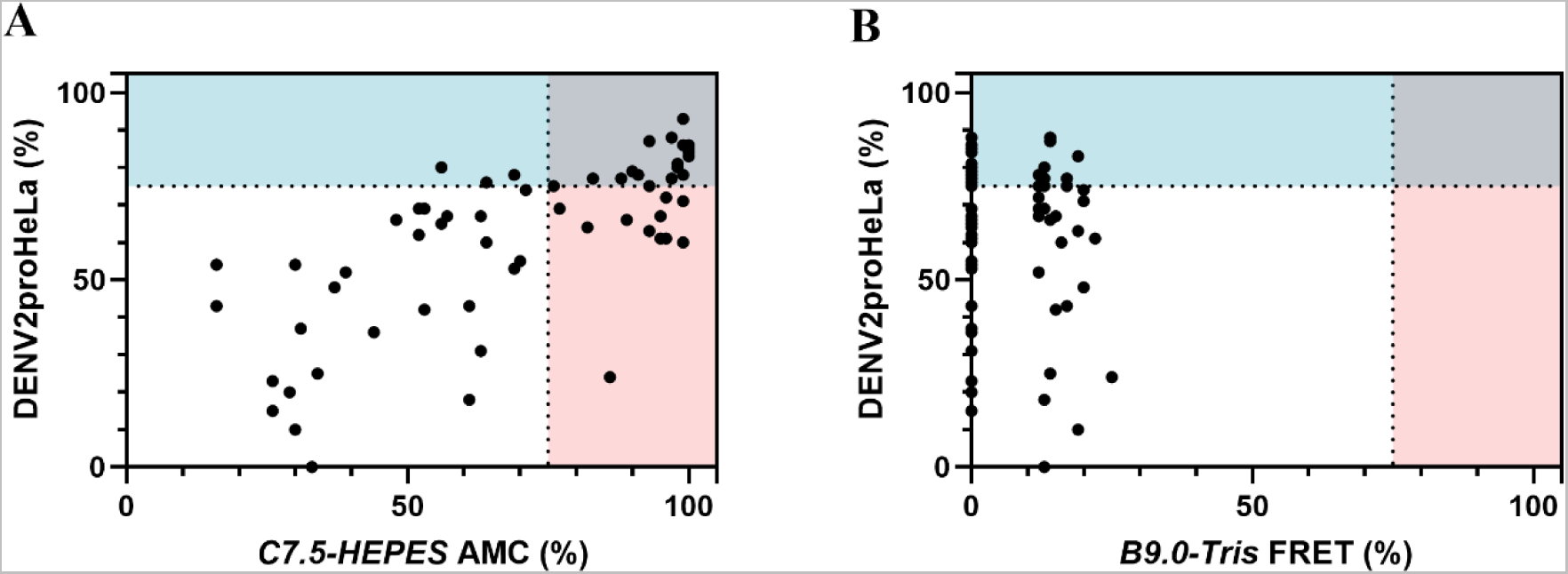
(**A**) Graphical comparison of the inhibitory values in % of all tested compounds in the DENV2proHeLa at 12.5 µM (**DENV2proHeLa (%)**) and using the *C7.5-HEPES* AMC conditions in the biochemical assay at 50 µM (***C7.5-HEPES* AMC (%)**). The thresholds (75%) are highlighted as dashed lines. (**B**) Graphical comparison of the inhibitory values in % of all tested compounds in the DENV2proHeLa at 12.5 µM (**DENV2proHeLa (%)**) and using the *B9.0-Tris* FRET conditions in the biochemical assay at 50 µM (***B9.0-Tris* FRET (%)**). The thresholds (75%) are highlighted as dashed lines.

To further quantify the relative performance of the assays, IC_50_ values in the biochemical assay and EC_50_ values in the DENV2proHeLa were determined (see *Supporting Information*). The set of compounds covered IC_50_ values ranging from 4.8 to 45 µM. Very similar half-maximal concentration values were found for **16** (IC_50_ and EC_50_ 11 µM), **21** (IC_50_ 6.6 µM, EC_50_ 6.0 µM), **63** (IC_50_ 4.8 µM, EC_50_ 5.1 µM), and **65** (IC_50_ 5.9 µM, EC_50_ 7.4 µM).

All compounds with an IC_50_ below 10 µM also had an EC_50_ below 10 µM. Additionally, 6 out of 8 compounds with an IC_50_ above 25 µM showed an EC_50_ below 10 µM. **1** and **7**, both with an EC_50_ of 0.7 µM, had higher IC_50_ values of 9.5 µM and 12.8 µM, respectively. **18** (EC_50_ 3.1 µM) and **23** (EC_50_ 2.4 µM) also had higher IC_50_ values with 16 µM and 17 µM. In the complete set of compounds, half-maximal concentration values in the DENV2proHeLa are generally lower than the biochemical ones except for **27** and **68**. This pattern was observed previously under similar biochemical conditions.^12, 13^ A possible explanation for this is the artificial constitution of the biochemical assay, including polyols^30^, choice of buffer systems, ionic strength, and artificial covalent linking within the NS2B(H)-NS3pro construct. Additionally, the intracellular environment of the DENV2proHeLa includes the effect of macromolecular crowding on protease dynamics, as shown for HIV-1 protease and SARS-CoV-1 3CL protease.^34, 35^

### 2.3 Structure-Activity Relationships

The compound set presented herein additionally gives valuable insight into the structure-activity relationship (SAR) of phenylglycine-derived DENV protease inhibitors. Variation of the aliphatic amino substituent of compound **2** with different cyclic, alkylic or halogen substituents did not lead to increase in activity in neither testing system. Yet, introduction of a bridging gabapentin could increase inhibitory potential. This increase was observed for both aliphatic and aromatic caps as can be seen in the equipotency of **12** (IC_50_ 9.2 µM, EC_50_ (DENV2proHeLa) 3.8 µM) and **24** (IC_50_ 12 µM, EC_50_ (DENV2proHeLa) 3.7 µM). However, **58** was explicitly less active (26% inhibition at 50 µM in biochemical assay, EC_50_ (DENV2proHeLa) 21 µM). It contains a gabapentin moiety with an unsubstituted nitrogen. The bulkiness of the gabapentin moiety appears to increase inhibitory potency as long as its polar amino group is substituted.

Lipophilic aromatic substituents of the phenylglycine’s nitrogen tended to be more active than aliphatic ones. **7** is the most potent compound in the DENV2proHeLa (EC_50_ 0.7 µM) and can compete with the activity of the structurally similar reference compound **1**. At the same time, exchanging one phenyl ring of **1** with a pyridine as in **33** led to a drastic activity loss in both testing systems (IC_50_ 34 µM, EC_50_ 8.8 µM). This *C*-to-*N* change seems to interfere interaction with the binding pocket. In general, the target seems to favour longer and bulkier lipophilic moieties in this position. This can be seen by the superiority of e.g. **4**, **22** or **23** to compounds with smaller residues such as **19**, **49** or **59**. The latter’s polar sulphonamide appears to have a detrimental effect on activity.

Replacing the central 4-benzyloxyphenylglycine of **1** with different natural and non-natural L-amino acids generally led to loss in activity in the DENV2proHeLa reporter gene assay. Nevertheless, especially compounds with a central aliphatic amino acid still had promising activity. The most active compound in the DENV2proHeLa is **74**, bearing a central L- cyclohexylglycine, with an EC_50_ of 3.9 µM. Even a central glycine as in **78** showed some activity in both testing systems. A sterically demanding group seems not to be essential for inhibition. This could open up new possibilities for the development of DENV protease inhibitors beyond phenylglycine derivatives.

To identify non-specific inhibitors, the inhibitory activity against thrombin and trypsin was determined. Both enzymes are serine proteases that show a similar substrate recognition compared to the DENV protease. No relevant inhibition of these off-targets was observed (see *Supporting Information*).

### 2.4 Testing in Anti-DENV Immunofluorescence-Based Assay

#### 2.4.1 Antiviral Screening

With the new biochemical conditions in hand, and having shown their capability to predict activity in the DENV2proHeLa reporter gene assay, we proceeded to evaluate their predictive power for antiviral activity. For this purpose, an even broader set of inhibitors including previously published DENV protease inhibitors with different chemical scaffolds^15, 36–38^ were tested in an immunofluorescence-based anti-DENV-2 assay. The inhibition values at 25 µM inhibitor concentration are compared to the screening results in the DENV2proHeLa as well as the previous and the novel biochemical conditions in *Figure 6*A.

**Figure 6.**
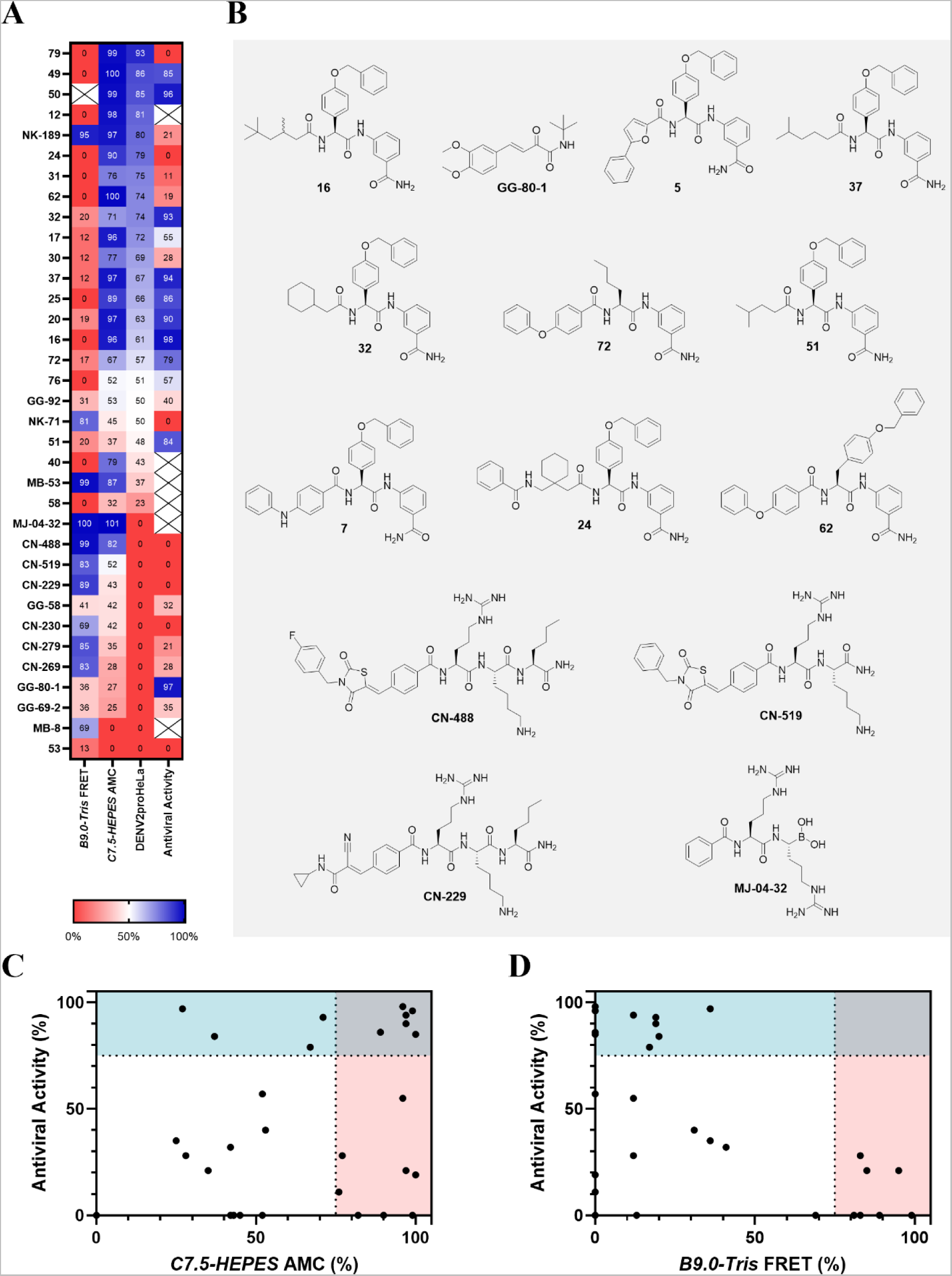
Comparison of inhibitory results in the biochemical assay, the DENV2proHeLa and immunofluorescence-based antiviral assay. **(A)** Heatmap comparing the inhibitory activity in the biochemical assay using the *B9.0-Tris* FRET (***B9.0-Tris* FRET**) and the *C7.5-HEPES* AMC (***C7.5-HEPES* AMC**) conditions at 50 µM, the DENV2proHeLa (**DENV2proHeLa**) at 12.5 µM and in the immunofluorescence-based antiviral assay (**Antiviral Activity**) at 25 µM. Values in each box represent inhibitory activity in %. Values are arranged in a decreasing order according to the *C7.5-HEPES* AMC conditions. The “X” symbol indicates: ***B9.0-Tris* FRET**, X = compound showed autofluorescence at the used conditions. **Antiviral Activity**, X = compound led to increase of viral load. See also *Table S3*. **(B)** Structural formulae of selected inhibitors. **(C)** Plot of inhibitory values in % in the antiviral testing at 25 µM (**Antiviral Inhibition (%)**) and the biochemical *C7.5-HEPES* AMC conditions at 50 µM (***C7.5-HEPES* AMC (%)**). The thresholds (75%) are highlighted as dashed lines. **(D)** Plot of inhibitory values in % in the antiviral testing at 25 µM (**Antiviral Activity (%)**) and the biochemical B*9.0-Tris* FRET conditions at 50 µM (***B9.0-Tris* FRET (%)**). The thresholds (75%) are highlighted as dashed lines.

The most active compounds in the antiviral assay at the tested concentration were **16** (98% inhibition), **GG-80-1** (97% inhibition), **5** (96% inhibition), **37** (94% inhibition) and **32** (93% inhibition). Apart from **GG-80-1**, all these compounds showed more than 60% inhibition in the other upstream testing setups.

On the other hand, several hits from the biochemical and reporter gene assay had low activity in the antiviral assay. In particular, **7**, **24** and **62**, some of the most potent compounds in both upstream testing setups, failed to show significant antiviral activity. It can be hypothesized that this is due to factors that are neither covered by the biochemical assay nor the DENV2proHeLa, e.g. intracellular competition with substrates within the infected cells, interaction with different viral proteins and presence of viral replication complexes.^39^ These complexes consist of eight non-structural viral proteins, including NS2B and NS3. This complexity and its influence on NS2B-NS3 protease activity and recognition is not reflected in any of the upstream assay systems.

**CN-229**, **CN-488** and **CN-519** were inactive in the cellular assays despite having some moderate activity in the biochemical assay. Their inactivity under cellular conditions can likely be attributed to insufficient membrane permeability (polar sidechains, polar moieties at the *N*- termini). The peptide-boronic acid **MJ-04-32**, a covalent-reversible electrophilic inhibitor that binds to the catalytic serine, was the most active compound in the biochemical assay (IC_50_ 0.068 µM, see *Supporting Information*) but inactive in the DENV2proHeLa and even increased viral load in the antiviral assay. This is in line with previous observations^14, 37^ and can be attributed to insufficient membrane permeability of this highly charged and polar compound, along with non-specific binding of the boronic acid moiety to a variety of glycans in the cell culture medium and cellular structures.^40^ However, peptide-boronic acids like **MJ-04-32** are extremely valuable as reference inhibitors and site-specific labels for biochemical assays, mechanistic and structural studies, and have recently become easily accessible by novel chemical methodologies.^14, 41, 42^

Four compounds with antiviral activity, **32**, **51**, **72** and **GG-80-1**, were not identified as protease inhibitors in the biochemical assay (see *Figure 6*C). Of these, **72** can be considered a borderline case, being positioned near the threshold in both the biochemical and the antiviral assays (67% inhibition at 50 µM vs. 79% reduction at 25 µM). The antiviral activity of **GG-80-1** (97% reduction at 25 µM) is very likely due to non-specific effects of the highly reactive β,γ- unsaturated α-ketoamide structure.

The B*9.0-Tris* FRET conditions were not capable to predict any of the antiviral hits and, *vice versa*, all of the nine biochemical hits were inactive in the antiviral testing (see *Figure 6*D).

These results underpin the superiority of the novel *C7.5-HEPES* AMC conditions. They had a sensitivity of 60% (threshold: 75%) resp. 67% (threshold: 80%) for antiviral activity (see *Figure 6*C). More importantly, they had a negative predictive value of 73% resp. 82%. In comparison, the previously used *B9.0-Tris* FRET predicted 0% (both thresholds) of the antiviral hits and the negative predictive value was only 55% resp. 60% (see *Figure 6*D). Additionally, none of the *B9.0-Tris* FRET hits could be confirmed in antiviral testing whereas the *C7.5-HEPES* AMC conditions had a precision of 50%.

#### 2.4.2 Antiviral EC_50_ Determination

Antiviral dose-response curves were determined for selected compounds (see *Figure 7*). All compounds showed dose-dependent reduction of viral load. EC_50_ values ranged from 2.8 to 20 µM. **4** with a 4-phenoxybenzylacetic acid substituent on the phenylglycine’s nitrogen had the lowest EC_50_ value. **5**, **16**, **17** and **32** gave around 2.5-fold higher values. **51**, with a 4-methyl substituted pentanoic acid cap gave the highest EC_50_ value, almost 10-fold higher than **4**. Albeit less active than previously published phenylglycine derivatives^12, 13^, these results again confirm the antiviral potency of this inhibitor class. Besides, the data proves that no bulky aromatic substituent on the phenylglycine’s nitrogen is required, as shown by **16**. This offers potential for the future development of this compound class.

**Figure 7.**
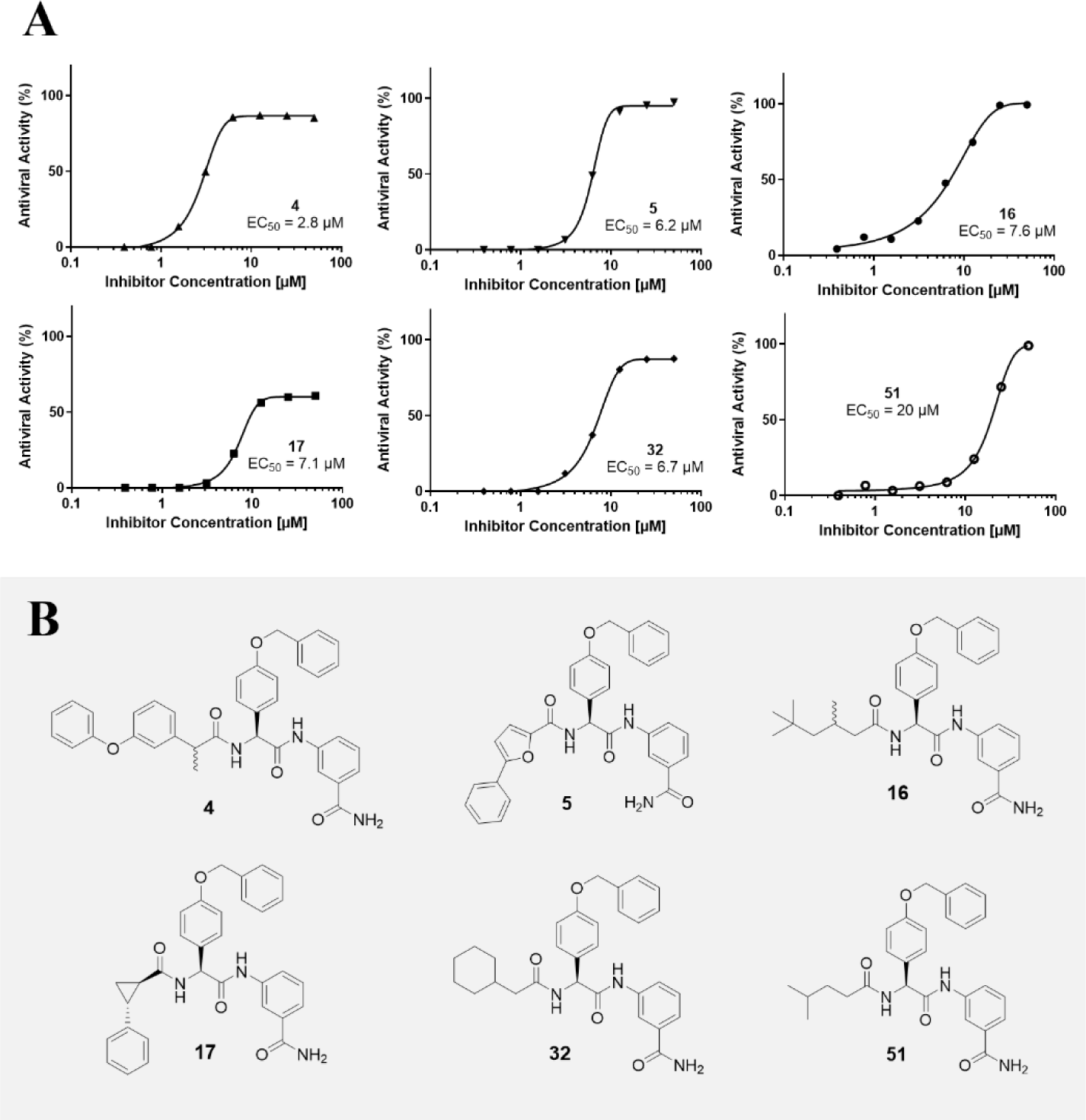
Anti-DENV-2 EC_50_ determination of selected compounds. **(A)** Dose-response curves of selected compounds. Determined EC_50_ values are given in the graph. **(B)** Structural formulae of evaluated compounds.

## 3. Conclusions

We herein demonstrate that the *C7.5-HEPES* AMC biochemical assay conditions have a considerably higher predictive power for cellular protease and functional antiviral activity compared to the commonly used *B9.0-Tris* FRET conditions. This finding was substantiated using a broad set of chemically diverse compounds. We find it particularly important for an application in drug discovery and development that the *C7.5-HEPES* AMC conditions are capable to identify and characterize DENV protease inhibitors with a less polar and more drug-like profile, such as the phenylglycines. Relative to the DENV2proHeLa reporter gene assay, the *C7.5-HEPES* AMC conditions had a negative predictive value of 90%. At the same time, the sensitivity was 60%. Even though the latter value is comparably lower, we interpret these results to be very satisfying especially with the *B9.0-Tris* FRET conditions being unable to predict any hit. Considering the biochemical assay as an early-stage high-throughput system, it is of higher value to use a testing setup that reliably eliminates compounds rather than one that reliably predicts true hits but simultaneously expels promising candidates. Therefore, one is willing to include additional compounds that are eliminated in later, more complex testing setups.

Considering antiviral activity, the sensitivity could be improved from 0% (*B9.0-Tris* FRET, both thresholds) to 60% resp. 67% (*C7.5-HEPES* AMC, 75% resp. 80% threshold). At the same time, the *C7.5-HEPES* AMC conditions had a negative predictive value of 73% resp. 82% (*B9.0-Tris* FRET: 55% resp. 60%) and a precision of 43% resp. 50% (*B9.0-Tris* FRET: 0%). Thus, the superior predictive power of the *C7.5-HEPES* AMC conditions can also be applied to antiviral testing.

The reasons for the drastically different results using different conditions can only be speculated. It was found before that biochemical protease assay readouts are influenced by several factors such as detergents,^16, 32^ salt and polyol concentrations as well as pH value.^19^ NMR studies could show that the interaction of NS3 and NS2B is influenced by salt concentration and pH value.^43, 44^ A proposed ‘closed’ conformation was destabilised with increasing pH value.^43^ Therefore, we hypothesise that the high pH value of *B9.0-Tris* FRET conditions leads to a conformation of the NS2B(H)-NS3pro construct that does not reflect the true intracellular behaviour of the NS2B-NS3 protease.

In addition, new insights were gained into the antiviral potency of the phenylglycines. Aliphatic carboxylic acid substituents on phenylglycine’s nitrogen showed comparable activity to aromatic ones. A central 4-benzyloxyphenylglycine moiety is not essential to achieve antiviral activity. In this way, new SAR study options arise that could lead to smaller and less lipophilic inhibitors of the DENV protease.

We consider the *C7.5-HEPES* AMC conditions to be a major improvement. Reliability of early-stage high-throughput screenings using the biochemical assay will be eminently increased. Besides, more demanding cellular and/or antiviral testing can be circumvented early in the drug discovery process. These findings could lead to a paradigm change in early-stage flaviviral protease inhibitor development.

## Acknowledgements

We thank Heiko Rudy for measuring ESI high resolution spectra, Natascha Stefan and Tobias Timmermann for technical support as well as Dr. Stefan Hinkes, Luisa Burger, Judith Stirn, Mareike Thimig and Svea Wupper for synthetic support. Leah Glanzmann, Judith Stirn and Marius Werner provided valuable suggestions during the preparation of the manuscript. The antiviral assay was funded by a Eurostars E! 12504 -DRIVE grant. This project was supported by the Volkswagen Foundation under grant number 9A836 (‘Preclinical development of antiviral protease inhibitors targeting flavi- and coronaviruses’).

## References

(1) Messina, J. P.; Brady, O. J.; Golding, N.; Kraemer, M. U. G.; Wint, G. R. W.; Ray, S. E.; Pigott, D. M.; Shearer, F. M.; Johnson, K.; Earl, L.;, et al. The current and future global distribution and population at risk of dengue. Nature Microbiology 2019, 4 (9), 1508–1515. DOI: 10.1038/s41564-019-0476-8.

(2) Biswal, S.; Borja-Tabora, C.; Martinez Vargas, L.; Velásquez, H.; Theresa Alera, M.; Sierra, V.; Johana Rodriguez-Arenales, E.; Yu, D.; Wickramasinghe, V. P.; Duarte Moreira, E.;, et al. Efficacy of a tetravalent dengue vaccine in healthy children aged 4–16 years: a randomised, placebo-controlled, phase 3 trial. The Lancet 2020, 395 (10234), 1423–1433. DOI: 10.1016/S0140-6736(20)30414-1.

(3) Thoresen, D.; Matsuda, K.; Urakami, A.; Ngwe Tun Mya, M.; Nomura, T.; Moi Meng, L.; Watanabe, Y.; Ishikawa, M.; Hau Trang Thi, T.; Yamamoto, H.;, et al. A tetravalent dengue virus-like particle vaccine induces high levels of neutralizing antibodies and reduces dengue replication in non-human primates. Journal of Virology 2024, 0 (0), e00239–00224. DOI: 10.1128/jvi.00239-24 (acccessed 2024/05/02).

(4) Behnam, M. A. M.; Nitsche, C.; Boldescu, V.; Klein, C. D. The Medicinal Chemistry of Dengue Virus. Journal of Medicinal Chemistry 2016, 59 (12), 5622–5649. DOI: 10.1021/acs.jmedchem.5b01653.

(5) Braun, N. J.; Quek, J. P.; Huber, S.; Kouretova, J.; Rogge, D.; Lang-Henkel, H.; Cheong, E. Z. K.; Chew, B. L. A.; Heine, A.; Luo, D.;, et al. Structure-Based Macrocyclization of Substrate Analogue NS2B-NS3 Protease Inhibitors of Zika, West Nile and Dengue viruses. ChemMedChem 2020, 15 (15), 1439–1452. DOI: 10.1002/cmdc.202000237 (acccessed 2024/05/21).

(6) del Rosario García-Lozano, M.; Dragoni, F.; Gallego, P.; Mazzotta, S.; López-Gómez, A.; Boccuto, A.; Martínez-Cortés, C.; Rodríguez-Martínez, A.; Pérez-Sánchez, H.; Manuel Vega-Pérez, J.;, et al. Piperazine-derived small molecules as potential Flaviviridae NS3 protease inhibitors. In vitro antiviral activity evaluation against Zika and Dengue viruses. Bioorganic Chemistry 2023, 133, 106408. DOI: 10.1016/j.bioorg.2023.106408.

(7) Rassias, G.; Zogali, V.; Swarbrick, C. M. D.; Ki Chan, K. W.; Chan, S. A.; Gwee, C. P.; Wang, S.; Kaplanai, E.; Canko, A.; Kiousis, D.;, et al. Cell-active carbazole derivatives as inhibitors of the zika virus protease. European Journal of Medicinal Chemistry 2019, 180, 536–545. DOI: 10.1016/j.ejmech.2019.07.007.

(8) Swarbrick, C.; Zogali, V.; Chan, K. W. K.; Kiousis, D.; Gwee, C. P.; Wang, S.; Lescar, J.; Luo, D.; von Itzstein, M.; Matsoukas, M.-T.;, et al. Amidoxime prodrugs convert to potent cell-active multimodal inhibitors of the dengue virus protease. European Journal of Medicinal Chemistry 2021, 224, 113695. DOI: 10.1016/j.ejmech.2021.113695.

(9) Zogali, V.; Kiousis, D.; Voutyra, S.; Kalyva, G.; Abdul Mahid, M. B.; Bist, P.; Ki Chan, K. W.; Vasudevan, S. G.; Rassias, G. Carbazole to indolazepinone scaffold morphing leads to potent cell-active dengue antivirals. European Journal of Medicinal Chemistry 2024, 268, 116213. DOI: 10.1016/j.ejmech.2024.116213.

(10) Lin, X.; Cheng, J.; Wu, Y.; Zhang, Y.; Jiang, H.; Wang, J.; Wang, X.; Cheng, M. Identification and In Silico Binding Study of a Highly Potent DENV NS2B-NS3 Covalent Inhibitor. ACS Medicinal Chemistry Letters 2022, 13 (4), 599–607. DOI: 10.1021/acsmedchemlett.1c00653.

(11) Maus, H.; Barthels, F.; Hammerschmidt, S. J.; Kopp, K.; Millies, B.; Gellert, A.; Ruggieri, A.; Schirmeister, T. SAR of novel benzothiazoles targeting an allosteric pocket of DENV and ZIKV NS2B/NS3 proteases. Bioorganic & Medicinal Chemistry 2021, 47, 116392. DOI: 10.1016/j.bmc.2021.116392.

(12) Kühl, N.; Graf, D.; Bock, J.; Behnam, M. A. M.; Leuthold, M.-M.; Klein, C. D. A New Class of Dengue and West Nile Virus Protease Inhibitors with Submicromolar Activity in Reporter Gene DENV-2 Protease and Viral Replication Assays. Journal of Medicinal Chemistry 2020, 63 (15), 8179–8197. DOI: 10.1021/acs.jmedchem.0c00413.

(13) Kühl, N.; Leuthold, M. M.; Behnam, M. A. M.; Klein, C. D. Beyond Basicity: Discovery of Nonbasic DENV-2 Protease Inhibitors with Potent Activity in Cell Culture. Journal of Medicinal Chemistry 2021, 64 (8), 4567–4587. DOI: 10.1021/acs.jmedchem.0c02042.

(14) Nitsche, C.; Zhang, L.; Weigel, L. F.; Schilz, J.; Graf, D.; Bartenschlager, R.; Hilgenfeld, R.; Klein, C. D. Peptide–Boronic Acid Inhibitors of Flaviviral Proteases: Medicinal Chemistry and Structural Biology. Journal of Medicinal Chemistry 2017, 60 (1), 511–516. DOI: 10.1021/acs.jmedchem.6b01021.

(15) Behnam, M. A. M.; Graf, D.; Bartenschlager, R.; Zlotos, D. P.; Klein, C. D. Discovery of Nanomolar Dengue and West Nile Virus Protease Inhibitors Containing a 4-Benzyloxyphenylglycine Residue. Journal of Medicinal Chemistry 2015, 58 (23), 9354–9370. DOI: 10.1021/acs.jmedchem.5b01441.

(16) Ehlert, F. G. R.; Linde, K.; Diederich, W. E. What Are We Missing? The Detergent Triton X-100 Added to Avoid Compound Aggregation Can Affect Assay Results in an Unpredictable Manner. 2017, 12 (17), 1419–1423. DOI: 10.1002/cmdc.201700329.

(17) Kühl, N.; Lang, J.; Leuthold, M. M.; Klein, C. D. Discovery of potent benzoxaborole inhibitors against SARS-CoV-2 main and dengue virus proteases. European Journal of Medicinal Chemistry 2022, 240, 114585. DOI: 10.1016/j.ejmech.2022.114585.

(18) Nitsche, C.; Holloway, S.; Schirmeister, T.; Klein, C. D. Biochemistry and Medicinal Chemistry of the Dengue Virus Protease. Chemical Reviews 2014, 114 (22), 11348–11381. DOI: 10.1021/cr500233q.

(19) Leung, D.; Schroder, K.; White, H.; Fang, N.-X.; Stoermer, M. J.; Abbenante, G.; Martin, J. L.; Young, P. R.; Fairlie, D. P. Activity of Recombinant Dengue 2 Virus NS3 Protease in the Presence of a Truncated NS2B Co-factor, Small Peptide Substrates, and Inhibitors *. Journal of Biological Chemistry 2001, 276 (49), 45762–45771. DOI: 10.1074/jbc.M107360200 (acccessed 2022/10/27).

(20) Maus, H.; Gellert, A.; Englert, O. R.; Chen, J.-X.; Schirmeister, T.; Barthels, F. Designing photoaffinity tool compounds for the investigation of the DENV NS2B–NS3 protease allosteric binding pocket. RSC Medicinal Chemistry 2023, 14 (11), 2365–2379, 10.1039/D3MD00331K. DOI: 10.1039/D3MD00331K.

(21) Phoo, W. W.; El Sahili, A.; Zhang, Z.; Chen, M. W.; Liew, C. W.; Lescar, J.; Vasudevan, S. G.; Luo, D. Crystal structures of full length DENV4 NS2B-NS3 reveal the dynamic interaction between NS2B and NS3. Antiviral Research 2020, 182, 104900. DOI: 10.1016/j.antiviral.2020.104900.

(22) Hill, M. E.; Yildiz, M.; Hardy, J. A. Cysteine Disulfide Traps Reveal Distinct Conformational Ensembles in Dengue Virus NS2B-NS3 Protease. Biochemistry 2019, 58 (6), 776–787. DOI: 10.1021/acs.biochem.8b00978.

(23) Shannon, A. E.; Chappell, K. J.; Stoermer, M. J.; Chow, S. Y.; Kok, W. M.; Fairlie, D. P.; Young, P. R. Simultaneous uncoupled expression and purification of the Dengue virus NS3 protease and NS2B co-factor domain. Protein Expression and Purification 2016, 119, 124–129. DOI: 10.1016/j.pep.2015.11.022.

(24) Shannon, A. E.; Pedroso, M. M.; Chappell, K. J.; Watterson, D.; Liebscher, S.; Kok, W. M.; Fairlie, D. P.; Schenk, G.; Young, P. R. Product release is rate-limiting for catalytic processing by the Dengue virus protease. Scientific Reports 2016, 6 (1), 37539. DOI: 10.1038/srep37539.

(25) Behnam, M. A. M.; Klein, C. D. Alternate recognition by dengue protease: Proteolytic and binding assays provide functional evidence beyond an induced-fit. bioRxiv 2024, 2024.2004.2015.589505. DOI: 10.1101/2024.04.15.589505.

(26) Behnam, M. A. M.; Basché, T.; Klein, C. D. P. 2,2′5-Bithiophene as sensor tag for ligand–protein binding assays based on Förster resonance energy transfer. Analytical Biochemistry 2023, 682, 115335. DOI: 10.1016/j.ab.2023.115335.

(27) Wu, H.; Bock, S.; Snitko, M.; Berger, T.; Weidner, T.; Holloway, S.; Kanitz, M.; Diederich Wibke, E.; Steuber, H.; Walter, C.;, et al. Novel Dengue Virus NS2B/NS3 Protease Inhibitors. Antimicrobial Agents and Chemotherapy 2015, 59 (2), 1100–1109. DOI: 10.1128/aac.03543-14 (acccessed 2024/03/13).

(28) Müller, B.; Anders, M.; Akiyama, H.; Welsch, S.; Glass, B.; Nikovics, K.; Clavel, F.; Tervo, H.-M.; Keppler, O. T.; Kräusslich, H.-G. HIV-1 Gag Processing Intermediates Trans-dominantly Interfere with HIV-1 Infectivity*. Journal of Biological Chemistry 2009, 284 (43), 29692–29703. DOI: 10.1074/jbc.M109.027144.

(29) Yang, C.-C.; Hsieh, Y.-C.; Lee, S.-J.; Wu, S.-H.; Liao, C.-L.; Tsao, C.-H.; Chao, Y.-S.; Chern, J.-H.; Wu, C.-P.; Yueh, A. Novel Dengue Virus-Specific NS2B/NS3 Protease Inhibitor, BP2109, Discovered by a High-Throughput Screening Assay. Antimicrobial Agents and Chemotherapy 2011, 55 (1), 229–238. DOI: 10.1128/aac.00855-10 (acccessed 2024/03/13).

(30) Steuer, C.; Heinonen, K. H.; Kattner, L.; Klein, C. D. Optimization of Assay Conditions fo r Dengue Virus Protease: Effect of Various Polyols and Nonionic Detergents. Journal of Biomolecular Screening 2009, 14 (9), 1102–1108. DOI: 10.1177/1087057109344115 (acccessed 2024/03/13).

(31) Li, J.; Lim, S. P.; Beer, D.; Patel, V.; Wen, D.; Tumanut, C.; Tully, D. C.; Williams, J. A.; Jiricek, J.; Priestle, J. P.;, et al. Functional Profiling of Recombinant NS3 Proteases from All Four Serotypes of Dengue Virus Using Tetrapeptide and Octapeptide Substrate Libraries *. Journal of Biological Chemistry 2005, 280 (31), 28766–28774. DOI: 10.1074/jbc.M500588200 (acccessed 2024/03/13).

(32) Ezgimen, M. D.; Mueller, N. H.; Teramoto, T.; Padmanabhan, R. Effects of detergents on the West Nile virus protease activity. Bioorganic & Medicinal Chemistry 2009, 17 (9), 3278–3282. DOI: 10.1016/j.bmc.2009.03.050.

(33) Krupa, J. C.; Mort, J. S. Optimization of Detergents for the Assay of Cathepsins B, L, S, and K. Analytical Biochemistry 2000, 283 (1), 99–103. DOI: 10.1006/abio.2000.4621.

(34) Minh, D. D. L.; Chang, C.-e.; Trylska, J.; Tozzini, V.; McCammon, J. A. The Influence of Macromolecular Crowding on HIV-1 Protease Internal Dynamics. Journal of the American Chemical Society 2006, 128 (18), 6006–6007. DOI: 10.1021/ja060483s.

(35) Okamoto, D. N.; Oliveira, L. C. G.; Kondo, M. Y.; Cezari, M. H. S.; Szeltner, Z.; Juhász, T.; Juliano, M. A.; Polgár, L.; Juliano, L.; Gouvea, I. E. Increase of SARS-CoV 3CL peptidase activity due to macromolecular crowding effects in the milieu composition. 2010, 391 (12), 1461–1468. DOI: doi:10.1515/bc.2010.145 (acccessed 2024-04-11).

(36) Behnam, M. A. M.; Nitsche, C.; Vechi, S. M.; Klein, C. D. C-Terminal Residue Optimization and Fragment Merging: Discovery of a Potent Peptide-Hybrid Inhibitor of Dengue Protease. ACS Medicinal Chemistry Letters 2014, 5 (9), 1037–1042. DOI: 10.1021/ml500245v.

(37) Nitsche, C.; Schreier, V. N.; Behnam, M. A. M.; Kumar, A.; Bartenschlager, R.; Klein, C. D. Thiazolidinone–Peptide Hybrids as Dengue Virus Protease Inhibitors with Antiviral Activity in Cell Culture. Journal of Medicinal Chemistry 2013, 56 (21), 8389–8403. DOI: 10.1021/jm400828u.

(38) Steuer, C.; Gege, C.; Fischl, W.; Heinonen, K. H.; Bartenschlager, R.; Klein, C. D. Synthesis and biological evaluation of α-ketoamides as inhibitors of the Dengue virus protease with antiviral activity in cell-culture. Bioorganic & Medicinal Chemistry 2011, 19 (13), 4067–4074. DOI: 10.1016/j.bmc.2011.05.015.

(39) Welsch, S.; Miller, S.; Romero-Brey, I.; Merz, A.; Bleck, C. K. E.; Walther, P.; Fuller, S. D.; Antony, C.; Krijnse-Locker, J.; Bartenschlager, R. Composition and Three-Dimensional Architecture of the Dengue Virus Replication and Assembly Sites. Cell Host & Microbe 2009, 5 (4), 365–375. DOI: 10.1016/j.chom.2009.03.007.

(40) Springsteen, G.; Wang, B. A detailed examination of boronic acid–diol complexation. Tetrahedron 2002, 58 (26), 5291–5300. DOI: 10.1016/S0040-4020(02)00489-1.

(41) Lei, J.; Hansen, G.; Nitsche, C.; Klein, C. D.; Zhang, L.; Hilgenfeld, R. Crystal structure of Zika virus NS2B-NS3 protease in complex with a boronate inhibitor. Science 2016, 353 (6298), 503–505. DOI: 10.1126/science.aag2419 (acccessed 2024/05/05).

(42) Hinkes, S. P. A.; Kämmerer, S.; Klein, C. D. P. Diversity-oriented synthesis of peptide-boronic acids by a versatile building-block approach. Chemical Science 2020, 11 (36), 9898–9903, 10.1039/D0SC03999C. DOI: 10.1039/D0SC03999C.

(43) de la Cruz, L.; Chen, W.-N.; Graham, B.; Otting, G. Binding mode of the activity-modulating C-terminal segment of NS2B to NS3 in the dengue virus NS2B–NS3 protease. 2014, 281 (6), 1517–1533. DOI: 10.1111/febs.12729.

(44) Zhu, L.; Yang, J.; Li, H.; Sun, H.; Liu, J.; Wang, J. Conformational change study of dengue virus NS2B-NS3 protease using 19F NMR spectroscopy. Biochemical and Biophysical Research Communications 2015, 461 (4), 677–680. DOI: 10.1016/j.bbrc.2015.04.090.

