## Supplemental Information for "Antiviral Drug Discovery with an Optimized Biochemical Dengue Protease Assay: Improved Predictive Power for Antiviral Efficacy"

### Substrate Synthesis

DENV protease substrates peptides were synthesised according to standard Fmoc SPPS procedure, as described for the dengue FRET substrate.[1] Purity was determined by RP-HPLC using the same method as for the inhibitors. The respective sequences were for FRET: 2-Abz-Nle-Lys-Arg-Arg-Ser-(3-$NO_2$)-Tyr-$NH_2$, AMC: Bz-Nle-Lys-Arg-Arg-AMC. The *C*-terminal Arg-AMC of the AMC substrate was introduced according to the following procedure:

A round-bottom flask was charged with the according intermediate peptide (1.0 eq.) synthesised on solid support. EDC*HCl (1.1 eq.), *H*-Arg-AMC*2 HCl (1.1 eq.) and HOAt (1.1 eq.) were added and the mixture was dissolved in 5 mL DMF. The resulting solution was cooled to 0 °C and basified with DIPEA to pH 8-9. The resulting pale-yellow solution was stirred for 30 min at 0 °C for 30 min and at room temperature for 3 h. Then, the solution was diluted with ethyl acetate (20 mL) and washed with 0.1 N aq. HCl (3 x 20 mL), sat. aq. NaCl solution (1 x 20 mL), sat. aq. $NaHCO_3$ solution (3 x 20 mL) and sat. aq. NaCl solution (1 x 20 mL) subsequently. The organic phase was dried over anhydrous $MgSO_4$, filtered and dried *in vacuo* to give a pale-yellow crude. This was dissolved in a solution of TFA/TIPS/$H_2O$ (95:2.5:2.5 v/v, 5 mL) and stirred for 3 h at room temperature. The solution was then added dropwise to cold diethyl ether (20 mL) to obtain a colourless precipitate that was washed with additional ether (3 x 20 mL). After concentration *in vacuo*, the precipitate was further purified by RP-HPLC using the same method as for the inhibitors (see General Procedure for Synthesis of Inhibitors and Intermediates on Solid Support).

### DENV Protease Relative Inhibition Assay

The DENV and WNV protease relative inhibition assays were performed as described before.[1, 2] Continuous enzymatic assays were performed in black 96-well V-bottom plates (Greiner Bio-One, Germany) using a BMG Labtech Fluostar OPTIMA Microtiter fluorescence plate reader at an excitation wavelength of 330 nm (FRET substrate) resp. 355 nm (AMC substrate) and an emission wavelength of 430 nm (FRET substrate) resp. 460 nm (AMC substrate). Stock solutions of the inhibitors (10 mM in DMSO) were diluted to a final concentration of 50 μM in triplicates, and preincubated for 15 min with the DENV-2 protease (100 nM) in the assay buffer. The reaction was then initiated by the addition of the respective substrate (final concentration 50 μM) to obtain a final assay volume of 100 μL per well. The enzymatic activity was monitored for 15 min and determined as a slope of relative fluorescence units per second (RFU/s) for each concentration. Compounds **MB-8**[3] and **MB-53**[4] were used as control inhibitors. Percentage inhibition was calculated relative to a positive control (without the inhibitor), as a mean of the triplicates and respective standard deviation. For $IC_{50}$ calculations, data were fitted and calculated with Prism 6.01 (GraphPad Software, Inc.) using a 4-parameter nonlinear dose-response curve.

The compositions of the buffers were the following:

*C7.5-Tris*: 50 mM Tris-HCl, 1 mM CHAPS, 10% ethylene glycol, pH 7.5

*C9.0-Tris*: 50 mM Tris-HCl, 1 mM CHAPS, 10% ethylene glycol, pH 9.0

*B7.5-Tris*: 50 mM Tris-HCl, 0.0016% Brij 58, 10% ethylene glycol, pH 7.5

*B9.0-Tris*: 50 mM Tris-HCl, 0.0016% Brij 58, 10% ethylene glycol, pH 9.0

*C7.5-HEPES*: 25 mM HEPES, 1 mM CHAPS, 10% ethylene glycol, pH 7.5

*C7.5-MOPS*: 25 mM MOPS, 1 mM CHAPS, 10% ethylene glycol, pH 7.5

*C7.5-PBS*: Dulbeccos phosphate-buffered saline, 1 mM CHAPS, 10% ethylene glycol, pH 7.5

*C7.5-PIPES*: 25 mM PIPES, 1 mM CHAPS, 10% ethylene glycol, pH 7.5

**Cell Culture**

If not stated otherwise, HeLa cells were maintained in DMEM supplemented with 100 U/mL penicillin, 100 µg/mL streptomycin, and 10% heat-inactivated FCS. During infection of Huh-7 cells, DMEM was supplemented with 10 mM HEPES.

C6/36 (CLR-1660) mosquito cell line (American Type Culture Collection (ATCC)) were cultured in Eagle's Minimum Essential Medium (EMEM) (Lonza) supplemented with 10% heat-inactivated fetal bovine serum (HIFBS) (Lonza Benelux BV, Breda, The Netherlands), 0.75% sodium bicarbonate (NaHCO3) (Lonza), 10 mM HEPES buffer (Lonza) and 1% penicillin, streptomycin (Pen-Strep) (Lonza)) at 28°C incubator without $CO_2$. Vero cell (ATCC® CCL-81™, Manassas, VA, USA) cultured in complete media (composition: Dulbecco's modified Eagle medium (DMEM) with 10% HI-FBS (Lonza Benelux BV, Breda, The Netherlands), supplemented with 0.75% NaHCO3, 10 mM HEPES buffer (Lonza) and 1% penicillin, streptomycin (Pen-Strep) (Lonza)) at 37°C in a humified incubator with 5% $CO_2$. All three Cell lines were routinely tested negative for mycoplasma using an in-house developed RT-PCR primer and probes.

**DENV2 Reporter Gene Assay (DENV2proHeLa)**

Stable cells were seeded into 96-well plates with a density of 2 × 104 cells per well and treated immediately with inhibitors of the dengue virus protease in a final volume of 100 µL. After incubation for 24 h at 37 °C, the medium was removed, and cells were lysed by adding 25 µL of lysis buffer (Promega) for 15 min at room temperature. Luciferase activity was recorded using a FLUOstar omega plate reader (BMG Labtech) with injections of 100 µL per well of coelenterazine (2.75 µM in PBS). Luminescence was recorded for 5 s. Each concentration was assayed in triplicates. Percent inhibition was calculated in relation to an untreated control. For $EC_{50}$ calculations, data were fitted and calculated with Prism 6.01 (GraphPad Software, Inc.) using a 4-parameter nonlinear dose-response curve with background subtraction (wells without cells).

**Cytotoxicity**

Cell viability in Hela cells in the presence of compound dilutions was determined using CellTiter-Blue (Promega) according to the manufacturer's instructions. Plates were prepared in parallel to the cell-based DENV reporter assay with analogous treatment. Each concentration was assayed in triplicates.

### Procedure Antiviral Testing

#### Virus

Dengue serotypes-2 (VR1584, New Guinea C) was first amplified in C6/36 cells. The resulting supernatant containing the virus was then used to infect Vero cells at a multiplicity of infection (MOI) of 0.01. After 72 hours, the virus was harvested. The harvested virus stocks were clarified by centrifugation and stored at -80°C.

The titers of Dengue serotypes-2 were determined by incubating 10-fold serial dilutions of virus stock on Vero cells for 4 days at 37°C with 5% $CO_2$. To determine the number of virus-infected cells an In-house developed immunostaining method was used. Briefly, the Infected cells were fixed with 2.5% formalin and permeabilized with 0.1% Triton-X-100 in 70% ethanol. The infected cells were stained with Rabbit anti-flavivirus group antigen monoclonal antibody (Absolute antibody) and detected with Goat anti-rabbit IgG (H+L) Highly Cross-Adsorbed Secondary Antibody, Alexa Fluor Plus 488 (2 mg), (Invitrogen). Following this, wells were incubated with 4',6-Diamidino-2-Phenylindole, Dihydrochloride (DAPI) (Thermo Fisher Scientific), to counterstain the nucleus. Plates were scanned using Cytation1V Imaging Reader (BioTek) at a 4× objective and analyzed by the Gen5 software (BioTek). The virus stock titer was calculated using the Karber formula.[5]

#### DENV-2 specific antiviral assay

The antiviral activity of compounds against DENV-2 was determined using in-house developed DENV-2 specific antiviral assay. Briefly, a day before the experiment 96 well substrate plate was prepared using Vero cell ($2 \times 10^4$ cells/well) cultured in complete media as mentioned above. The cells were infected with 0.1moi of DENV-2, except the cell control and incubated at 37°C in 5% $CO_2$ for one hour. During the time of incubation, the compounds were three-fold serially diluted, in four steps, in infection media (composition: DMEM supplemented with 0.75% NaHCO3, 10mM HEPES buffer, 1% Pen-Strep and 1% HI-FBS), starting at 25µM to 0.90µM. After one hour of incubation the monolayer was washed once with PBS followed by the addition of diluted compounds and incubated at 37°C in 5% $CO_2$ for 48 hours. To quantify the percentage of infected cells we used an in-house developed immunofluorescent assay, as described above. The antiviral activity (inhibition percentage) of each compound was determined by comparing the number of infected cells with the virus control. Each compound was tested in duplicate.

### Biological Evaluation Supplementary

**Table S1**. Inhibitory activity of compounds with modified amino groups against the isolated DENV-2 protease, in the DENV2proHeLa as well as cytotoxicity in DENV2proHeLa cells.

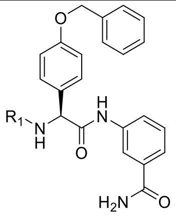

| **No.** | **$R_1$** | **DENV[a]** | | **DENV2proHeLa[b]** | | **$CC_{50}$[c] [µM]** |
|---|---|---|---|---|---|---|
| | | ***C9.0-Tris* FRET** | ***C7.5-HEPES* AMC** | | | |
| | | **% [50 µM]** | | **% [12.5 µM]** | **$EC_{50}$ [µM]** | |
| **1**[6] | 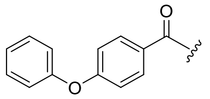 | 19 | 100<br>$IC_{50}$ 13 µM | 83 | 0.7 | > 100 |
| **2**[6] | 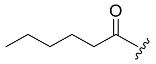 | n.i. | 64 | 76 | 4.6 | > 50 |
| **3** | 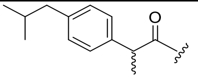 | 9.2 | 100<br>$IC_{50}$ 8.0 µM | 84 | 3.8 | > 25 |
| **4** | 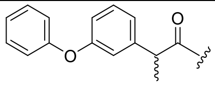 | n.i. | 100<br>$IC_{50}$ 9.7 µM | 86 | 4.0 | > 50 |
| **5** | 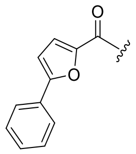 | n.d.[d] | 100<br>$IC_{50}$ 9.3 µM | 85 | 5.9 | > 100 |
| **6** | 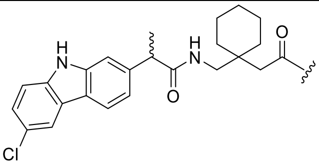 | n.i. | 99<br>$IC_{50}$ 8.0 µM | 60 | 2.9 | > 50 |
| **7** | 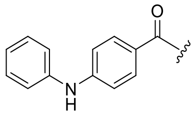 | n.d.[d] | 99<br>$IC_{50}$ 9.5 µM | 93 | 0.7 | ≥ 100 |
| **8** | 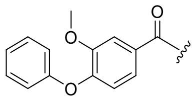 | n.i. | 99<br>$IC_{50}$ 12 µM | 86 | 1.4 | > 100 |
| **9** | 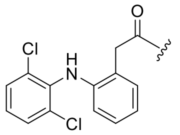 | n.i. | 99<br>$IC_{50}$ 10 µM | 60 | 8.6 | > 50 |
| **10** | 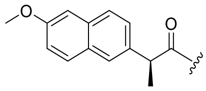 | 20 | 99<br>$IC_{50}$ 12 µM | 71 | 3.7 | > 100 |
| **11** | 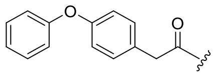 | n.i. | 99<br>$IC_{50}$ 14 µM | 78 | 6.9 | > 50 |

**Table S1**. Inhibitory activity of compounds with modified amino groups against the isolated DENV-2 protease, in the DENV2proHeLa as well as cytotoxicity in DENV2proHeLa cells.

| | | | | | | |
|---|---|---|---|---|---|---|
| **12** | 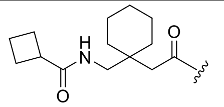 | n.i. | 98<br>IC$_{50}$ 9.2 µM | 81 | 3.8 | > 100 |
| **13** | 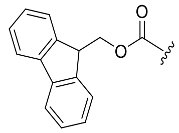 | n.i. | 98<br>IC$_{50}$ 8.7 µM | 80 | 4.8 | > 50 |
| **14** | 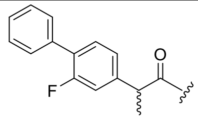 | n.i. | 97<br>IC$_{50}$ 13 µM | 77 | 5.8 | > 50 |
| **15** | 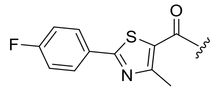 | n.i. | 97<br>IC$_{50}$ 12 µM | 88 | 3.4 | > 100 |
| **16** | 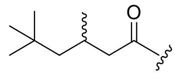 | 22 ± 14 | 96<br>IC$_{50}$ 11 µM | 61 | 11 | ≥ 100 |
| **17** | 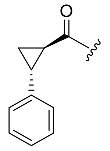 | 12 | 96<br>IC$_{50}$ 15 µM | 72 | 8.2 | > 100 |
| **18** | 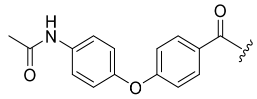 | n.i. | 95<br>IC$_{50}$ 17 µM | 67 | 3.1 | > 50 |
| **19** | 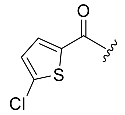 | n.i. | 95<br>IC$_{50}$ 16 µM | 61 | 11 | > 50 |
| **20** | 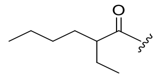 | 19 | 93<br>IC$_{50}$ 22 µM | 63 | 11 | > 50 |
| **21** | 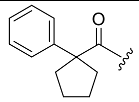 | 14 | 93<br>IC$_{50}$ 6.6 µM | 87 | 6.0 | 36 |
| **22** | 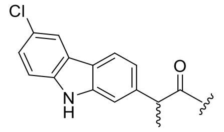 | 12 | 93<br>IC$_{50}$ 13 µM | 75 | 2.7 | 19 |
| **23** | 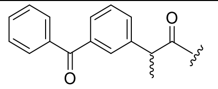 | 12 | 91<br>IC$_{50}$ 16 µM | 78 | 2.4 | > 100 |
| **24** | 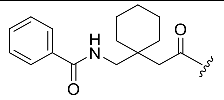 | n.i. | 90<br>IC$_{50}$ 12 µM | 79 | 3.7 | > 50 |
| **25** | 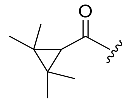 | 14 | 89<br>IC$_{50}$ 14 µM | 66 | 8.8 | > 100 |
| **26** | 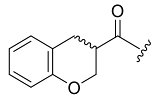 | n.i. | 88<br>IC$_{50}$ 16 µM | 77 | 6.8 | > 50 |
| **27** | 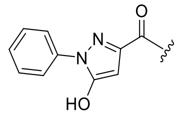 | 25 | 86<br>IC$_{50}$ 30 µM | 24 | 39 | > 50 |

**Table S1**. Inhibitory activity of compounds with modified amino groups against the isolated DENV-2 protease, in the DENV2proHeLa as well as cytotoxicity in DENV2proHeLa cells.

| | | | | | | |
|---|---|---|---|---|---|---|
| **28** | 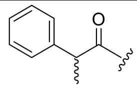 | n.i. | 83<br>$IC_{50}$ 17 µM | 77 | 8.0 | > 50 |
| **29** | 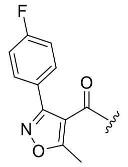 | n.i. | 82<br>$IC_{50}$ 14 µM | 64 | 8.1 | > 50 |
| **30** | 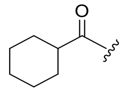 | 12 | 77 | 69 | 19 | > 50 |
| **31** | 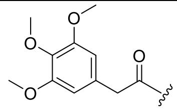 | n.i. | 76 | 75 | 12 | > 50 |
| **32** | 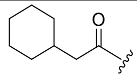 | 20 | 71<br>$IC_{50}$ 34 µM | 74 | 6.4 | > 50 |
| **33** | 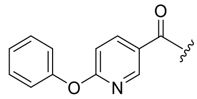 | n.i. | 70<br>$IC_{50}$ 34 µM | 55 | 8.8 | > 100 |
| **34** | 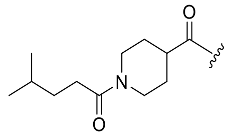 | n.i. | 69 | 53 | 11 | > 100 |
| **35** | 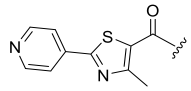 | n.i. | 69 | 78 | 5.1 | > 50 |
| **36** | 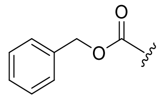 | 16 | 64<br>$IC_{50}$ 28 µM | 60 | 5.0 | ≥ 50 |
| **37** | 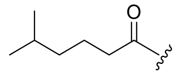 | 12 | 63<br>$IC_{50}$ 36 µM | 67 | 8.2 | > 100 |
| **38** | 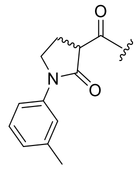 | n.i. | 63 | 31 | 19 | > 50 |
| **39** | 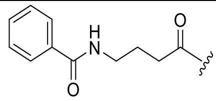 | 13 | 61 | 18 | 23 | > 50 |
| **40** | 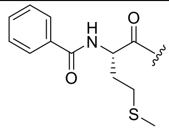 | n.i. | 61<br>$IC_{50}$ 39 µM | 43 | 17 | > 50 |
| **41** | 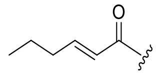 | 15 | 57 | 67 | 9.4 | > 100 |
| **42** | 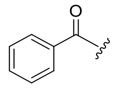 | n.i. | 56 | 80 | 8.5 | > 100 |
| **43** | 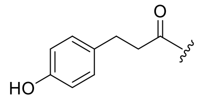 | n.i. | 56 | 65 | 15 | 34 |

**Table S1**. Inhibitory activity of compounds with modified amino groups against the isolated DENV-2 protease, in the DENV2proHeLa as well as cytotoxicity in DENV2proHeLa cells.

| | | | | | | |
|---|---|---|---|---|---|---|
| **44** | 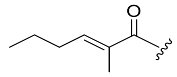 | n.i. | 53 | 69 | 7.8 | > 50 |
| **45** | 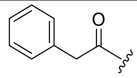 | 15 | 53 | 42 | 14 | > 50 |
| **46** | 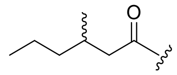 | 13 | 52 | 69 | 7.3 | > 25 |
| **47** | 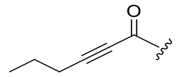 | n.i. | 52<br>$IC_{50}$ 45 µM | 62 | 7.1 | > 25 |
| **48** | 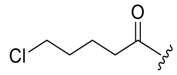 | n.i. | 48 | 66 | 6.9 | > 25 |
| **49** | 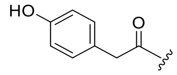 | n.i. | 44 | 36 | 15 | > 50 |
| **50** | 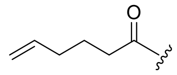 | 12 ± 17 | 39 | 52 | 12 | > 50 |
| **51** | 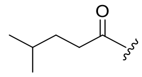 | 20 ± 14 | 37 | 48 | 8.8 | > 50 |
| **52** | 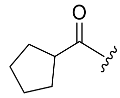 | 14 ± 18 | 34 | 25 | 23 | > 100 |
| **53** | 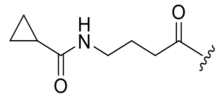 | 13 | 33 | n.i. | > 25 | > 50 |
| **54** | 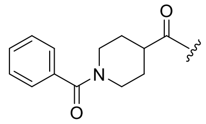 | n.i. | 31 | 37 | 15 | > 100 |
| **55** | 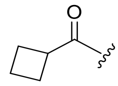 | 19 | 30 | 10 | 25 | > 50 |
| **56** | 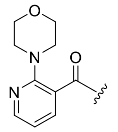 | n.i. | 30 | 54 | 11 | > 50 |
| **57** | 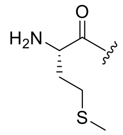 | n.i. | 29 | 20 | 36 | > 50 |
| **58** | 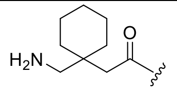 | n.i. | 26 | 23 | 21 | > 50 |
| **59** | 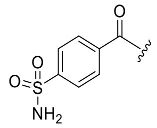 | n.i. | 26 | 15 | 35 | > 50 |
| **60** | 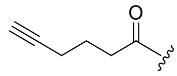 | n.i. | 16 | 54 | 9.2 | > 50 |
| **61** | 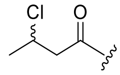 | 17 | 16 | 43 | 13 | > 50 |
| **MB-53**[7] | | 99 | 87 | 37 (41) | n.d. | n.d. |

**Table S1**. Inhibitory activity of compounds with modified amino groups against the isolated DENV-2 protease, in the DENV2proHeLa as well as cytotoxicity in DENV2proHeLa cells.

| | | | | | |
|---|---|---|---|---|---|
| **NK-189**[7] | 95 | 97 | 73 (80) | n.d. | n.d. |

[a]Inhibition against DENV NS2B-NS3 serotype 2 at 50 µM substrate concentration. [b]Inhibition values against DENV serotype 2 protease in reporter gene assay in HeLa cells. [c]Measured in DENV2proHeLa cells. If inhibition ≤ 10% = no inhibition (n.i.). SD ≤ 10% unless otherwise stated. [d]not determined due to autofluorescence at the used assay conditions.

**Table S2**. Inhibitory activity of compounds with modified amino acid core against the isolated DENV-2 protease, in the DENV2proHeLa as well as cytotoxicity in DENV2proHeLa cells.

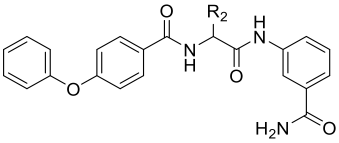

| No. | $R_2$ | DENV[a] | | DENV2proHeLa[b] | | $CC_{50}$[c] [µM] |
|---|---|---|---|---|---|---|
| | | *C9.0-Tris* FRET | *C7.5-HEPES* AMC | | | |
| | | % [50 µM] | | % [12.5 µM] | $EC_{50}$ [µM] | |
| **1**[6] | 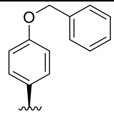 | 19 | 100<br>$IC_{50}$ 13 µM | 83 | 0.7 | > 100 |
| **62** | 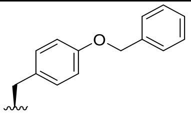 | n.i. | 100<br>$IC_{50}$ 6.2 µM | 74 | 4.5 | > 50 |
| **63** | 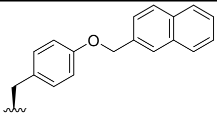 | n.i. | 100<br>$IC_{50}$ 4.8 µM | 82 | 5.1 | 16 |
| **64** | 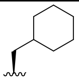 | 13 | 99<br>$IC_{50}$ 11 µM | 82 | 7.3 | 40 |
| **65** | 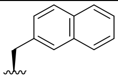 | 13 | 99<br>$IC_{50}$ 5.9 µM | 55 | 7.4 | > 50 |
| **66** | 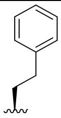 | 14 | 94<br>$IC_{50}$ 11 µM | 67 | 8.6 | > 50 |
| **67** | 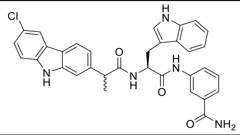 | n.i. | 91<br>$IC_{50}$ 7.5 µM | 91 | 2.6 | 16 |
| **68** | 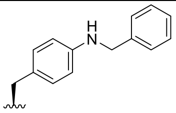 | 13 | 80<br>$IC_{50}$ 15 µM | n.i. | 25 | > 50 |
| **69** | 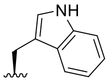 | n.i. | 80<br>$IC_{50}$ 21 µM | 65 | 8.5 | > 50 |
| **70** | 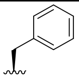 | 13 | 78<br>$IC_{50}$ 21 µM | 55 | 12 | > 50 |
| **71** | 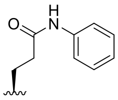 | 17 | 72<br>$IC_{50}$ 23 µM | 45 | 20 | > 50 |
| **72** | 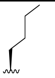 | n.i. | 67 | 57 | 9.5 | ≥ 50 |
| **73** | 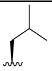 | 17 | 64 | 36 | 16 | ≥ 50 |

**Table S2**. Inhibitory activity of compounds with modified amino acid core against the isolated DENV-2 protease, in the DENV2proHeLa as well as cytotoxicity in DENV2proHeLa cells.

| | | | | | | |
|---|---|---|---|---|---|---|
| **74** | 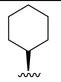 | n.i. | 59 | 73 | 4.6 | > 25 |
| **75** | 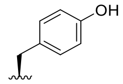 | n.i. | 53 | 45 | 12 | > 50 |
| **76** | 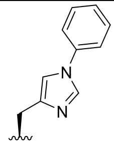 | n.i. | 52 | 51 | 11 | > 50 |
| **77** | 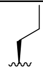 | n.i. | 51 | 42 | 15 | > 50 |
| **78** | 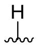 | n.i. | 23 | 37 | > 25 | > 50 |
| **79** | 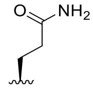 | n.i | 21 | 19 | > 50 | > 50 |

[a]Inhibition against DENV NS2B-NS3 serotype 2 at 50 μM substrate concentration. [b]Inhibition values against DENV serotype 2 protease in reporter gene assay in HeLa cells. [c]Measured in DENV2proHeLa cells. If inhibition ≤ 10% = no inhibition (n.i.). SD ≤ 10% unless otherwise stated.

**Table S3**. Inhibitory activity of selected compounds against the isolated DENV-2 protease, in the DENV2proHeLa as well as reduction of viral load in the antiviral immunofluorescence assay.

| No. | DENV[a] | | DENV2proHeLa[b] | Antiviral Testing[c] |
|---|---|---|---|---|
| | *C9.0-Tris* FRET | *C7.5-HEPES* AMC | | |
| | % [50 µM] | | $EC_{50}$ [µM] | Reduction at 25 µM (%) |
| **NK-189** | 95 | 97 | 0.5 | 21 |
| **7** | < 10 | 99 | 0.7 | < 10 |
| **24** | < 10 | 90 | 3.7 | < 10 |
| **12** | < 10 | 98 | 3.8 | / |
| **4** | < 10 | 100 | 4 | 85 |
| **62** | < 10 | 100 | 4.5 | 19 |
| **5** | n.d.[d] | 99 | 5.9 | 96 |
| **32** | 19 | 71 | 6.4 | 93 |
| **37** | 12 | 97 | 8.2 | 94 |
| **17** | 12 | 96 | 8.2 | 55 |
| **25** | < 10 | 89 | 8.8 | 86 |
| **51** | 20 | 37 | 8.8 | 84 |
| **72** | 17 | 67 | 9.5 | 79 |
| **20** | 19 | 97 | 11 | 90 |
| **16** | < 10 | 96 | 11 | 98 |
| **76** | < 10 | 52 | 11 | 57 |
| **31** | < 10 | 76 | 12 | 11 |
| **MB-53** | 99 | 87 | 16 | / |
| **40** | < 10 | 79 | 17 | / |
| **30** | 12 | 77 | 19 | 28 |
| **GG-92** | 31 | 53 | 20 | 40 |
| **58** | < 10 | 32 | 21 | / |
| **NK-71** | 81 | 45 | 23 | < 10 |
| **GG-80-1** | 36 | 27 | > 25 | 97 |

**Table S3**. Inhibitory activity of selected compounds against the isolated DENV-2 protease, in the DENV2proHeLa as well as reduction of viral load in the antiviral immunofluorescence assay.

| | | | | |
|---|---|---|---|---|
| **GG-69-2** | 36 | 25 | > 25 | 35 |
| **MJ-04-32** | 100 | 101 | > 50 | / |
| **CN-488** | 99 | 82 | > 50 | < 10 |
| **CN-519** | 83 | 52 | > 50 | < 10 |
| **CN-229** | 89 | 43 | > 50 | < 10 |
| **GG-58** | 41 | 42 | > 50 | 32 |
| **CN-230** | 69 | 42 | > 50 | < 10 |
| **CN-279** | 85 | 35 | > 50 | 21 |
| **CN-269** | 83 | 28 | > 50 | 28 |
| **MB-8** | 69 | < 10 | > 50 | / |
| **53** | 13 | < 10 | > 50 | < 10 |

[a]Inhibition against DENV NS2B-NS3 serotype 2 at 50 µM substrate concentration. [b]Inhibition values against DENV serotype 2 protease in reporter gene assay in HeLa cells. [c]Reduction of viral load in anti-DENV immunofluorescence assay. [d]not determined due to autofluorescence at the used assay conditions. / = increased viral load. If inhibition ≤ 10% = no inhibition (n.i.). SD ≤ 10% unless otherwise stated.

### Structural Formulae of Previously Published DENV Protease Inhibitors

**NK-189**[7]

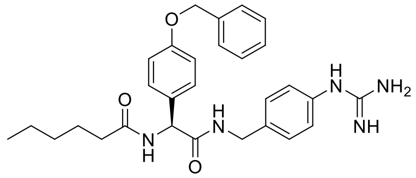

**MB-8**[3]

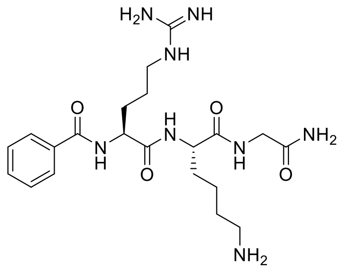

**MB-53**[4]

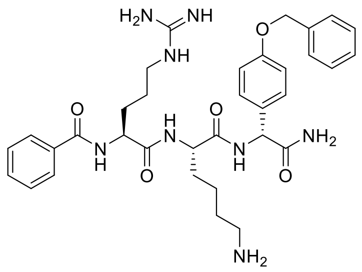

**MJ-04-32**[8]

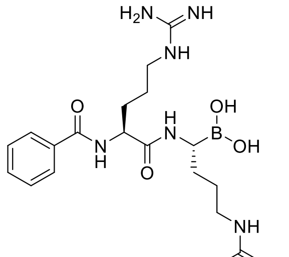

**GG-58**[9]

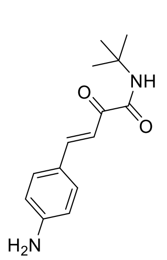

**GG-69-2**[9]

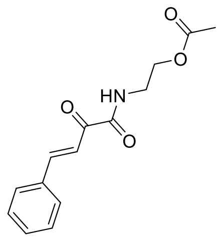

**GG-80-1**[9]

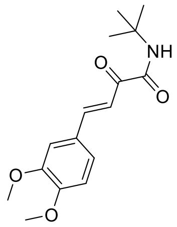

**GG-92**[9]

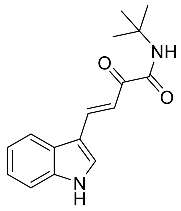

**CN-229**[10]

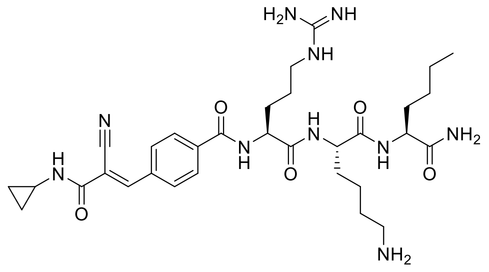

**CN-230**[10]

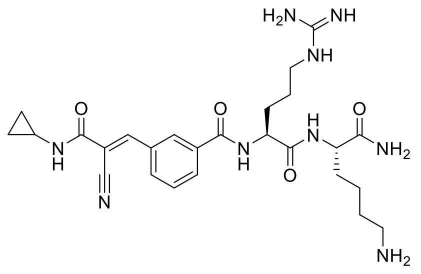

**CN-269**[10]

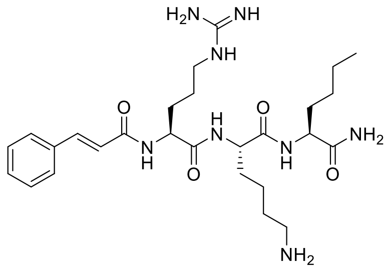

**CN-279**[10]

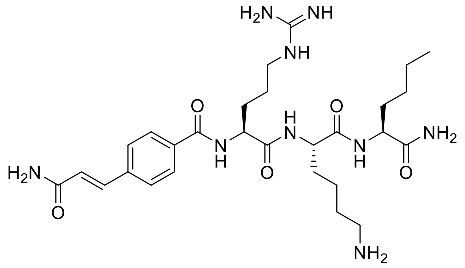

**CN-488**[2]

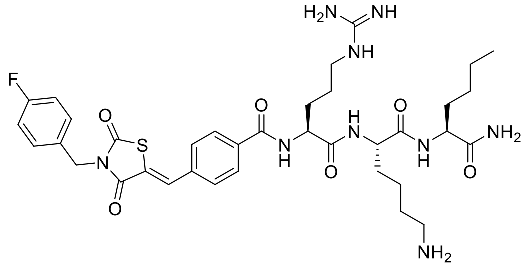

**CN-519**[2]

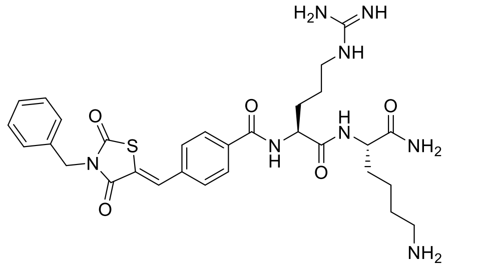

**Chemical and Analytical Supplementary**

All chemicals for the synthesis of precursors were obtained from Sigma-Aldrich (Germany), Alfa Aesar (Germany), Thermo Fisher Scientific (Germany/United States), Acros Organics (Belgium), TCI Europe (Belgium), and Carbolution Chemicals (Germany) and were of analytical grade. The amino acids were purchased from Carbolution Chemicals (Germany), Alfa Aesar (Germany), and TCI Europe (Belgium). HATU and Fmoc-Rink amide resin (75−150 mesh; loading capacity: 0.71 mmol/g resin) were purchased from Iris Biotech (Germany). Solvents were used as obtained from the commercial suppliers.

Mass spectra (HR-ESI, see Table S5) of all compounds were measured on a Bruker micrOTOF-Q II instrument.

**General Procedure for the Synthesis of Inhibitors and Intermediates on a Solid Support**. All sequences and intermediates were assembled by stepwise solid-phase synthesis on Rink amid resin using the standard Fmoc strategy, as previously described.[1, 2, 11] Solid-phase synthesis was done manually in plastic syringes equipped with a frit; all steps were performed at room temperature under continuous shaking. The Rink amide resin was preswollen in DCM for at least 20 min and then washed 3× with DMF. For Fmoc deprotection, a piperidine solution (10% in DMF) was added 2× for 10 and 5 min. Following each deprotection or coupling step, the resin was washed 3× with DMF, 3× with DCM, and again 3× with DMF. COMU and TMP were used for coupling steps. In detail, the coupling solution contained the $N_\alpha$-Fmoc protected amino acid or the respective building block (2.0−3.0 equiv), COMU (2.0−3.0 equiv), and TMP (2.3−3.9 equiv) in DMF (1.0 mL per 100 mg of resin). The solution was added to the resin, and the mixture was shaken for 60−120 min. Afterward, the resin was washed as described before. Fmoc deprotection and coupling steps were iteratively repeated until the desired sequence was obtained. The resin loaded with the finished peptide was washed 5× with diethyl ether and dried under reduced pressure. The final product was cleaved 2× off the resin with TFA/TIPS/$H_2O$ solution (95:1:4, 1−2 mL per 100 mg of resin), and the mixture was shaken for 2 h. The cleavage solution was dispensed into cold diethyl ether (35 mL per 100 mg of resin), and the resulting precipitate was centrifuged (4000g, 10 min), washed with diethyl ether/hexane, and dried under reduced pressure. If the peptides were soluble in the organic solvent, then the organic phase was washed 2 times with a small amount of water, and all solvents were removed *in vacuo*. The resulting residue was co-evaporated with toluene several times. All final compounds were purified by preparative RP-HPLC on an ÄKTA Purifier, GE Healthcare (Germany), with an RP-18 pre and main column (Rephospher, Dr. Maisch GmbH, Germany; C18-DE, 5 µm, 30 mm × 16 mm, and 120 mm × 16 mm). The following conditions were used: eluent A, water (0.1% TFA); eluent B, methanol (0.1% TFA) or eluent A, water (0.1% TFA); eluent B, acetonitrile (0.1% TFA); flow rate, 8 mL/min; and gradient, 10% B (2.5 min), 100% B (23.5 min), 100% B (26 min), 10% B (26.1 min), and 10% B (30 min). Detection was performed at 214, 254, and 280 nm. After purification, the organic solvent was evaporated, and the compounds and intermediates were freeze-dried in $H_2O$/ACN and stored at −20 °C.

**3 (mixture of diastereomers)**
$^1$H NMR (300 MHz, DMSO-$d_6$) δ 10.36 (d, $J$ = 16.3 Hz, 1H), 8.63 (dd, $J$ = 20.2, 7.6 Hz, 1H), 8.04 – 7.96 (m, 1H), 7.92 (d, $J$ = 7.4 Hz, 1H), 7.71 (dd, $J$ = 21.7, 8.1 Hz, 1H), 7.52 (t, $J$ = 7.6 Hz, 1H), 7.46 – 7.18 (m, 10H), 7.04 (q, $J$ = 8.6, 8.2 Hz, 3H), 6.93 (d, $J$ = 8.7 Hz, 1H), 5.63 – 5.41 (m, 1H), 5.07 (d, $J$ = 11.1 Hz, 2H), 3.87 (q, $J$ = 6.9 Hz, 1H), 2.39 (d, $J$ = 7.1 Hz, 2H), 1.80 (dq, $J$ = 13.4, 6.6 Hz, 1H), 1.30 (dd, $J$ = 9.8, 7.0 Hz, 3H), 0.84 (d, $J$ = 6.6 Hz, 7H).

**4 (mixture of diastereomers)**
$^1$H NMR (300 MHz, DMSO-$d_6$) δ 10.36 (d, $J$ = 13.4 Hz, 1H), 8.68 (dd, $J$ = 8.3 Hz, 1H), 7.98 (dt, $J$ = 14.6, 1.9 Hz, 1H), 7.94 – 7.87 (m, 1H), 7.75 – 7.64 (m, 1H), 7.55 – 7.47 (m, 1H), 7.46 – 7.23 (m, 12H), 7.15 – 6.91 (m, 7H), 6.82 (dd, $J$ = 8.2, 2.5 Hz, 1H), 5.50 (dd, $J$ = 24.8, 7.5 Hz, 1H), 5.06 (d, $J$ = 8.7 Hz, 2H), 3.94 – 3.84 (m, 1H), 1.28 (dd, $J$ = 10.7, 7.0 Hz, 3H). $^{13}$C NMR (75 MHz, DMSO-$d_6$) δ 173.37, 169.59, 168.20, 158.42, 157.04, 156.78, 144.75, 139.24, 137.43, 135.59, 130.77, 130.43, 130.15, 128.96, 128.87, 128.25, 128.04, 128.01, 123.74, 122.91, 122.61, 122.18, 118.95, 118.89, 118.16, 115.23, 115.15, 69.63, 56.74, 44.36, 19.02.

**5**
$^1$H NMR (300 MHz, DMSO-$d_6$) δ 10.46 (s, 1H), 8.88 (d, $J$ = 7.5 Hz, 1H), 8.05 (d, $J$ = 2.1 Hz, 1H), 7.94 – 7.92 (m, 2H), 7.90 (s, 1H), 7.78 (d, $J$ = 8.2 Hz, 1H), 7.58 – 7.27 (m, 16H), 7.12 (d, $J$ = 3.6 Hz, 1H), 7.04 (d, $J$ = 8.7 Hz, 2H), 5.76 (d, $J$ = 7.2 Hz, 1H), 5.11 (s, 2H). $^{13}$C NMR (75 MHz, DMSO-$d_6$) δ 168.99, 167.77, 158.13, 157.43, 154.91, 146.44, 138.80, 138.52, 137.02, 135.14, 129.54, 129.33, 129.17, 128.89, 128.61, 128.43, 127.80, 127.59, 124.43, 118.81, 116.60, 114.80, 107.63, 69.18.

**6 (mixture of diastereomers)**
$^1$H NMR (500 MHz, DMSO-$d_6$) δ 11.31 (d, $J$ = 4.6 Hz, 1H), 10.34 (d, $J$ = 6.8 Hz, 1H), 8.73 (d, $J$ = 7.1 Hz, 1H), 8.14 – 8.06 (m, 1H), 8.05 – 7.98 (m, 2H), 7.91 (s, 1H), 7.82 (s, 1H), 7.71 (d, $J$ = 8.3 Hz, 1H), 7.50 (d, $J$ = 7.8 Hz, 1H), 7.47 – 7.26 (m, 13H), 7.16 – 7.09 (m, 1H), 7.01 – 6.92 (m, 2H), 5.49 (t, $J$ = 8.1 Hz, 1H), 5.04 (d, $J$ = 19.3 Hz, 2H), 3.87 – 3.78 (m, 1H), 3.30 – 3.19 (m, 1H), 3.10 – 3.01 (m, 1H), 2.16 – 2.04 (m, 2H), 1.45 – 1.03 (m, 17H). $^{13}$C NMR (126 MHz, DMSO-$d_6$) δ 174.32, 171.08, 169.79, 168.23, 158.44, 141.54, 140.95, 139.34, 138.71, 137.43, 135.56, 130.43, 130.30, 129.20, 129.05, 128.86, 128.84, 128.25, 128.04, 125.35, 124.11, 123.19, 122.50, 122.17, 120.72, 120.56, 120.00, 119.30, 119.10, 115.18, 112.71, 109.93, 69.61, 57.15, 46.04, 37.81, 21.51, 19.45.

**7**
$^1$H NMR (300 MHz, DMSO-$d_6$) δ 10.38 (s, 1H), 8.64 – 8.48 (m, 2H), 8.04 (s, 1H), 7.92 (s, 1H), 7.83 (d, J = 8.4 Hz, 2H), 7.77 (d, J = 8.7 Hz, 1H), 7.52 – 7.27 (m, 12H), 7.15 (d, J = 7.6 Hz, 2H), 7.07 – 6.99 (m, 4H), 6.93 (t, J = 7.3 Hz, 1H), 5.73 (d, J = 7.4 Hz, 1H), 5.11 (s, 2H). $^{13}$C NMR (APT, 75 MHz, DMSO-$d_6$) δ 169.91, 168.26, 166.33, 158.24, 147.24, 142.40, 139.37, 135.56, 130.45, 129.83, 129.73, 129.55, 128.89, 128.26, 128.04, 124.16, 122.50, 122.19, 121.59, 119.22, 118.92, 115.15, 114.62, 69.63, 57.70.

**8**
$^1$H NMR (300 MHz, DMSO-$d_6$) δ 10.43 (s, 1H), 8.94 (d, J = 7.3 Hz, 1H), 8.05 (s, 2H), 7.93 (s, 1H), 7.78 (d, J = 8.3 Hz, 1H), 7.70 (s, 1H), 7.58 (d, J = 8.6 Hz, 1H), 7.50 (d, J = 8.8 Hz, 2H), 7.47 – 7.29 (m, 9H), 7.10 (d, J = 7.4 Hz, 1H), 7.07 – 7.00 (m, 3H) 6.90 (d, J = 8.0 Hz, 2H), 5.76 (d, J = 7.2 Hz, 1H), 5.11 (s, 2H), 3.81 (s, 3H). $^{13}$C NMR (APT, 75 MHz, DMSO-$d_6$) δ 169.71, 168.24, 166.13, 158.54, 157.49, 150.98, 147.12, 139.36, 137.49, 135.58, 130.93, 130.30,

130.04, 129.89, 129.69, 129.06, 128.89, 128.26, 128.04, 123.26, 122.57, 122.23, 121.65, 120.55, 119.22, 117.28, 115.20, 113.16, 69.62, 58.02, 56.33.

**9**

$^{1}$H NMR (500 MHz, DMSO-$d_6$) δ 10.46 (s, 1H), 9.14 (d, $J$ = 7.9 Hz, 1H), 8.15 (s, 1H), 8.00 (s, 2H), 7.91 (s, 0H), 7.73 (d, $J$ = 8.2 Hz, 1H), 7.55 – 7.47 (m, 4H), 7.44 – 7.23 (m, 13H), 7.21 – 7.12 (m, 2H), 7.05 – 6.97 (m, 3H), 6.84 (t, $J$ = 7.4 Hz, 2H), 5.63 (d, $J$ = 7.9 Hz, 1H), 5.08 (s, 2H), 3.75 (dd, $J$ = 13.7 Hz, 4H). $^{13}$C NMR (126 MHz, DMSO-$d_6$) δ 171.27, 167.73, 158.03, 142.99, 138.66, 136.97, 135.14, 130.58, 129.92, 129.55, 129.14, 128.50, 128.40, 127.78, 125.13, 120.61, 118.72, 115.83, 114.80, 69.18.

**10**

$^{1}$H NMR (300 MHz, DMSO-$d_6$) δ 10.34 (s, 1H), 8.75 (d, $J$ = 7.7 Hz, 1H), 7.95 (t, $J$ = 1.9 Hz, 1H), 7.89 (s, 1H), 7.80 – 7.72 (m, 3H), 7.66 (dd, $J$ = 7.7, 2.2 Hz, 1H), 7.54 – 7.24 (m, 12H), 7.13 (dd, $J$ = 8.9, 2.6 Hz, 1H), 7.03 (d, $J$ = 8.7 Hz, 2H), 5.58 (d, $J$ = 7.6 Hz, 1H), 5.10 (s, 2H), 4.04 (q, $J$ = 7.0 Hz, 1H), 3.85 (s, 3H), 1.39 (d, $J$ = 7.0 Hz, 3H). $^{13}$C NMR (75 MHz, DMSO-$d_6$) δ 173.27, 169.00, 167.75, 157.99, 156.95, 138.67, 137.18, 137.02, 135.11, 133.10, 130.39, 129.09, 128.54, 128.43, 127.81, 127.60, 126.62, 126.53, 125.33, 122.17, 118.68, 118.51, 114.78, 105.68, 69.18, 66.38, 65.91, 55.11, 44.00, 18.54.

**11**

$^{1}$H NMR (300 MHz, DMSO-$d_6$) δ 10.39 (s, 1H), 8.80 (d, $J$ = 7.5 Hz, 1H), 8.00 (s, 1H), 7.91 (s, 1H), 7.72 (dd, $J$ = 8.1, 2.1 Hz, 1H), 7.51 (d, $J$ = 7.7 Hz, 1H), 7.45 – 7.23 (m, 14H), 7.09 (t, $J$ = 7.3 Hz, 1H), 7.04 – 6.87 (m, 7H), 5.53 (d, $J$ = 7.5 Hz, 1H), 5.07 (s, 2H), 3.55 (s, 2H). $^{13}$C NMR (75 MHz, DMSO-$d_6$) δ 170.48, 169.64, 168.23, 158.48, 157.25, 155.52, 139.24, 137.46, 135.59, 131.99, 131.08, 130.55, 130.43, 129.07, 128.88, 128.26, 128.05, 123.67, 122.64, 122.23, 119.21, 118.99, 118.81, 115.25, 69.64, 57.01, 41.28.

**12**

$^{1}$H NMR (300 MHz, DMSO-$d_6$) δ 10.36 (s, 1H), 8.81 (d, $J$ = 7.1 Hz, 1H), 8.02 (s, 1H), 7.92 (s, 1H), 7.73 (d, $J$ = 7.9 Hz, 1H), 7.52 (d, $J$ = 6.8 Hz, 2H), 7.52 – 7.23 (m, 11H), 7.01 (d, $J$ = 8.7 Hz, 2H), 5.52 (d, $J$ = 7.1 Hz, 1H), 5.09 (s, 2H), 3.31 – 3.19 (m, 1H), 3.13 – 2.96 (m, 2H), 2.17 (s, 2H), 2.16 – 1.94 (m, 3H), 1.90 – 1.67 (m, 1H), 1.56 – 1.12 (m, 1H). $^{13}$C NMR (75 MHz, DMSO-$d_6$) δ 174.34, 170.71, 169.37, 167.79, 158.01, 138.91, 137.00, 135.12, 129.87, 128.77, 128.41, 127.80, 127.60, 122.04, 121.69, 118.66, 114.75, 69.17, 56.75, 38.78, 37.16, 24.81, 24.74, 21.08, 17.81.

**13**

$^{1}$H NMR (300 MHz, DMSO-$d_6$) δ 10.34 (s, 1H), 8.15 (d, $J$ = 8.0 Hz, 1H), 8.01 (s, 1H), 7.94 – 7.83 (m, 3H), 7.80 – 7.71 (m, 3H), 7.52 (d, $J$ = 7.7 Hz, 1H), 7.47 – 7.24 (m, 12H), 7.00 (d, $J$ = 8.5 Hz, 2H), 5.33 (d, $J$ = 7.9 Hz, 1H), 5.09 (s, 2H), 4.31 – 4.15 (m, 3H). $^{13}$C NMR (75 MHz, DMSO-$d_6$) δ 169.77, 168.21, 159.24, 158.51, 156.28, 144.16, 141.11, 139.25, 137.47, 135.57, 129.28, 128.88, 128.25, 128.05, 127.49, 125.87, 122.62, 122.21, 120.51, 119.18, 115.16, 69.62, 54.52, 47.05.

**14 (mixture of diastereomers)**

$^{1}$H NMR (300 MHz, DMSO-$d_6$) δ 10.38 (d, $J$ = 12.3 Hz, 1H), 8.80 (dd, $J$ = 10.1, 7.5 Hz, 1H), 7.99 (dt, $J$ = 18.6, 1.9 Hz, 1H), 7.90 (d, $J$ = 8.7 Hz, 1H), 7.77 – 7.65 (m, 1H), 7.55 – 7.19 (m, 19H), 7.05 – 6.92 (m, 2H), 5.52 (dd, $J$ = 18.9, 7.4 Hz, 1H), 5.07 (d, $J$ = 13.0 Hz, 2H), 4.03 –

3.92 (m, 1H), 1.35 (dd, *J* = 10.8, 7.0 Hz, 3H). $^{13}$C NMR (75 MHz, DMSO-$d_6$) δ 169.60, 169.60, 169.41, 168.18, 158.46, 157.60, 139.15, 137.46, 135.60, 130.63, 129.13, 129.01, 128.88, 128.85, 128.26, 128.05, 128.02, 124.36, 122.22, 121.68, 119.16, 115.27, 115.19, 69.65, 61.99, 40.49, 40.21, 39.92, 39.63, 39.37, 21.71.

**15**
$^{1}$H NMR (300 MHz, DMSO-$d_6$) δ 10.45 (s, 1H), 8.82 (d, *J* = 7.0 Hz, 1H), 8.09 – 7.91 (m, 4H), 7.81 – 7.72 (m, 1H), 7.61 – 7.47 (m, 3H), 7.46 – 7.28 (m, 8H), 7.04 (d, *J* = 8.7 Hz, 1H), 5.68 (d, *J* = 7.0 Hz, 1H), 5.11 (s, 2H), 2.63 (s, 3H). $^{13}$C NMR (75 MHz, DMSO-$d_6$) δ 168.80, 167.76, 165.05, 160.97, 158.17, 155.15, 138.79, 137.00, 135.15, 129.35, 129.08, 128.71, 128.60, 128.43, 127.82, 127.60, 121.78, 118.75, 116.56, 116.26, 114.80, 69.18, 57.49, 17.14.

**16**
$^{1}$H NMR (300 MHz, DMSO-$d_6$) δ 10.36 (d, *J* = 1.7 Hz, 1H), 8.55 – 8.46 (m, 1H), 8.01 (q, *J* = 2.2 Hz, 1H), 7.92 (s, 1H), 7.80 – 7.68 (m, 1H), 7.52 (dd, *J* = 7.9, 1.5 Hz, 1H), 7.47 – 7.25 (m, 10H), 7.00 (d, *J* = 8.7 Hz, 2H), 5.55 (dd, *J* = 7.6, 5.6 Hz, 1H), 5.09 (s, 2H), 2.22 – 2.00 (m, 2H), 2.01 – 1.85 (m, 1H), 1.25 (m, 2H), 1.01 (dd, *J* = 13.9, 6.6 Hz, 1H), 0.94 – 0.86 (m, 2H), 0.84 (d, *J* = 3.4 Hz, 11H). $^{13}$C NMR (75 MHz, DMSO-$d_6$) δ 171.49, 169.35, 167.78, 157.93, 138.86, 137.01, 135.12, 130.18, 128.64, 128.40, 127.78, 127.58, 127.56, 121.70, 118.70, 114.69, 69.15, 56.46, 50.09, 44.59, 30.73, 29.83, 26.96, 22.53.

**17**
$^{1}$H NMR (300 MHz, DMSO-$d_6$) δ 10.42 (d, *J* = 6.0 Hz, 1H), 8.89 (t, *J* = 8.4 Hz, 1H), 8.02 (s, 1H), 7.93 (s, 1H), 7.78 – 7.69 (m, 1H), 7.53 (d, *J* = 7.9 Hz, 1H), 7.45 – 7.21 (m, 9H), 7.23 – 7.08 (m, 2H), 7.06 – 6.96 (m, 2H), 5.60 (t, *J* = 7.0 Hz, 1H), 5.09 (d, *J* = 4.8 Hz, 2H), 3.09 (p, *J* = 7.0 Hz, 2H), 2.31 – 2.19 (m, 3H), 1.17 (t, *J* = 7.3 Hz, 4H). $^{13}$C NMR (75 MHz, DMSO-$d_6$) δ 170.89, 169.30, 167.78, 158.01, 141.03, 138.80, 136.99, 135.13, 130.15, 128.72, 128.62, 128.41, 128.31, 128.27, 127.79, 127.58, 125.85, 125.79, 121.74, 118.71, 114.79, 69.17, 45.72, 24.90, 24.16, 8.60.

**18**
$^{1}$H NMR (300 MHz, DMSO-$d_6$) δ 10.39 (s, 1H), 9.99 (s, 1H), 8.81 (d, J = 7.8 Hz, 1H), 8.03 (s, 1H), 7.95 (d, J = 8.8 Hz, 4H), 7.77 (d, J = 8.0 Hz, 1H), 7.62 (d, J = 8.8 Hz, 2H), 7.50 – 7.33 (m, 10H), 7.06 – 7.01 (m, 3H), 6.97 (d, J = 8.7 Hz, 2H), 5.73 (d, J = 7.2 Hz, 1H), 5.10 (s, 2H), 2.04 (s, 3H).

**19**
$^{1}$H NMR (300 MHz, DMSO-$d_6$) δ 10.41 (s, 1H), 9.08 (d, *J* = 7.2 Hz, 1H), 8.02 – 7.98 (m, 1H), 7.95 – 7.88 (m, 2H), 7.73 (dd, *J* = 8.1, 2.1 Hz, 1H), 7.51 (d, *J* = 7.7 Hz, 1H), 7.46 – 7.25 (m, 9H), 7.16 (d, *J* = 4.1 Hz, 1H), 7.01 (d, *J* = 8.5 Hz, 2H), 5.66 (d, *J* = 7.1 Hz, 1H), 5.08 (s, 2H). $^{13}$C NMR (75 MHz, DMSO-$d_6$) δ 169.32, 168.25, 160.58, 158.63, 139.25, 138.75, 137.45, 135.58, 133.80, 129.78, 129.72, 129.60, 129.08, 128.89, 128.61, 128.27, 128.04, 122.66, 122.26, 119.22, 115.25, 69.62, 57.71.

**20**
$^{1}$H NMR (500 MHz, Acetone-$d_6$) δ 9.74 (d, *J* = 3.6 Hz, 1H), 8.12 – 8.07 (m, 1H), 7.87 (t, *J* = 7.7 Hz, 1H), 7.77 (d, *J* = 7.4 Hz, 1H), 7.57 (dd, *J* = 7.5, 2.1 Hz, 1H), 7.46 (d, *J* = 8.1 Hz, 4H), 7.40 – 7.29 (m, 4H), 7.00 (d, *J* = 8.4 Hz, 2H), 6.67 (s, 1H), 5.77 – 5.72 (m, 1H), 5.11 (s, 2H), 2.39 – 2.31 (m, 1H), 1.67 – 1.51 (m, 1H), 1.49 – 1.14 (m, 4H), 0.93 (t, *J* = 7.4 Hz, 1H), 0.88 – 0.78 (m, 4H). $^{13}$C NMR (126 MHz, Acetone-$d_6$) δ 175.00, 169.28, 158.69, 158.67, 139.05,

137.40, 130.45, 128.88, 128.86, 128.60, 128.37, 127.71, 127.49, 127.47, 122.30, 121.94, 121.85, 118.63, 114.78, 69.51, 47.83, 32.41, 29.57, 26.04, 22.53, 13.40, 11.50, 11.41.

**21**
$^{1}$H NMR (300 MHz, DMSO-$d_6$) δ 10.27 (s, 1H), 7.97 (d, $J$ = 1.9 Hz, 1H), 7.92 (s, 1H), 7.66 (dd, $J$ = 8.0, 2.1 Hz, 1H), 7.52 (d, $J$ = 7.7 Hz, 1H), 7.46 – 7.26 (m, 12H), 7.31 – 7.11 (m, 3H), 6.93 (d, $J$ = 8.7 Hz, 2H), 5.45 (d, $J$ = 7.2 Hz, 1H), 5.06 (s, 2H), 2.01 – 1.75 (m, 2H), 1.67 – 1.55 (m, 4H). $^{13}$C NMR (75 MHz, DMSO-$d_6$) δ 138.62, 137.90, 135.19, 135.11, 130.03, 128.62, 128.41, 128.27, 127.79, 127.60, 126.61, 126.46, 118.70, 114.64, 100.31, 69.14, 58.90, 56.68, 36.19, 35.75, 23.24.

**22 (mixture of diastereomers)**
$^{1}$H NMR (300 MHz, DMSO-$d_6$) δ 11.32 (s, 1H), 10.36 (d, $J$ = 18.2 Hz, 1H), 8.69 (dd, $J$ = 16.5, 7.5 Hz, 1H), 8.13 (s, 1H), 8.09 – 7.98 (m, 2H), 7.98 – 7.84 (m, 2H), 7.69 (dd, $J$ = 32.3, 8.2 Hz, 1H), 7.56 – 7.11 (m, 16H), 6.96 (dd, $J$ = 34.8, 8.5 Hz, 2H), 5.52 (dd, $J$ = 30.7, 7.4 Hz, 1H), 5.05 (d, $J$ = 23.6 Hz, 2H), 4.16 – 3.95 (m, 1H), 1.44 – 1.34 (m, 3H). $^{13}$C NMR (75 MHz, DMSO-$d_6$) δ 173.72, 169.41, 168.20, 158.43, 141.09, 140.92, 138.73, 137.47, 135.54, 130.93, 130.28, 129.07, 128.95, 128.88, 128.84, 128.26, 128.22, 128.05, 128.01, 125.36, 124.12, 123.18, 122.61, 122.18, 120.77, 120.64, 120.01, 119.32, 119.13, 115.23, 115.12, 114.95, 112.71, 110.19, 109.99, 69.64, 56.74, 56.74, 44.99, 40.41, 40.14, 39.57, 19.50, 19.32.

**23 (mixture of diastereomers)**
$^{1}$H NMR (300 MHz, DMSO-$d_6$) δ 10.37 (d, $J$ = 15.8 Hz, 1H), 8.80 (t, $J$ = 7.2 Hz, 1H), 8.05 – 7.85 (m, 2H), 7.81 – 7.23 (m, 21H), 6.97 (dd, $J$ = 28.2, 8.4 Hz, 2H), 5.50 (dd, $J$ = 23.7, 7.4 Hz, 1H), 5.06 (d, $J$ = 16.8 Hz, 2H), 4.08 – 3.96 (m, 1H), 1.35 (dd, $J$ = 12.5, 7.0 Hz, 3H).

**24**
$^{1}$H NMR (300 MHz, DMSO-$d_6$) δ 10.37 (s, 1H), 8.89 (d, $J$ = 6.9 Hz, 1H), 8.53 – 8.43 (m, 1H), 8.00 (s, 1H), 7.90 (s, 1H), 7.83 – 7.68 (m, 3H), 7.54 – 7.25 (m, 13H), 7.00 (d, $J$ = 8.3 Hz, 2H), 5.54 (d, $J$ = 6.9 Hz, 1H), 5.07 (s, 2H), 3.48 – 3.24 (m, 2H), 2.31 (s, 2H), 1.40 (d, $J$ = 24.2 Hz, 10H). $^{13}$C NMR (126 MHz, DMSO-$d_6$) δ 171.63, 169.70, 168.21, 167.16, 158.49, 139.34, 137.44, 135.57, 135.30, 131.42, 130.25, 129.24, 129.05, 128.86, 128.69, 128.24, 128.05, 127.56, 122.51, 122.16, 119.12, 115.23, 69.62, 57.29, 40.47, 40.31, 40.19, 40.14, 39.97, 39.81, 39.64, 39.47, 37.81, 33.99, 33.82, 26.20, 21.59.

**25**
$^{1}$H NMR (300 MHz, DMSO-$d_6$) δ 10.31 (s, 1H), 8.36 (t, $J$ = 7.8 Hz, 2H), 8.01 (t, $J$ = 1.9 Hz, 1H), 7.90 (s, 1H), 7.77 – 7.67 (m, 1H), 7.54 – 7.47 (m, 1H), 7.46 – 7.25 (m, 15H), 6.99 (dd, $J$ = 9.0, 2.5 Hz, 4H), 5.57 – 5.16 (m, 2H), 5.09 (d, $J$ = 4.4 Hz, 5H), 1.39 – 1.28 (m, 2H), 1.20 – 1.04 (m, 27H). $^{13}$C NMR (126 MHz, DMSO-$d_6$) δ 172.51, 172.34, 170.82, 170.75, 169.63, 167.79, 157.94, 157.92, 157.60, 138.92, 137.10, 137.02, 135.11, 131.69, 130.35, 129.56, 128.94, 128.73, 128.56, 128.42, 128.40, 128.35, 127.78, 127.76, 127.57, 122.01, 121.67, 118.65, 114.69, 114.43, 69.16, 56.51, 55.54, 55.30, 35.46, 35.38, 35.36, 27.29, 27.26, 27.23, 23.55, 23.49, 16.56.

**26 (mixture of diastereomers)**
$^{1}$H NMR (300 MHz, DMSO-$d_6$) δ 10.41 (d, $J$ = 5.1 Hz, 1H), 8.87 (d, $J$ = 7.4 Hz, 1H), 8.06 – 8.00 (m, 1H), 7.93 (s, 1H), 7.79 – 7.70 (m, 1H), 7.53 (d, $J$ = 7.7 Hz, 1H), 7.48 – 7.29 (m, 10H), 7.16 – 6.99 (m, 4H), 6.88 – 6.70 (m, 2H), 5.56 (d, $J$ = 7.4 Hz, 1H), 5.10 (s, 2H), 4.41 – 4.24 (m, 1H), 3.88 (t, $J$ = 10.2 Hz, 1H), 3.11 – 2.81 (m, 3H). $^{13}$C NMR (75 MHz, DMSO-$d_6$) δ 171.86,

171.83, 169.52, 168.23, 158.52, 154.16, 139.26, 139.24, 137.46, 135.59, 130.31, 130.26, 129.13, 129.07, 128.88, 128.27, 128.05, 127.51, 122.63, 122.24, 121.84, 121.77, 120.72, 119.22, 116.56, 116.52, 115.28, 69.64, 67.62, 67.53, 60.60, 56.97, 38.22, 38.15, 28.15, 28.03.

**27**

$^{1}$H NMR (300 MHz, DMSO-$d_6$) δ 12.07 (s, 1H), 10.52 (s, 1H), 8.21 (d, $J$ = 7.8 Hz, 1H), 8.06 – 8.00 (m, 1H), 7.94 (s, 1H), 7.83 – 7.72 (m, 3H), 7.60 – 7.27 (m, 14H), 7.06 – 7.00 (m, 2H), 5.93 (s, 1H), 5.73 (d, $J$ = 7.8 Hz, 1H), 5.09 (s, 2H). $^{13}$C NMR (75 MHz, DMSO-$d_6$) δ 168.80, 167.68, 160.63, 158.15, 153.81, 145.04, 138.55, 138.13, 136.99, 135.17, 130.36, 130.27, 129.37, 129.05, 128.71, 128.43, 127.82, 127.61, 126.82, 122.45, 122.05, 121.94, 118.88, 114.93, 114.41, 87.46, 69.24, 55.88.

**28 (mixture of diastereomers)**

$^{1}$H NMR (300 MHz, DMSO-$d_6$) δ 10.34 (d, $J$ = 13.5 Hz, 1H), 8.65 (dd, $J$ = 15.4, 7.5 Hz, 1H), 7.97 (d, $J$ = 20.0 Hz, 1H), 7.92 – 7.84 (m, 1H), 7.68 (dd, $J$ = 21.9, 8.1 Hz, 1H), 7.49 (t, $J$ = 7.7 Hz, 1H), 7.45 – 7.12 (m, 16H), 6.96 (dd, $J$ = 21.2, 8.3 Hz, 2H), 5.49 (dd, $J$ = 27.3, 7.5 Hz, 1H), 5.05 (d, $J$ = 10.6 Hz, 2H), 3.95 – 3.82 (m, 1H), 1.29 (dd, $J$ = 10.4, 7.0 Hz, 3H). $^{13}$C NMR (75 MHz, DMSO-$d_6$) δ 173.72, 169.42, 168.22, 158.40, 142.49, 139.12, 137.42, 135.57, 130.28, 129.03, 128.95, 128.86, 128.85, 128.60, 128.57, 128.22, 128.04, 128.01, 127.78, 127.75, 126.87, 122.60, 122.18, 119.12, 115.21, 115.13, 69.60, 57.02, 44.49, 19.10.

**29**

$^{1}$H NMR (300 MHz, DMSO-$d_6$) δ 10.44 (s, 1H), 9.09 (d, $J$ = 6.8 Hz, 1H), 8.07 (t, $J$ = 2.1 Hz, 1H), 7.94 (s, 1H), 7.81 – 7.70 (m, 3H), 7.54 (d, $J$ = 7.7 Hz, 1H), 7.47 – 7.24 (m, 12H), 7.08 – 6.99 (m, 2H), 5.63 (d, $J$ = 6.8 Hz, 1H), 5.10 (s, 2H), 2.55 (s, 3H). $^{13}$C NMR (75 MHz, DMSO-$d_6$) δ 170.76, 169.30, 168.23, 161.88, 159.76, 158.67, 139.27, 137.43, 135.59, 130.75, 130.64, 129.51, 129.32, 129.12, 128.88, 128.28, 128.06, 122.62, 122.20, 119.20, 116.33, 116.04, 115.30, 69.64, 57.80, 12.44.

**30**

$^{1}$H NMR (300 MHz, Acetone-$d_6$) δ 9.73 (s, 1H), 8.13 (m, $J$ = 2.0 Hz, 1H), 7.86 (d, $J$ = 7.6 Hz, 1H), 7.68 – 7.55 (m, 2H), 7.51 – 7.42 (m, 4H), 7.41 – 7.27 (m, 4H), 7.04 – 6.96 (m, 2H), 6.65 (s, 1H), 5.68 (d, $J$ = 7.6 Hz, 1H), 5.11 (s, 2H), 2.45 – 2.23 (m, 1H), 1.94 – 1.68 (m, 4H), 1.63 (d, $J$ = 9.3 Hz, 1H), 1.54 – 1.38 (m, 1H), 1.26 (s, 4H). $^{13}$C NMR (75 MHz, Acetone-$d_6$) δ 170.54, 169.37, 167.98, 158.66, 139.14, 137.42, 132.30, 130.64, 128.76, 128.60, 128.37, 127.71, 127.48, 122.35, 121.98, 118.76, 114.81, 109.99, 69.51, 56.97, 44.33, 29.40, 25.68, 25.48.

**31**

$^{1}$H NMR (300 MHz, DMSO-$d_6$) δ 10.40 (s, 1H), 8.80 (d, $J$ = 7.6 Hz, 1H), 8.00 (s, 1H), 7.91 (s, 1H), 7.72 (d, $J$ = 7.9 Hz, 1H), 7.51 (d, $J$ = 7.7 Hz, 1H), 7.45 – 7.24 (m, 10H), 7.00 (d, $J$ = 8.3 Hz, 2H), 6.59 (s, 2H), 5.54 (d, $J$ = 7.6 Hz, 1H), 5.08 (s, 2H), 3.71 (s, 7H), 3.60 (s, 3H), 3.47 (s, 2H). $^{13}$C NMR (75 MHz, DMSO-$d_6$) δ 170.36, 169.66, 168.21, 158.46, 153.00, 139.23, 137.45, 135.60, 132.48, 130.52, 129.07, 128.87, 128.24, 128.02, 122.60, 122.15, 119.18, 115.22, 106.72, 69.63, 60.38, 56.99, 56.18.

**32**

$^{1}$H NMR (500 MHz, DMSO-$d_6$) δ 10.33 (s, 1H), 8.45 (d, $J$ = 7.5 Hz, 1H), 8.01 – 7.97 (m, 1H), 7.91 (s, 1H), 7.74 – 7.68 (m, 1H), 7.53 – 7.47 (m, 1H), 7.43 – 7.26 (m, 8H), 7.02 – 6.96 (m, 2H), 5.51 (d, $J$ = 7.5 Hz, 1H), 5.07 (s, 2H), 2.07 (d, $J$ = 6.8 Hz, 2H), 1.67 – 1.52 (m, 7H), 1.23 – 1.05 (m, 3H), 0.94 – 0.82 (m, 2H). $^{13}$C NMR (126 MHz, DMSO-$d_6$) δ 171.84, 169.80, 168.23,

158.39, 139.32, 137.46, 135.56, 130.61, 129.11, 129.04, 128.86, 128.24, 128.04, 122.51, 122.15, 119.13, 115.16, 69.61, 56.93, 43.04, 35.28, 32.96, 32.92, 26.33, 26.08.

**33**
$^{1}$H NMR (300 MHz, DMSO-$d_6$) δ 10.39 (s, 1H), 8.84 (d, J = 7.0 Hz, 1H), 8.51 (d, J = 2.6 Hz, 1H), 8.02 (s, 1H), 7.96 (dd, J = 9.6, 2.6 Hz, 1H), 7.92 (s, 1H), 7.76 (d, J = 7.7 Hz, 1H), 7.55 – 7.30 (m, 15H), 7.01 (d, J = 8.5 Hz, 2H), 6.56 – 6.49 (m, 1H), 5.67 (d, J = 6.9 Hz, 1H), 5.09 (s, 2H). $^{13}$C NMR (APT, 75 MHz, DMSO-$d_6$) δ 169.67, 168.24, 163.96, 161.54, 158.55, 141.68, 140.96, 139.70, 139.32, 137.47, 135.57, 129.86, 129.82, 129.68, 129.57, 129.05, 128.99, 128.88, 128.25, 128.01, 127.43, 122.58, 122.25, 119.86, 119.25, 115.20, 114.86, 112.49, 69.60, 57.90.

**34**
$^{1}$H NMR (300 MHz, DMSO-$d_6$) δ 10.37 (s, 1H), 8.53 (d, $J$ = 7.6 Hz, 1H), 8.04 (d, $J$ = 11.3 Hz, 1H), 7.90 (d, $J$ = 8.5 Hz, 2H), 7.73 (d, $J$ = 8.3 Hz, 1H), 7.63 (t, $J$ = 6.8 Hz, 1H), 7.52 (d, $J$ = 7.8 Hz, 1H), 7.47 – 7.24 (m, 7H), 7.01 (d, $J$ = 8.3 Hz, 1H), 5.52 (d, $J$ = 7.4 Hz, 1H), 5.09 (s, 1H), 3.86 (t, $J$ = 11.8 Hz, 1H), 2.25 (d, $J$ = 8.1 Hz, 2H), 1.71 (d, $J$ = 17.3 Hz, 1H), 1.50 (dt, $J$ = 13.2, 6.6 Hz, 1H), 1.36 (q, $J$ = 7.4 Hz, 1H), 1.18 (t, $J$ = 7.2 Hz, 1H), 0.86 (d, $J$ = 6.5 Hz, 4H).

**35**
$^{1}$H NMR (300 MHz, DMSO-$d_6$) δ 10.46 (s, 1H), 9.02 (d, $J$ = 7.0 Hz, 1H), 8.81 – 8.73 (m, 2H), 8.08 – 8.02 (m, 1H), 8.01 – 7.96 (m, 2H), 7.94 (s, 1H), 7.81 – 7.72 (m, 1H), 7.58 – 7.30 (m, 12H), 7.09 – 7.01 (m, 2H), 5.68 (d, $J$ = 7.0 Hz, 1H), 5.11 (s, 2H), 2.66 (s, 3H). $^{13}$C NMR (75 MHz, DMSO-$d_6$) δ 169.16, 168.21, 163.42, 161.15, 158.66, 156.13, 150.29, 140.43, 139.24, 137.45, 135.61, 129.65, 129.57, 129.12, 129.01, 128.89, 128.28, 128.06, 122.68, 122.24, 120.92, 119.22, 115.28, 69.64, 58.02, 17.53.

**36**
$^{1}$H NMR (300 MHz, DMSO-$d_6$) δ 10.32 (s, 1H), 7.99 (s, 2H), 7.90 (s, 1H), 7.72 (d, J = 8.0 Hz, 1H), 7.51 (d, J = 7.7 Hz, 1H), 7.44 – 7.28 (m, 14H), 6.97 (d, J = 8.5 Hz, 2H), 5.31 (d, J = 7.9 Hz, 1H), 5.06 (s, 2H), 5.03 (s, 2H).

**37**
$^{1}$H NMR (300 MHz, DMSO-$d_6$) δ 10.35 (s, 1H), 8.47 (d, $J$ = 7.6 Hz, 1H), 8.00 (d, $J$ = 1.9 Hz, 1H), 7.91 (s, 1H), 7.77 – 7.69 (m, 1H), 7.51 (d, $J$ = 7.8 Hz, 1H), 7.46 – 7.26 (m, 9H), 7.05 – 6.94 (m, 2H), 5.53 (d, $J$ = 7.5 Hz, 1H), 5.08 (s, 2H), 2.17 (t, $J$ = 7.4 Hz, 2H), 1.55 – 1.41 (m, 3H), 1.24 – 1.05 (m, 2H), 0.82 (d, $J$ = 6.6 Hz, 6H). $^{13}$C NMR (75 MHz, DMSO-$d_6$) δ 172.59, 169.80, 168.23, 158.39, 139.30, 137.47, 135.57, 130.69, 129.09, 128.87, 128.24, 128.03, 122.57, 122.21, 119.17, 115.17, 69.58, 56.89, 38.43, 35.46, 27.66, 22.91.

**38 (mixture of diastereomers)**
$^{1}$H NMR (300 MHz, DMSO-$d_6$) δ 10.43 (d, $J$ = 6.5 Hz, 1H), 8.96 (dd, $J$ = 47.3, 7.5 Hz, 1H), 8.05 (dt, $J$ = 8.9, 1.9 Hz, 1H), 7.93 (s, 1H), 7.83 – 7.71 (m, 1H), 7.59 – 7.51 (m, 1H), 7.50 – 7.20 (m, 12H), 7.08 – 6.94 (m, 3H), 5.60 (dd, $J$ = 18.8, 7.5 Hz, 1H), 5.10 (s, 2H), 3.91 – 3.78 (m, 3H), 2.35 – 2.20 (m, 4H). $^{13}$C NMR (75 MHz, DMSO-$d_6$) δ 171.44, 171.24, 169.40, 169.19, 168.83, 168.22, 158.50, 139.58, 139.51, 139.21, 139.05, 138.46, 137.47, 135.69, 130.75, 130.16, 129.02, 128.89, 128.27, 128.06, 125.59, 122.33, 120.75, 119.32, 117.41, 115.30, 69.66, 56.60, 49.83, 47.29, 22.36, 21.63.

**39**

[1]H NMR (300 MHz, DMSO-$d_6$) δ 10.36 (s, 1H), 8.56 (d, *J* = 7.5 Hz, 1H), 8.44 (t, *J* = 5.6 Hz, 1H), 8.03 – 7.97 (m, 1H), 7.94 – 7.88 (m, 1H), 7.85 – 7.79 (m, 2H), 7.73 (dd, *J* = 7.9, 2.2 Hz, 1H), 7.60 – 7.12 (m, 14H), 6.99 (dd, *J* = 9.1, 2.7 Hz, 2H), 5.53 (d, *J* = 7.5 Hz, 1H), 5.07 (s, 2H), 3.29 – 3.21 (m, 2H), 2.27 (t, *J* = 7.3 Hz, 2H), 1.74 (p, *J* = 7.3 Hz, 2H). [13]C NMR (75 MHz, DMSO-$d_6$) δ 172.37, 169.81, 168.24, 166.63, 158.43, 139.30, 137.48, 135.58, 135.11, 131.44, 130.56, 129.14, 129.04, 128.87, 128.65, 128.25, 128.04, 127.59, 122.58, 122.24, 119.21, 115.21, 114.86, 69.62, 57.04, 39.32, 34.17, 32.93, 25.86.

**40**

[1]H NMR (500 MHz, DMSO-$d_6$) δ 10.37 (d, *J* = 4.6 Hz, 1H), 8.68 (t, *J* = 6.2 Hz, 1H), 8.61 (t, *J* = 7.0 Hz, 1H), 7.98 (s, 1H), 7.94 – 7.84 (m, 3H), 7.72 (d, *J* = 8.0 Hz, 1H), 7.56 – 7.27 (m, 13H), 7.04 – 6.97 (m, 2H), 5.49 (dd, *J* = 7.1, 2.8 Hz, 1H), 5.08 (s, 2H), 4.76 – 4.60 (m, 1H), 2.95 – 2.73 (m, 2H), 2.55 (d, *J* = 3.3 Hz, 2H), 2.25 – 2.02 (m, 2H). [13]C NMR (126 MHz, DMSO-$d_6$) δ 171.28, 169.50, 168.17, 166.97, 158.56, 139.19, 137.43, 135.58, 134.39, 131.87, 130.05, 129.18, 128.87, 128.69, 128.25, 128.05, 127.95, 122.65, 122.18, 119.15, 115.26, 69.64, 57.14, 52.79, 52.56, 50.46, 49.80, 38.58, 25.68, 25.25.

**41**

[1]H NMR (500 MHz, Acetone-$d_6$) δ 9.77 (s, 1H), 8.17 – 8.13 (m, 1H), 7.90 – 7.84 (m, 1H), 7.81 (d, *J* = 7.6 Hz, 1H), 7.60 (dt, *J* = 7.8, 1.3 Hz, 1H), 7.50 – 7.43 (m, 4H), 7.41 – 7.28 (m, 4H), 7.03 – 6.96 (m, 2H), 6.80 (dt, *J* = 15.4, 7.0 Hz, 1H), 6.22 – 6.15 (m, 1H), 5.80 – 5.76 (m, 1H), 5.11 (s, 2H), 2.15 (dq, *J* = 7.1, 1.6 Hz, 2H), 1.46 (hept, *J* = 7.4 Hz, 2H), 0.91 (t, *J* = 7.4 Hz, 3H).[13]C NMR (126 MHz, Acetone-$d_6$) δ 173.02, 171.71, 169.13, 158.70, 143.87, 139.06, 137.40, 130.46, 128.89, 128.63, 128.37, 127.71, 127.47, 123.93, 122.34, 121.92, 118.72, 114.85, 69.51, 57.03, 33.67, 21.31, 13.04.

**42**

[1]H NMR (300 MHz, DMSO-$d_6$) δ 10.41 (s, 1H), 8.90 (d, *J* = 7.3 Hz, 1H), 8.07 – 8.00 (m, 1H), 7.92 (dd, *J* = 6.4, 2.6 Hz, 3H), 7.77 (d, *J* = 8.0 Hz, 1H), 7.58 – 7.27 (m, 16H), 7.03 (d, *J* = 8.7 Hz, 2H), 5.75 (d, *J* = 7.3 Hz, 1H), 5.11 (s, 3H). [13]C NMR (75 MHz, DMSO-$d_6$) δ 169.22, 167.78, 166.40, 158.07, 138.88, 137.04, 135.12, 133.79, 131.40, 129.19, 128.42, 128.13, 127.77, 127.58, 121.78, 114.71, 69.15.

**43**

[1]H NMR (300 MHz, Acetone-$d_6$) δ 9.69 (s, 1H), 8.16 – 8.12 (m, 1H), 7.91 – 7.83 (m, 1H), 7.75 (d, *J* = 7.6 Hz, 1H), 7.64 – 7.56 (m, 1H), 7.51 – 7.27 (m, 8H), 7.04 (d, 2H), 6.97 (d, 2H), 6.71 (d, 2H), 5.71 (d, *J* = 7.5 Hz, 1H), 5.11 (s, 2H), 2.88 – 2.79 (m, 2H), 2.63 – 2.54 (m, 2H). [13]C NMR (75 MHz, Acetone-$d_6$) δ 171.56, 169.19, 158.64, 155.56, 139.11, 137.41, 135.15, 132.03, 130.44, 129.22, 128.76, 128.65, 128.38, 127.71, 127.49, 122.35, 122.09, 118.89, 115.08, 114.80, 69.51, 57.15, 37.69, 30.60.

**44**

[1]H NMR (300 MHz, DMSO-$d_6$) δ 10.34 (s, 1H), 8.08 (d, *J* = 7.3 Hz, 1H), 8.02 (d, *J* = 2.4 Hz, 1H), 7.92 (s, 1H), 7.74 (d, *J* = 8.1 Hz, 1H), 7.52 (d, *J* = 7.7 Hz, 1H), 7.47 – 7.28 (m, 14H), 7.04 – 6.94 (m, 3H), 6.40 – 6.26 (m, 2H), 5.57 (d, *J* = 7.3 Hz, 1H), 5.09 (s, 2H), 2.09 (q, *J* = 7.2 Hz, 3H), 1.76 (d, *J* = 7.5 Hz, 5H), 1.48 – 1.33 (m, 3H), 0.89 (t, *J* = 7.3 Hz, 5H). [13]C NMR (126 MHz, DMSO-$d_6$) δ 169.34, 168.39, 167.75, 157.98, 138.82, 137.02, 135.70, 135.10, 130.39,

130.02, 129.20, 128.90, 128.57, 128.41, 127.78, 127.56, 122.11, 121.74, 118.73, 114.68, 114.59, 69.15, 56.92, 29.81, 21.52, 13.78, 12.65.

**45**
$^{1}$H NMR (500 MHz, DMSO-$d_6$) δ 10.38 (s, 1H), 8.79 (d, $J$ = 7.6 Hz, 1H), 7.98 (t, $J$ = 2.0 Hz, 1H), 7.90 (s, 1H), 7.73 – 7.68 (m, 1H), 7.53 – 7.48 (m, 1H), 7.43 – 7.23 (m, 11H), 7.21 – 7.16 (m, 1H), 7.08 – 6.93 (m, 3H), 6.83 – 6.76 (m, 1H), 5.52 (d, $J$ = 7.6 Hz, 1H), 5.07 (s, 2H), 3.55 (s, 2H). $^{13}$C NMR (126 MHz, DMSO-$d_6$) δ 170.41, 169.63, 168.21, 158.45, 139.22, 137.45, 136.82, 135.57, 130.52, 129.81, 129.50, 129.05, 128.87, 128.59, 128.25, 128.04, 126.73, 122.60, 122.20, 119.17, 115.23, 114.85, 69.62, 67.23, 56.98, 42.10.

**46 (mixture of diastereomers)**
$^{1}$H NMR (500 MHz, DMSO-$d_6$) δ 10.34 (d, $J$ = 3.6 Hz, 1H), 8.47 (dd, $J$ = 7.6, 3.0 Hz, 1H), 8.01 – 7.97 (m, 1H), 7.90 (s, 1H), 7.75 – 7.69 (m, 1H), 7.52 – 7.47 (m, 1H), 7.43 – 7.26 (m, 9H), 7.02 – 6.95 (m, 2H), 5.53 (dd, $J$ = 9.1, 7.5 Hz, 1H), 5.07 (s, 2H), 2.15 (dt, $J$ = 13.5, 5.7 Hz, 1H), 2.07 – 1.97 (m, 1H), 1.83 (s, 1H), 1.32 – 1.14 (m, 2H), 1.11 – 1.02 (m, 1H), 0.87 – 0.75 (m, 6H). $^{13}$C NMR (126 MHz, DMSO-$d_6$) δ 172.14, 169.80, 169.77, 168.23, 158.39, 139.33, 137.46, 135.57, 130.58, 129.13, 129.07, 129.03, 128.86, 128.24, 128.03, 122.53, 122.14, 119.11, 115.15, 109.99, 69.61, 56.87, 44.20, 42.83, 40.47, 40.30, 40.13, 40.08, 39.97, 39.91, 39.86, 39.80, 39.63, 39.46, 39.02, 30.41, 22.66, 22.05, 19.89, 14.60.

**47**
$^{1}$H NMR (300 MHz, DMSO-$d_6$) δ 10.34 (s, 1H), 9.20 (d, $J$ = 7.5 Hz, 1H), 7.98 (t, $J$ = 1.9 Hz, 1H), 7.90 (s, 1H), 7.76 – 7.66 (m, 1H), 7.51 (dd, $J$ = 7.7, 1.6 Hz, 1H), 7.44 – 7.26 (m, 9H), 7.03 – 6.94 (m, 2H), 5.52 (d, $J$ = 7.5 Hz, 1H), 5.07 (s, 2H), 2.30 (t, $J$ = 6.9 Hz, 2H), 1.49 (h, $J$ = 7.2 Hz, 2H), 0.94 (t, $J$ = 7.3 Hz, 3H). $^{13}$C NMR (75 MHz, DMSO-$d_6$) δ 169.04, 168.20, 158.56, 152.78, 139.20, 137.43, 135.57, 129.51, 129.49, 129.06, 128.86, 128.24, 128.03, 122.62, 122.19, 119.15, 115.17, 87.68, 76.49, 69.62, 57.28, 21.29, 20.18, 13.76.

**48**
$^{1}$H NMR (300 MHz, DMSO-$d_6$) δ 10.33 (s, 1H), 8.51 (d, $J$ = 7.5 Hz, 1H), 7.98 (t, $J$ = 1.9 Hz, 1H), 7.89 (s, 1H), 7.74 – 7.67 (m, 1H), 7.52 – 7.46 (m, 1H), 7.43 – 7.24 (m, 9H), 7.02 – 6.94 (m, 2H), 5.50 (d, $J$ = 7.5 Hz, 1H), 5.06 (s, 2H), 3.60 (t, $J$ = 6.3 Hz, 2H), 2.22 (t, $J$ = 7.0 Hz, 2H), 1.74 – 1.51 (m, 4H). $^{13}$C NMR (75 MHz, DMSO-$d_6$) δ 172.23, 169.76, 168.23, 158.42, 139.29, 137.46, 135.57, 130.57, 129.11, 129.04, 128.87, 128.25, 128.03, 122.56, 122.19, 119.17, 115.20, 69.62, 56.95, 45.55, 34.29, 31.96, 23.06.

**49**
$^{1}$H NMR (300 MHz, Acetone-$d_6$) δ 9.72 (s, 1H), 8.08 (t, $J$ = 1.8 Hz, 1H), 7.84 – 7.77 (m, 2H), 7.60 – 7.55 (m, 1H), 7.49 – 7.28 (m, 8H), 7.20 – 7.14 (m, 2H), 7.00 – 6.94 (m, 2H), 6.80 – 6.74 (m, 2H), 5.73 – 5.67 (m, 1H), 5.10 (s, 2H), 3.56 (s, 2H). $^{13}$C NMR (75 MHz, Acetone-$d_6$) δ 170.74, 168.20, 158.71, 156.17, 139.01, 137.38, 135.07, 130.33, 130.26, 128.73, 128.63, 128.37, 127.72, 127.50, 126.65, 122.40, 122.06, 118.77, 115.17, 114.85, 69.51, 57.22, 41.61.

**50**
$^{1}$H NMR (300 MHz, Acetone-$d_6$) δ 9.72 (s, 1H), 8.13 (s, 1H), 7.87 (d, J = 8.4 Hz, 1H), 7.75 (d, J = 7.3 Hz, 1H), 7.59 (d, J = 7.7 Hz, 1H), 7.49 – 7.42 (m, 4H), 7.41 – 7.29 (m, 4H), 7.03 – 6.96 (m, 2H), 6.71 (s, 1H), 5.88 – 5.74 (m, 1H), 5.74 – 5.68 (m, 1H), 5.11 (s, 2H), 5.04 – 4.88 (m, 2H), 2.33 (t, J = 7.4 Hz, 2H), 2.09 – 2.07 (m, 1H), 1.70 (p, J = 7.4 Hz, 2H).

**51**

$^1$H NMR (300 MHz, DMSO-$d_6$) δ 10.36 (s, 1H), 8.50 (d, $J$ = 7.6 Hz, 1H), 8.01 (t, $J$ = 1.9 Hz, 1H), 7.92 (s, 1H), 7.78 – 7.69 (m, 1H), 7.52 (dt, $J$ = 7.8, 1.3 Hz, 1H), 7.48 – 7.24 (m, 10H), 7.00 (d, $J$ = 8.7 Hz, 2H), 5.53 (d, $J$ = 7.6 Hz, 1H), 5.09 (s, 2H), 2.27 – 2.11 (m, 2H), 1.59 – 1.39 (m, 2H), 1.43 – 1.32 (m, 1H), 0.85 (d, $J$ = 6.4 Hz, 7H). $^{13}$C NMR (75 MHz, DMSO-$d_6$) δ 172.32, 169.38, 167.79, 157.94, 138.85, 137.02, 135.12, 130.19, 128.64, 128.41, 127.79, 127.58, 122.09, 121.73, 118.71, 114.72, 69.15, 56.42, 34.26, 32.92, 27.24, 22.29.

**52**

$^1$H NMR (300 MHz, Acetone-$d_6$) δ 10.32 (s, 1H), 8.63 (s, 1H), 8.58 (t, $J$ = 1.8 Hz, 1H), 8.38 – 8.28 (m, 1H), 8.08 – 7.98 (m, 1H), 7.97 – 7.86 (m, 4H), 7.93 – 7.70 (m, 4H), 7.44 (d, $J$ = 8.7 Hz, 2H), 6.19 (s, 1H), 5.56 (s, 2H), 3.40 – 3.23 (m, 1H), 2.43 – 2.18 (m, 3H), 2.20 – 2.03 (m, 2H), 2.07 – 1.91 (m, 1H). $^{13}$C NMR (75 MHz, Acetone-$d_6$) δ 176.35, 174.57, 170.26, 159.58, 140.01, 138.33, 136.04, 131.43, 129.73, 129.51, 129.29, 128.63, 128.40, 123.22, 122.85, 119.57, 115.73, 70.43, 58.01, 45.36, 31.02, 26.74.

**53**

$^1$H NMR (300 MHz, DMSO-$d_6$) δ 10.36 (s, 1H), 8.54 (d, $J$ = 7.5 Hz, 1H), 8.06 – 7.98 (m, 2H), 7.91 (s, 1H), 7.73 (dd, $J$ = 7.9, 2.2 Hz, 1H), 7.51 (d, $J$ = 7.7 Hz, 1H), 7.44 – 7.26 (m, 9H), 7.04 – 6.96 (m, 2H), 5.52 (d, $J$ = 7.4 Hz, 1H), 5.08 (s, 2H), 3.15 – 2.92 (m, 2H), 2.27 – 2.11 (m, 2H), 1.62 (p, $J$ = 7.2 Hz, 2H), 1.54 – 1.44 (m, 1H), 0.70 – 0.53 (m, 4H). $^{13}$C NMR (75 MHz, DMSO-$d_6$) δ 172.81, 172.29, 169.82, 168.25, 158.44, 139.30, 137.48, 135.58, 130.54, 129.15, 129.04, 128.88, 128.25, 128.04, 122.58, 122.22, 119.20, 115.21, 69.62, 57.03, 38.74, 32.86, 26.05, 14.04, 6.52.

**54**

$^1$H NMR (300 MHz, DMSO-$d_6$) δ 10.37 (s, 1H), 8.55 (d, $J$ = 7.6 Hz, 1H), 8.02 (t, $J$ = 1.9 Hz, 1H), 7.92 (s, 1H), 7.73 (d, $J$ = 8.0 Hz, 1H), 7.52 (d, $J$ = 7.7 Hz, 1H), 7.47 – 7.28 (m, 14H), 7.00 (d, $J$ = 8.7 Hz, 2H), 5.53 (d, $J$ = 7.5 Hz, 1H), 5.09 (s, 2H), 3.80 – 3.69 (m, 1H), 2.74 – 2.58 (m, 1H), 1.89 – 1.38 (m, 2H). $^{13}$C NMR (75 MHz, DMSO-$d_6$) δ 173.77, 169.24, 168.97, 167.78, 157.97, 138.81, 137.01, 136.30, 135.12, 135.12, 130.07, 129.36, 128.61, 128.42, 127.80, 127.58, 126.63, 121.73, 118.73, 114.74, 69.16.

**55**

$^1$H NMR (300 MHz, Acetone-$d_6$) δ 9.76 (s, 1H), 8.14 – 8.10 (m, 1H), 7.90 – 7.83 (m, 1H), 7.72 (s, 1H), 7.64 – 7.55 (m, 1H), 7.49 – 7.41 (m, 4H), 7.41 – 7.27 (m, 4H), 7.04 – 6.94 (m, 2H), 6.76 (s, 1H), 5.76 – 5.71 (m, 1H), 5.11 (s, 2H), 3.36 – 3.21 (m, 1H), 2.37 – 2.09 (m, 3H), 2.01 – 1.73 (m, 1H). $^{13}$C NMR (75 MHz, Acetone-$d_6$) δ 173.90, 169.24, 168.00, 158.69, 139.04, 137.40, 135.13, 130.44, 128.82, 128.64, 128.37, 127.71, 127.48, 122.31, 121.91, 118.70, 114.84, 69.51, 56.97, 39.07, 24.78, 17.92.

**56**

$^1$H NMR (300 MHz, DMSO-$d_6$) δ 10.47 (s, 1H), 9.58 (d, $J$ = 7.0 Hz, 1H), 8.33 (dd, $J$ = 4.9, 1.9 Hz, 1H), 8.01 (t, $J$ = 1.9 Hz, 1H), 7.98 – 7.91 (m, 2H), 7.82 – 7.74 (m, 1H), 7.58 – 7.52 (m, 1H), 7.51 – 7.27 (m, 10H), 7.10 – 7.01 (m, 3H), 5.69 (d, $J$ = 7.0 Hz, 1H), 5.10 (s, 2H), 3.69 – 3.48 (m, 4H), 3.17 (t, $J$ = 4.7 Hz, 4H). $^{13}$C NMR (75 MHz, DMSO-$d_6$) δ 169.28, 168.18, 166.08, 158.65, 148.58, 139.81, 139.18, 137.42, 135.61, 130.01, 129.36, 128.87, 128.26, 128.04, 122.67, 119.24, 117.41, 115.37, 69.65, 66.18, 57.24, 50.41.

**57**
$^1$H NMR (500 MHz, DMSO-$d_6$) δ 10.43 (s, 1H), 9.14 (d, $J$ = 6.9 Hz, 1H), 8.13 (s, 3H), 7.98 (t, $J$ = 1.9 Hz, 1H), 7.92 (s, 1H), 7.74 – 7.69 (m, 1H), 7.55 – 7.50 (m, 1H), 7.43 – 7.26 (m, 9H), 7.07 – 7.01 (m, 2H), 5.52 (d, $J$ = 6.8 Hz, 1H), 5.09 (s, 2H), 3.95 (t, $J$ = 6.4 Hz, 1H), 2.56 (t, $J$ = 7.9 Hz, 2H), 2.05 (s, 3H), 2.04 – 1.97 (m, 2H). $^{13}$C NMR (126 MHz, DMSO-$d_6$) δ 168.96, 168.43, 168.11, 158.72, 139.13, 137.39, 135.59, 129.40, 129.37, 129.14, 128.89, 128.28, 128.01, 122.67, 122.15, 119.17, 115.42, 69.62, 57.28, 51.88, 31.65, 28.54, 14.99.

**58**
$^1$H NMR (300 MHz, DMSO-$d_6$) δ 10.41 (s, 1H), 8.94 (d, $J$ = 6.8 Hz, 1H), 8.02 (t, $J$ = 1.9 Hz, 1H), 7.96 – 7.86 (m, 4H), 7.72 (dd, $J$ = 7.7, 2.2 Hz, 1H), 7.58 – 7.49 (m, 1H), 7.47 – 7.25 (m, 9H), 7.03 (d, $J$ = 8.7 Hz, 2H), 5.48 (d, $J$ = 6.7 Hz, 1H), 5.10 (s, 2H), 2.86 (d, $J$ = 5.7 Hz, 2H), 1.48 – 1.39 (m, 8H), 1.39 – 1.20 (m, 3H). $^{13}$C NMR (75 MHz, DMSO-$d_6$) δ 171.22, 168.84, 167.74, 158.15, 138.81, 136.97, 135.11, 129.40, 128.82, 128.64, 128.44, 127.83, 127.59, 122.14, 121.74, 114.86, 69.17, 56.93, 35.18, 32.81, 20.56.

**59**
$^1$H NMR (300 MHz, DMSO-$d_6$) δ 10.43 (s, 1H), 9.14 (d, $J$ = 7.2 Hz, 1H), 8.12 – 8.02 (m, 3H), 8.00 – 7.84 (m, 3H), 7.81 – 7.73 (m, 1H), 7.57 – 7.28 (m, 12H), 7.07 – 7.00 (m, 2H), 5.75 (d, $J$ = 7.2 Hz, 1H), 5.11 (s, 2H). $^{13}$C NMR (75 MHz, DMSO-$d_6$) δ 169.48, 168.23, 165.97, 158.60, 146.87, 139.31, 137.56, 137.15, 137.15, 135.59, 135.59, 129.81, 129.72, 129.08, 128.95, 128.89, 128.27, 128.04, 125.88, 122.25, 119.23, 115.21, 69.62, 57.79.

**60**
$^1$H NMR (300 MHz, DMSO-$d_6$) δ 10.35 (s, 1H), 8.54 (d, $J$ = 7.4 Hz, 1H), 8.03 – 7.97 (m, 1H), 7.94 – 7.88 (m, 1H), 7.77 – 7.68 (m, 1H), 7.54 – 7.48 (m, 1H), 7.44 – 7.26 (m, 9H), 7.03 – 6.95 (m, 2H), 5.51 (d, $J$ = 7.4 Hz, 1H), 5.08 (s, 2H), 2.76 (t, $J$ = 2.6 Hz, 1H), 2.29 (dt, 2H), 2.14 (dt, $J$ = 7.2, 2.7 Hz, 2H), 1.66 (p, $J$ = 7.3 Hz, 2H). $^{13}$C NMR (75 MHz, DMSO-$d_6$) δ 171.98, 169.77, 168.24, 158.43, 139.30, 137.47, 135.57, 130.51, 129.16, 129.04, 128.87, 128.25, 128.04, 122.57, 122.20, 119.17, 115.20, 84.60, 71.91, 69.63, 57.00, 34.15, 24.78, 17.85.

**61**
$^1$H NMR (300 MHz, DMSO-$d_6$) δ 10.36 (d, $J$ = 2.6 Hz, 1H), 8.77 – 8.65 (m, 1H), 7.98 (d, $J$ = 2.0 Hz, 1H), 7.89 (s, 1H), 7.70 (d, $J$ = 8.1 Hz, 1H), 7.53 – 7.45 (m, 1H), 7.43 – 7.25 (m, 10H), 7.01 – 6.95 (m, 2H), 5.58 – 5.49 (m, 1H), 5.06 (s, 2H), 4.48 – 4.35 (m, 1H), 2.80 – 2.58 (m, 2H), 1.45 (dd, $J$ = 9.9, 6.5 Hz, 3H).

**62**
$^1$H NMR (300 MHz, DMSO-$d_6$) δ 10.31 (s, 1H), 8.62 (d, $J$ = 19.3, 8.1 Hz, 1H), 8.05 (t, $J$ = 1.9 Hz, 1H), 7.93 (s, 1H), 7.91 – 7.82 (m, 2H), 7.82 – 7.76 (m, 1H), 7.57 – 7.50 (m, 1H), 7.46 – 7.25 (m, 11H), 7.24 – 7.15 (m, 2H), 7.09 – 6.97 (m, 4H), 6.95 – 6.88 (m, 2H), 5.02 (s, 2H), 4.82 – 4.72 (m, 1H), 3.13 – 2.91 (m, 2H). $^{13}$C NMR (75 MHz, DMSO-$d_6$) δ 171.18, 168.27, 166.20, 159.95, 157.39, 156.07, 139.37, 137.59, 135.54, 130.72, 130.67, 130.16, 129.06, 129.02, 128.81, 128.19, 128.14, 124.70, 122.53, 122.44, 119.88, 119.40, 117.79, 114.85, 69.55.

**63**
$^1$H NMR (500 MHz, DMSO-$d_6$) δ 10.26 (s, 1H), 9.20 (s, 1H), 8.53 (d, $J$ = 21.4, 8.1 Hz, 1H), 8.04 (t, $J$ = 2.0 Hz, 1H), 7.92 (s, 1H), 7.85 – 7.80 (m, 2H), 7.80 – 7.73 (m, 2H), 7.69 – 7.61 (m, 2H), 7.58 (s, 1H), 7.53 (d, $J$ = 7.7 Hz, 1H), 7.45 – 7.28 (m, 7H), 7.23 – 7.15 (m, 2H), 7.08 –

7.02 (m, 3H), 6.98 – 6.92 (m, 2H), 6.72 – 6.67 (m, 1H), 4.79 – 4.46 (m, 1H), 4.01 – 3.88 (m, 2H), 3.04 – 2.84 (m, 2H). $^{13}$C NMR (126 MHz, cdcl$_3$) δ 170.77, 167.84, 165.70, 159.48, 155.67, 153.53, 138.95, 138.87, 135.08, 133.04, 131.46, 131.44, 130.23, 129.68, 129.54, 128.67, 128.54, 128.25, 128.07, 127.63, 127.43, 127.34, 127.26, 126.54, 126.30, 125.82, 125.11, 124.25, 121.98, 119.44, 119.41, 118.95, 117.35, 117.30, 114.87, 56.17, 36.46, 35.60.

**64**
$^1$H NMR (300 MHz, DMSO-*d*$_6$) δ 10.17 (s, 1H), 8.55 – 8.46 (m, 1H), 8.04 (t, *J* = 2.0 Hz, 1H), 7.98 – 7.86 (m, 3H), 7.77 (dt, *J* = 7.7, 2.1 Hz, 1H), 7.51 (dt, *J* = 7.8, 1.4 Hz, 1H), 7.46 – 7.26 (m, 4H), 7.24 – 7.13 (m, 1H), 7.05 (dd, *J* = 11.7, 8.2, 1.7 Hz, 5H), 4.71 – 4.36 (m, 1H), 1.84 – 1.51 (m, 10H), 1.48 – 1.31 (m, 1H), 1.25 – 1.04 (m, 4H), 1.02 – 0.78 (m, 2H). $^{13}$C NMR (75 MHz, DMSO-*d*$_6$) δ 172.19, 168.31, 166.22, 159.92, 156.16, 139.50, 135.50, 130.67, 130.27, 129.19, 128.96, 124.67, 122.43, 119.83, 119.42, 117.85, 50.19, 34.30, 33.67, 32.17, 26.28, 26.06.

**65**
$^1$H NMR (300 MHz, DMSO-*d*$_6$) δ 10.38 (s, 1H), 8.73 (d, *J* = 19.4, 8.1 Hz, 1H), 8.10 – 8.04 (m, 1H), 7.93 (s, 1H), 7.91 – 7.75 (m, 7H), 7.63 – 7.51 (m, 2H), 7.50 – 7.30 (m, 6H), 7.18 (t, *J* = 7.4 Hz, 1H), 7.04 (d, *J* = 7.9 Hz, 2H), 6.99 (d, *J* = 8.5 Hz, 2H), 5.01 – 4.67 (m, 1H), 3.39 – 3.17 (m, 2H). $^{13}$C NMR (75 MHz, DMSO-*d*$_6$) δ 171.08, 168.29, 166.25, 159.98, 156.07, 139.36, 136.35, 135.56, 133.40, 132.26, 130.67, 130.15, 130.00, 129.04, 129.00, 128.28, 127.94, 127.78, 126.49, 125.91, 124.70, 123.72, 122.48, 119.88, 119.45, 117.78, 56.35, 37.48.

**66**
$^1$H NMR (300 MHz, DMSO-*d*$_6$) δ 10.20 (s, 1H), 8.68 – 8.55 (m, 1H), 8.04 (t, *J* = 1.9 Hz, 1H), 8.01 – 7.87 (m, 3H), 7.81 – 7.72 (m, 1H), 7.51 (d, *J* = 7.7 Hz, 1H), 7.46 – 7.12 (m, 9H), 7.06 (dd, *J* = 8.3 Hz, 4H), 4.66 – 4.20 (m, 1H), 2.85 – 2.56 (m, 1H), 2.18 – 2.00 (m, 2H). $^{13}$C NMR (75 MHz, DMSO-*d*$_6$) δ 171.46, 168.28, 166.46, 159.94, 156.15, 141.71, 139.40, 135.50, 130.69, 130.31, 129.22, 128.96, 128.77, 126.34, 124.69, 122.51, 119.85, 119.46, 117.86, 54.78, 33.70, 32.50.

**67 (mixture of diastereomers)**
$^1$H NMR (300 MHz, Acetone-*d*$_6$) δ 10.46 (d, *J* = 31.3 Hz, 1H), 9.95 (d, *J* = 53.8 Hz, 1H), 9.35 (d, *J* = 66.6 Hz, 1H), 8.12 – 7.99 (m, 3H), 7.81 (d, *J* = 7.8 Hz, 1H), 7.69 – 7.44 (m, 6H), 7.40 – 6.95 (m, 7H), 6.92 – 6.83 (m, 1H), 4.98 – 4.85 (m, 1H), 3.98 – 3.85 (m, 1H), 3.43 – 3.12 (m, 2H), 1.46 (t, *J* = 7.3 Hz, 3H).

**68**
$^1$H NMR (500 MHz, DMSO-*d*$_6$) δ 10.24 (s, 1H), 8.51 (d, *J* = 26.1, 8.0 Hz, 1H), 8.03 (t, *J* = 1.9 Hz, 1H), 7.93 – 7.89 (m, 1H), 7.86 – 7.80 (m, 2H), 7.77 (dt, *J* = 7.9, 2.2 Hz, 1H), 7.52 (dt, *J* = 7.8, 1.4 Hz, 1H), 7.45 – 7.39 (m, 2H), 7.38 – 7.33 (m, 1H), 7.32 – 7.15 (m, 8H), 7.09 – 7.04 (m, 4H), 7.02 – 6.94 (m, 3H), 6.50 – 6.43 (m, 2H), 4.73 – 4.39 (m, 1H), 4.19 (s, 2H), 3.01 – 2.81 (m, 2H).

**69**
$^1$H NMR (300 MHz, DMSO-*d*$_6$) δ 10.79 (s, 1H), 10.35 (s, 1H), 8.60 (d, *J* = 7.7 Hz, 1H), 8.11 – 8.05 (m, 1H), 7.97 – 7.71 (m, 5H), 7.54 (d, *J* = 7.9 Hz, 1H), 7.47 – 7.15 (m, 8H), 7.10 – 6.96 (m, 7H), 4.92 – 4.83 (m, 1H), 3.33 – 3.16 (m, 2H).

**70**

$^1$H NMR (300 MHz, DMSO-$d_6$) δ 10.32 (s, 1H), 8.65 (d, $J$ = 18.9, 8.1 Hz, 1H), 8.04 (t, $J$ = 1.9 Hz, 1H), 7.92 (s, 1H), 7.89 – 7.74 (m, 4H), 7.53 (d, $J$ = 7.7 Hz, 1H), 7.46 – 7.11 (m, 14H), 7.02 (dd, $J$ = 15.3, 8.2 Hz, 6H), 4.88 – 4.76 (m, 1H), 3.21 – 2.98 (m, 3H). $^{13}$C NMR (75 MHz, DMSO-$d_6$) δ 171.09, 168.26, 166.19, 159.95, 156.06, 139.34, 138.60, 135.52, 130.66, 130.14, 129.98, 129.63, 129.46, 129.01, 129.00, 128.61, 128.54, 126.76, 124.68, 122.53, 122.43, 119.85, 119.40, 117.83, 117.77, 56.35, 54.61, 37.53.

**71**

$^1$H NMR (300 MHz, DMSO-$d_6$) δ 10.23 (s, 1H), 9.94 (s, 1H), 8.59 (d, $J$ = 7.3 Hz, 1H), 8.08 (t, $J$ = 1.9 Hz, 1H), 8.03 – 7.90 (m, 3H), 7.85 – 7.74 (m, 1H), 7.56 (t, $J$ = 7.5 Hz, 3H), 7.50 – 7.16 (m, 8H), 7.13 – 7.00 (m, 6H), 4.67 – 4.53 (m, 1H), 2.30 – 2.04 (m, 2H).

**72**

$^1$H NMR (300 MHz, DMSO-$d_6$) δ 10.19 (s, 1H), 8.55 – 8.45 (m, 1H), 8.08 – 8.02 (m, 1H), 7.99 – 7.87 (m, 2H), 7.77 (dd, $J$ = 8.1, 2.2 Hz, 1H), 7.54 – 7.49 (m, 1H), 7.46 – 7.27 (m, 3H), 7.19 (t, $J$ = 7.4 Hz, 1H), 7.05 (td, $J$ = 9.0, 4.0 Hz, 4H), 4.59 – 4.27 (m, 1H), 1.87 – 1.70 (m, 2H), 1.49 – 1.19 (m, 4H), 0.92 – 0.81 (m, 3H). $^{13}$C NMR (75 MHz, DMSO-$d_6$) δ 174.38, 171.78, 168.32, 166.30, 159.92, 155.14, 139.47, 135.53, 130.68, 130.27, 130.14, 129.20, 128.97, 124.67, 122.38, 119.84, 119.81, 119.35, 117.89, 117.83, 54.92, 52.99, 31.71, 30.74, 28.47, 22.34, 22.17, 14.35.

**73**

$^1$H NMR (500 MHz, DMSO-$d_6$) δ 10.20 (s, 1H), 8.54 – 8.48 (m, 1H), 8.05 (t, $J$ = 1.9 Hz, 1H), 7.97 – 7.85 (m, 3H), 7.81 – 7.76 (m, 1H), 7.54 – 7.49 (m, 1H), 7.45 – 7.39 (m, 2H), 7.38 – 7.28 (m, 2H), 7.22 – 7.16 (m, 1H), 7.09 – 7.05 (m, 2H), 7.05 – 7.01 (m, 2H), 4.67 – 4.39 (m, 1H), 1.83 – 1.63 (m, 2H), 1.61 – 1.52 (m, 1H), 0.96 – 0.83 (m, 6H). $^{13}$C NMR (126 MHz, DMSO-$d_6$) δ 172.08, 168.32, 166.24, 166.14, 159.92, 159.90, 156.18, 139.48, 135.51, 130.68, 130.27, 130.18, 130.13, 129.82, 129.21, 128.96, 124.70, 124.67, 124.66, 122.44, 119.82, 119.79, 119.43, 117.92, 117.86, 114.87, 104.95, 98.83, 67.25, 55.79, 55.56, 53.24, 51.27, 40.69, 25.00, 24.98, 23.51, 23.41, 23.30, 21.89, 21.59.

**74**

$^1$H NMR (300 MHz, DMSO-$d_6$) δ 10.28 (s, 1H), 8.38 (d, $J$ = 20.0, 8.0 Hz, 1H), 8.12 – 8.07 (m, 1H), 8.00 – 7.88 (m, 3H), 7.78 (dd, $J$ = 8.1, 2.2 Hz, 1H), 7.53 (d, $J$ = 7.7 Hz, 1H), 7.48 – 7.29 (m, 3H), 7.20 (t, $J$ = 7.4 Hz, 1H), 7.11 – 6.98 (m, 4H), 4.35 (dt, $J$ = 44.3, 8.0 Hz, 1H), 1.97 – 1.81 (m, 2H), 1.77 – 1.56 (m, 4H), 1.29 – 0.98 (m, 5H). $^{13}$C NMR (75 MHz, DMSO-$d_6$) δ 170.89, 168.15, 166.15, 159.67, 155.99, 139.06, 135.40, 130.51, 130.13, 129.07, 128.82, 124.50, 122.34, 122.20, 119.65, 119.15, 117.67, 59.69, 39.40, 29.55, 26.10, 25.82.

**75**

$^1$H NMR (300 MHz, DMSO-$d_6$) δ 10.29 (s, 1H), 8.58 (d, $J$ = 17.5, 8.0 Hz, 1H), 8.05 (t, $J$ = 1.9 Hz, 1H), 7.98 – 7.74 (m, 5H), 7.60 – 7.49 (m, 1H), 7.48 – 7.28 (m, 5H), 7.24 – 7.13 (m, 3H), 7.12 – 6.97 (m, 6H), 6.70 – 6.57 (m, 3H), 4.80 – 4.69 (m, 1H), 3.10 – 2.86 (m, 2H). $^{13}$C NMR (75 MHz, DMSO-$d_6$) δ 173.82, 171.28, 168.30, 166.18, 159.94, 156.23, 139.39, 135.53, 130.67, 130.59, 130.40, 130.16, 130.01, 129.11, 129.01, 128.60, 124.69, 122.51, 122.44, 119.88, 119.39, 117.85, 117.79, 115.42, 115.35, 56.73.

**76**

$^1$H NMR (300 MHz, DMSO-$d_6$) δ 10.24 (s, 1H), 9.43 (s, 2H), 8.83 (d, $J$ = 13.0, 7.8 Hz, 2H), 8.08 (t, $J$ = 1.9 Hz, 1H), 8.01 – 7.83 (m, 6H), 7.81 – 7.75 (m, 2H), 7.72 – 7.30 (m, 17H), 7.26 – 7.17 (m, 2H), 7.12 – 7.00 (m, 7H), 4.96 (q, $J$ = 7.2 Hz, 1H), 3.42 – 3.17 (m, 3H). $^{13}$C NMR (75 MHz, DMSO-$d_6$) δ 169.75, 168.17, 166.39, 160.16, 156.03, 139.11, 135.45, 134.50, 130.71, 130.57, 130.30, 128.95, 128.88, 127.78, 124.78, 123.03, 121.98, 119.90, 117.82, 54.14, 27.74.

**77**

$^1$H NMR (300 MHz, DMSO-$d_6$) δ 10.19 (s, 1H), 8.56 – 8.43 (m, 1H), 8.08 – 8.03 (m, 1H), 7.99 – 7.87 (m, 3H), 7.82 – 7.73 (m, 1H), 7.52 (d, $J$ = 7.7 Hz, 1H), 7.47 – 7.27 (m, 4H), 7.24 – 7.14 (m, 1H), 7.13 – 6.99 (m, 5H), 4.55 (q, $J$ = 7.4 Hz, 1H), 1.77 (p, $J$ = 7.8 Hz, 2H), 1.54 – 1.27 (m, 2H), 0.97 – 0.83 (m, 3H). $^{13}$C NMR (75 MHz, DMSO-$d_6$) δ 171.78, 168.31, 166.30, 159.90, 156.16, 139.45, 135.52, 130.66, 130.25, 130.12, 129.20, 128.96, 122.42, 122.37, 119.82, 119.34, 117.82, 54.66, 34.00, 19.55, 14.08.

**78**

$^1$H NMR (500 MHz, DMSO-$d_6$) δ 10.16 (s, 1H), 8.80 (t, $J$ = 5.9 Hz, 1H), 8.06 – 8.02 (m, 1H), 7.96 – 7.86 (m, 5H), 7.81 – 7.75 (m, 1H), 7.56 – 7.51 (m, 1H), 7.48 – 7.41 (m, 4H), 7.37 (t, $J$ = 7.9 Hz, 1H), 7.32 (s, 1H), 7.25 – 7.18 (m, 2H), 7.13 – 6.98 (m, 8H), 4.06 (d, $J$ = 5.8 Hz, 2H).

**79**

$^1$H NMR (500 MHz, DMSO-$d_6$) δ 10.18 (s, 1H), 8.79 – 8.59 (m, 2H), 8.06 – 8.02 (m, 1H), 7.97 – 7.85 (m, 7H), 7.80 – 7.75 (m, 1H), 7.52 (dt, $J$ = 7.7, 1.4 Hz, 1H), 7.46 – 7.39 (m, 5H), 7.38 – 7.25 (m, 4H), 7.23 – 7.15 (m, 3H), 7.11 – 6.95 (m, 11H), 6.83 (d, $J$ = 17.9 Hz, 2H), 4.53 – 4.34 (m, 2H), 2.32 – 2.15 (m, 4H), 2.12 – 1.85 (m, 3H). $^{13}$C NMR (126 MHz, DMSO-$d_6$) δ 174.08, 171.25, 168.02, 165.41, 159.98, 156.45, 139.64, 135.81, 130.69, 130.18, 130.12, 130.07, 129.14, 128.96, 128.86, 124.72, 124.70, 122.45, 119.93, 119.90, 119.85, 119.81, 117.91, 117.85, 117.72, 110.00, 52.92, 52.29, 31.75, 27.48, 26.51.

### HPLC Purity of Inhibitors

Purity of inhibitors was determined by HPLC on a Jasco HPLC system with a Jasco UV-2070 Plus Intelligent UV/VIS Detector on an RP-18 column (ReproSil-Pur-ODS-3, Dr. Maisch GmbH, Germany, 5 µm, 50 mm × 2 mm) using the following method: eluent A: water (0.1% TFA); eluent B: acetonitrile (0.1% TFA); injection volume: 10 µL; flow rate: 1 mL/min; and gradient: 1% B (0.2 min), 100% B (7 min), 100% B (8 min), 1% B (8.1 min), and 1% B (10 min). Chromatograms recorded at 254 nm were used for purity assessment.

**Table S5**. RP-HPLC and MS data of compounds

| Cpd. | HRMS (ESI) for $[M+X]^+$ | | | HPLC $t_R$ [min] | HPLC purity (254 nm) |
|---|---|---|---|---|---|
| | Molecular Formula | Calc. | Found | | |
| **3** | $C_{35}H_{38}N_3O_4$ | 564.2857 | 564.2868 | 4.59 | > 95% |
| **4** | $C_{37}H_{34}N_3O_5$ | 600.2493 | 600.2492 | 4.33 | > 95% |
| **5** | $C_{33}H_{28}N_3O_5$ | 546.2023 | 546.2030 | 4.28 | > 95% |
| **6** | $C_{46}H_{45}N_5NaO_5$ | 784.3260 | 784.3253 | 4.68 | > 95% |
| **7** | $C_{35}H_{31}N_4O_4$ | 571.2340 | 571.2333 | 4.31 | > 95% |
| **8** | $C_{36}H_{32}N_3O_6$ | 602.2286 | 602.2265 | 4.23 | > 95% |
| **9** | $C_{36}H_{31}Cl_2N_4O_4$ | 653.1717 | 653.1690 | 4.60 | > 95% |
| **10** | $C_{36}H_{34}N_3O_5$ | 588.2493 | 588.2494 | 4.25 | > 95% |
| **11** | $C_{36}H_{32}N_3O_5$ | 586.2336 | 586.2301 | 4.22 | 90% |
| **12** | $C_{36}H_{43}N_4O_5$ | 611.3228 | 611.3233 | 3.96 | > 95% |
| **13** | $C_{37}H_{32}N_3O_5$ | 598.2336 | 598.2337 | 4.48 | 83% |
| **14** | $C_{37}H_{33}FN_3O_4$ | 602.2450 | 602.2437 | 4.37 | > 95% |
| **15** | $C_{33}H_{28}FN_4O_4S$ | 595.1810 | 595.1799 | 4.32 | > 95% |
| **16** | $C_{31}H_{38}N_3O_4$ | 516.2857 | 516.2857 | 4.31 | > 95% |
| **17** | $C_{32}H_{30}N_3O_4$ | 520.2231 | 520.2239 | 4.03 | > 95% |
| **18** | $C_{37}H_{33}N_4O_6$ | 629.2395 | 629.2383 | 3.95 | 91% |
| **19** | $C_{27}H_{23}ClN_3O_4S$ | 520.1092 | 520.1085 | 3.93 | > 95% |
| **20** | $C_{30}H_{36}N_3O_4$ | 502.2700 | 502.2708 | 4.16 | > 95% |
| **21** | $C_{34}H_{34}N_3O_4$ | 548.2544 | 548.2526 | 4.35 | > 95% |
| **22** | $C_{37}H_{32}ClN_4O_4$ | 631.2107 | 631.2100 | 4.43 | > 95% |
| **23** | $C_{38}H_{34}N_3O_5$ | 612.2493 | 612.2490 | 4.12 | > 95% |
| **24** | $C_{38}H_{41}N_4O_5$ | 633.3071 | 633.3094 | 4.02 | > 95% |
| **25** | $C_{30}H_{34}N_3O_4$ | 500.2544 | 500.2548 | 4.12 | 93% |
| **26** | $C_{32}H_{30}N_3O_5$ | 536.2180 | 536.2179 | 3.80 | 87% |
| **27** | $C_{32}H_{28}N_5O_5$ | 562.2085 | 562.2085 | 3.95 | 91% |
| **28** | $C_{31}H_{30}N_3O_4$ | 508.2231 | 508.2229 | 3.82 | > 95% |
| **29** | $C_{33}H_{28}FN_4O_5$ | 579.2038 | 579.2021 | 3.93 | > 95% |
| **30** | $C_{29}H_{32}N_3O_4$ | 486.2387 | 486.2389 | 3.91 | > 95% |
| **31** | $C_{33}H_{34}N_3O_7$ | 584.2391 | 584.2393 | 3.58 | > 95% |
| **32** | $C_{30}H_{34}N_3O_4$ | 500.2544 | 500.2529 | 3.95 | > 95% |
| **33** | $C_{34}H_{29}N_4O_5$ | 573.2132 | 573.2139 | 3.67 | > 95% |

**Table S5**. RP-HPLC and MS data of compounds

| | | | | | |
|---|---|---|---|---|---|
| **34** | $C_{34}H_{41}N_4O_5$ | 583.3071 | 583.3093 | 3.84 | > 95% |
| **35** | $C_{32}H_{28}N_5O_4S$ | 578.1857 | 578.1869 | 3.33 | > 95% |
| **36** | $C_{30}H_{28}N_3O_5$ | 510.2023 | 510.2001 | 3.96 | > 95% |
| **37** | $C_{29}H_{34}N_3O_4$ | 488.2544 | 488.2541 | 3.97 | > 95% |
| **38** | $C_{34}H_{33}N_4O_5$ | 577.2445 | 577.2435 | 3.89 | > 95% |
| **39** | $C_{33}H_{33}N_4O_5$ | 565.2445 | 565.2452 | 3.51 | > 95% |
| **40** | $C_{34}H_{34}N_4NaO_5S$ | 633.2142 | 633.2138 | 3.45 | > 95% |
| **41** | $C_{28}H_{30}N_3O_4$ | 472.2231 | 472.2232 | 3.86 | > 95% |
| **42** | $C_{29}H_{26}N_3O_4$ | 480.1923 | 480.1908 | 3.77 | > 95% |
| **43** | $C_{31}H_{30}N_3O_5$ | 524.2180 | 524.2173 | 3.42 | > 95% |
| **44** | $C_{29}H_{32}N_3O_4$ | 486.2387 | 486.2381 | 4.07 | > 95% |
| **45** | $C_{30}H_{28}N_3O_4$ | 494.2074 | 494.2054 | 3.69 | > 95% |
| **46** | $C_{29}H_{34}N_3O_4$ | 488.2544 | 488.2534 | 3.92 | > 95% |
| **47** | $C_{28}H_{28}N_3O_4$ | 470.2074 | 470.2080 | 3.71 | > 95% |
| **48** | $C_{27}H_{29}ClN_3O_4$ | 494.1841 | 494.1829 | 3.62 | > 95% |
| **49** | $C_{30}H_{28}N_3O_5$ | 510.2023 | 510.2021 | 3.36 | > 95% |
| **50** | $C_{28}H_{30}N_3O_4$ | 472.2231 | 472.2236 | 3.76 | > 95% |
| **51** | $C_{28}H_{32}N_3O_4$ | 474.2387 | 474.2387 | 3.87 | > 95% |
| **52** | $C_{28}H_{30}N_3O_4$ | 472.2231 | 472.2237 | 3.73 | > 95% |
| **53** | $C_{30}H_{33}N_4O_5$ | 529.2445 | 529.2454 | 3.28 | > 95% |
| **54** | $C_{35}H_{35}N_4O_5$ | 591.2602 | 591.2602 | 3.71 | > 95% |
| **55** | $C_{27}H_{28}N_3O_4$ | 458.2074 | 458.2074 | 3.59 | > 95% |
| **56** | $C_{32}H_{32}N_5O_5$ | 566.2398 | 566.2411 | 3.22 | > 95% |
| **57** | $C_{27}H_{31}N_4O_4S$ | 507.2061 | 507.2050 | 3.07 | > 95% |
| **58** | $C_{31}H_{37}N_4O_4$ | 529.2809 | 529.2817 | 3.39 | 93% |
| **59** | $C_{29}H_{27}N_4O_6S$ | 559.1646 | 559.1634 | 3.32 | > 95% |
| **60** | $C_{28}H_{27}N_3NaO_4$ | 492.1894 | 492.1871 | 3.47 | > 95% |
| **61** | $C_{26}H_{27}ClN_3O_4$ | 480.1685 | 480.1669 | 3.49 | > 95% |
| **62** | $C_{36}H_{32}N_3O_5$ | 586.2336 | 586.2311 | 4.31 | > 95% |
| **63** | $C_{40}H_{34}N_3O_5$ | 636.2493 | 636.2490 | 4.17 | > 95% |
| **64** | $C_{29}H_{32}N_3O_4$ | 486.2387 | 486.2388 | 4.14 | > 95% |
| **65** | $C_{33}H_{28}N_3O_4$ | 530.2074 | 530.2068 | 3.71 | > 95% |
| **66** | $C_{30}H_{28}N_3O_4$ | 494.2074 | 494.2079 | 3.95 | > 95% |

**Table S5**. RP-HPLC and MS data of compounds

| | | | | | |
|---|---|---|---|---|---|
| **67** | $C_{33}H_{28}ClN_5NaO_3$ | 600.1773 | 600.1760 | 3.99 | > 95% |
| **68** | $C_{36}H_{32}N_4O_4$ | 585.2496 | 585.2484 | 3.49 | 77% |
| **69** | $C_{31}H_{27}N_4O_4$ | 519.2027 | 519.2010 | 3.76 | > 95% |
| **70** | $C_{29}H_{26}N_3O_4$ | 480.1918 | 480.1915 | 3.79 | 90% |
| **71** | $C_{31}H_{28}N_4O_5$ | 537.2132 | 537.2138 | 3.67 | 91% |
| **72** | $C_{26}H_{28}N_3O_4$ | 446.2074 | 446.2073 | 3.77 | > 95% |
| **73** | $C_{26}H_{27}N_3NaO_4$ | 468.1894 | 468.1904 | 3.71 | > 95% |
| **74** | $C_{28}H_{30}N_3O_4$ | 472.2231 | 472.2225 | 3.92 | > 95% |
| **75** | $C_{29}H_{26}N_3O_5$ | 496.1867 | 496.1845 | 3.42 | 87% |
| **76** | $C_{32}H_{28}N_5O_4$ | 546.2136 | 546.2131 | 3.32 | 92% |
| **77** | $C_{25}H_{26}N_3O_4$ | 432.1918 | 432.1919 | 3.50 | 78% |
| **78** | $C_{22}H_{19}N_3NaO_4$ | 412.1259 | 412.1268 | 3.06 | > 95% |
| **79** | $C_{25}H_{25}N_4O_5$ | 461.1819 | 461.1814 | 3.43 | 83% |

**Off-Target Testing Supplementary**

**Thrombin Assay**

Thrombin was purchased from Sigma-Aldrich (Germany). The thrombin assay was performed as reported.[2] Continuous fluorimetric assay was done in black 96 well V-bottom plates (Greiner Bio-One, Germany), using a BMG Labtech Fluostar OPTIMA microtiter fluorescence plate reader. Excitation wavelength of 355 nm and an emission wavelength of 460 nm were used. The inhibitors (final concentration 25 µM, from 10 mM stock solutions in DMSO) were preincubated with thrombin (10 nM) in the assay buffer (50 mM Tris-HCl pH 7.5, 150 mM NaCl, 0.05% Tween 20) for 15 min. Enzymatic cleavage was initiated by the addition of the Boc-Val-Pro-Arg-AMC substrate (Bachem, Germany) at a final concentration of 50 µM. The activity of thrombin was monitored for 15 min and determined as a slope of relative fluorescence units per second (RFU/s). Camostat mesylate was used as inhibition control. All experiments were performed in triplicates and percentage inhibition was calculated as the mean and respective standard deviation of the values. Values were obtained in relation to a positive control.

**Trypsin Assay**

Trypsin was purchased from Sigma-Aldrich (Germany). The inhibition of trypsin was determined as described before.[12] Continuous fluorimetric assay was done in black 96 well V-bottom plates (Greiner Bio-One, Germany), using a BMG Labtech Fluostar OPTIMA microtiter fluorescence plate reader. Excitation wavelength of 355 nm and an emission wavelength of 460 nm were used. The inhibitors (final concentration 50 µM, from 10 mM stock solutions in DMSO) were preincubated with trypsin (1 nM) in the assay buffer (50 mM Tris-HCl pH 7.5, 150 mM NaCl, 0.05% Tween 20) for 15 min. Enzymatic cleavage was initiated by the addition of the Boc-Val-Pro-Arg-AMC substrate (Bachem, Germany) at a final concentration of 50 µM. The activity of thrombin was monitored for 15 min and determined as a slope of relative fluorescence units per second (RFU/s). Camostat mesylate was used as positive control. All experiments were performed in triplicates and percentage inhibition was calculated as the mean and respective standard deviation of the values. Values were obtained in relation to a positive control (without inhibitor).

**Table S6**. Inhibitory activity of compounds against thrombin and trypsin.

| Cpd. | Thrombin[a] (%) | Trypsin[b] (%) |
|---|---|---|
| **3** | 12 | n.i. |
| **4** | n.i. | n.i. |
| **5** | n.i. | n.i. |
| **6** | n.i. | n.i. |
| **7** | 14 | n.i. |
| **8** | 13 | 36 |
| **9** | n.i. | n.i. |
| **10** | n.i. | n.i. |
| **11** | n.i. | n.i. |
| **12** | n.i. | n.i. |
| **13** | n.i. | n.i. |
| **14** | n.i. | n.i. |
| **15** | 14 | n.i. |
| **16** | n.i. | n.i. |
| **17** | n.i. | n.i. |
| **18** | n.i. | n.i. |
| **19** | n.i. | n.i. |
| **20** | n.i. | n.i. |
| **21** | n.i. | n.i. |
| **22** | n.i. | n.i. |
| **23** | n.i. | n.i. |
| **24** | n.i. | n.i. |
| **25** | n.i. | n.i. |
| **26** | n.i. | n.i. |
| **27** | 17 | 12 |
| **28** | n.i. | n.i. |
| **29** | n.i. | n.i. |
| **30** | n.i. | n.i. |
| **31** | n.i. | n.i. |
| **32** | n.i. | n.i. |
| **33** | n.i. | n.i. |
| **34** | n.i. | n.i. |
| **35** | n.i. | n.i. |
| **36** | n.i. | n.i. |
| **37** | n.i. | n.i. |
| **38** | n.i. | n.i. |
| **39** | n.i. | n.i. |
| **40** | n.i. | n.i. |
| **41** | n.i. | n.i. |
| **42** | n.i. | n.i. |
| **43** | n.i. | n.i. |
| **44** | n.i. | n.i. |
| **45** | n.i. | n.i. |

**Table S6**. Inhibitory activity of compounds against thrombin and trypsin.

| | | |
|---|---|---|
| **46** | 14 | n.i. |
| **47** | n.i. | n.i. |
| **48** | n.i. | n.i. |
| **49** | n.i. | n.i. |
| **50** | n.i. | n.i. |
| **51** | n.i. | n.i. |
| **52** | n.i. | n.i. |
| **53** | n.i. | n.i. |
| **54** | n.i. | n.i. |
| **55** | n.i. | n.i. |
| **56** | n.i. | n.i. |
| **57** | n.i. | n.i. |
| **58** | n.i. | n.i. |
| **59** | n.i. | n.i. |
| **60** | n.i. | n.i. |
| **61** | n.i. | n.i. |
| **62** | n.i. | n.i. |
| **63** | n.i. | n.i. |
| **64** | n.i. | n.i. |
| **65** | n.i. | n.i. |
| **66** | n.i. | 11 |
| **67** | n.i. | n.i. |
| **68** | 14 | n.i. |
| **69** | n.i. | 11 |
| **70** | n.i. | n.i. |
| **71** | n.i. | n.i. |
| **72** | n.i. | n.i. |
| **73** | n.i. | 23 |
| **74** | n.i. | n.i. |
| **75** | n.i. | n.i. |
| **76** | n.i. | n.i. |
| **77** | n.i. | n.i. |
| **78** | n.i. | n.i. |
| **79** | n.i. | n.i. |
| **Camostat** | 96 (99[6]) | 100 (101[6]) |

[a]Inhibition of thrombin (inhibitor 25 µM, substrate 50 µM). [b]Inhibition of trypsin (inhibitor 50 µM, substrate 50 µM). If inhibition ≤ 10% = no inhibition (n.i.). All measurements were carried out in triplicate.

### References

(1) Nitsche, C.; Klein, C. D. Fluorimetric and HPLC-Based Dengue Virus Protease Assays Using a FRET Substrate. In *Antiviral Methods and Protocols*, Gong, E. Y. Ed.; Humana Press, 2013; pp 221-236.
(2) Nitsche, C.; Schreier, V. N.; Behnam, M. A. M.; Kumar, A.; Bartenschlager, R.; Klein, C. D. Thiazolidinone–Peptide Hybrids as Dengue Virus Protease Inhibitors with Antiviral Activity in Cell Culture. *Journal of Medicinal Chemistry* **2013**, *56* (21), 8389-8403. DOI: 10.1021/jm400828u.
(3) Behnam, M. A. M.; Nitsche, C.; Vechi, S. M.; Klein, C. D. C-Terminal Residue Optimization and Fragment Merging: Discovery of a Potent Peptide-Hybrid Inhibitor of Dengue Protease. *ACS Medicinal Chemistry Letters* **2014**, *5* (9), 1037-1042. DOI: 10.1021/ml500245v.
(4) Behnam, M. A. M.; Graf, D.; Bartenschlager, R.; Zlotos, D. P.; Klein, C. D. Discovery of Nanomolar Dengue and West Nile Virus Protease Inhibitors Containing a 4-Benzyloxyphenylglycine Residue. *Journal of Medicinal Chemistry* **2015**, *58* (23), 9354-9370. DOI: 10.1021/acs.jmedchem.5b01441.
(5) Wulff, N. H.; Tzatzaris, M.; Young, P. J. Monte Carlo simulation of the Spearman-Kaerber TCID50. *Journal of Clinical Bioinformatics* **2012**, *2* (1), 5. DOI: 10.1186/2043-9113-2-5.
(6) Kühl, N.; Leuthold, M. M.; Behnam, M. A. M.; Klein, C. D. Beyond Basicity: Discovery of Nonbasic DENV-2 Protease Inhibitors with Potent Activity in Cell Culture. *Journal of Medicinal Chemistry* **2021**, *64* (8), 4567-4587. DOI: 10.1021/acs.jmedchem.0c02042.
(7) Kühl, N.; Graf, D.; Bock, J.; Behnam, M. A. M.; Leuthold, M.-M.; Klein, C. D. A New Class of Dengue and West Nile Virus Protease Inhibitors with Submicromolar Activity in Reporter Gene DENV-2 Protease and Viral Replication Assays. *Journal of Medicinal Chemistry* **2020**, *63* (15), 8179-8197. DOI: 10.1021/acs.jmedchem.0c00413.
(8) Behnam, M. A. M.; Klein, C. D. Alternate recognition by dengue protease: Proteolytic and binding assays provide functional evidence beyond an induced-fit. *bioRxiv* **2024**, 2024.2004.2015.589505. DOI: 10.1101/2024.04.15.589505.
(9) Steuer, C.; Gege, C.; Fischl, W.; Heinonen, K. H.; Bartenschlager, R.; Klein, C. D. Synthesis and biological evaluation of α-ketoamides as inhibitors of the Dengue virus protease with antiviral activity in cell-culture. *Bioorganic & Medicinal Chemistry* **2011**, *19* (13), 4067-4074. DOI: https://doi.org/10.1016/j.bmc.2011.05.015.
(10) Nitsche, C.; Behnam, M. A. M.; Steuer, C.; Klein, C. D. Retro peptide-hybrids as selective inhibitors of the Dengue virus NS2B-NS3 protease. *Antiviral Research* **2012**, *94* (1), 72-79. DOI: https://doi.org/10.1016/j.antiviral.2012.02.008.
(11) Amblard, M.; Fehrentz, J.-A.; Martinez, J.; Subra, G. J. M. b. Methods and protocols of modern solid phase peptide synthesis. **2006**, *33*, 239-254.
(12) Weigel, L. F.; Nitsche, C.; Graf, D.; Bartenschlager, R.; Klein, C. D. Phenylalanine and Phenylglycine Analogues as Arginine Mimetics in Dengue Protease Inhibitors. *Journal of Medicinal Chemistry* **2015**, *58* (19), 7719-7733. DOI: 10.1021/acs.jmedchem.5b00612.
